# Mechanistic theory predicts the effects of temperature and humidity on inactivation of SARS-CoV-2 and other enveloped viruses

**DOI:** 10.1101/2020.10.16.341883

**Authors:** Dylan H. Morris, Kwe Claude Yinda, Amandine Gamble, Fernando W. Rossine, Qishen Huang, Trenton Bushmaker, Robert J. Fischer, M. Jeremiah Matson, Neeltje van Doremalen, Peter J. Vikesland, Linsey C. Marr, Vincent J. Munster, James O. Lloyd-Smith

## Abstract

Environmental conditions affect virus inactivation rate and transmission potential. Understanding those effects is critical for anticipating and mitigating epidemic spread. Ambient temperature and humidity strongly affect the inactivation rate of enveloped viruses, but a mechanistic, quantitative theory of those effects has been elusive. We measure the stability of the enveloped respiratory virus SARS-CoV-2 on an inert surface at nine temperature and humidity conditions and develop a mechanistic model to explain and predict how temperature and humidity alter virus inactivation. We find SARS-CoV-2 survives longest at low temperatures and extreme relative humidities; median estimated virus half-life is over 24 hours at 10 °C and 40 % RH, but approximately 1.5 hours at 27 °C and 65 % RH. Our mechanistic model uses simple chemistry to explain the increase in virus inactivation rate with increased temperature and the U-shaped dependence of inactivation rate on relative humidity. The model accurately predicts quantitative measurements from existing studies of five different human coronaviruses (including SARS-CoV-2), suggesting that shared mechanisms may determine environmental stability for many enveloped viruses. Our results indicate scenarios of particular transmission risk, point to pandemic mitigation strategies, and open new frontiers in the mechanistic study of virus transmission.

## Introduction

For viruses to transmit from one host to the next, virus particles must remain infectious in the period between release from the transmitting host and uptake by the recipient host. Virus environmental stability thus determines the potential for surface (fomite) transmission and for mid-to-long range transmission through the air. Empirical evidence suggests that virus environmental stability depends strongly on ambient temperature and humidity, particularly for enveloped viruses; examples among enveloped viruses that infect humans include influenza viruses [40], endemic human coronaviruses [25], and the zoonotic coronaviruses SARS-CoV-1 [11] and MERS-CoV [59].

In late 2019, a new zoonotic coronavirus now called SARS-CoV-2 emerged; it has since caused a global pandemic (COVID-19), and is poised to become an endemic human pathogen. As the northern hemisphere enters winter, many countries in the temperate north have seen an increase in transmission. Epidemiologists anticipated that increase [46, 33] based on observations from other enveloped respiratory viruses, such as endemic human coronaviruses [45] and influenza viruses [39], which spread more readily in temperate winters than in temperate summers. Like the related SARS-CoV-1 virus [38], SARS-CoV-2 displays epidemic dynamics that are strongly shaped by superspreading events, in which one person transmits to many others [20, 29].

Virus transmission is governed by many factors, among them properties of the virus and properties of the host population. But anticipating seasonal changes in transmission and preventing superspreading events both require an understanding of virus persistence in the environment, as ambient conditions can facilitate or impede virus spread.

Empirical evidence suggests that SARS-CoV-2, like other enveloped viruses, varies in its environmental stability as a function of temperature and humidity [6, 42], but the joint effect of these two factors remains unclear.

Moreover, despite years of research on virus environmental stability, there do not exist mechanistically motivated quantitative models for virus inactivation as a function of both temperature and humidity. This makes it difficult to generalize from any given experiment to unobserved conditions, or to real-world settings. Existing predictive models for the environmental stability of SARS-CoV-2 [6, 22] and other viruses [50] are phenomenological regression models that do not model the underlying biochemical mechanisms of inactivation. This limits both our insight into the underlying inactivation process and our ability to extrapolate reliably. A lack of quantitative, mechanistic models also makes it difficult to determine which environmental factors are most important, for instance whether absolute humidity [54] or relative humidity [40] best explains influenza inactivation and seasonality.

We measured the environmental stability of SARS-CoV-2 virus particles (virions) suspended in cell culture medium and deposited onto a polypropylene plastic surface at nine environmental conditions: three relative humidities (RH; 40 %, 65 %, and 85 %) at each of three temperatures (10 °C, 22 °C, and 27 °C). We quantified viable (infectious) virus titer over time and estimated virus decay rates and corresponding half-lives in each condition using a simple Bayesian regression model (see Methods). We quantified the evaporation of the suspension medium and compared virus stability during the sample evaporation phase—while substantial water loss was ongoing—to virus stability after a quasi-equilibrium phase was reached—when further evaporation was not evident over the timescale of the experiment.

We then created a mechanistic biochemical model of virus inactivation kinetics, drawing upon existing hypotheses for how temperature and humidity affect the inactivation chemistry of virus particles in microdroplets [40, 37]. We fit this mechanistic model to our SARS-CoV-2 data, and used it to predict observations from other human coronaviruses and other studies of SARS-CoV-2, in addition to unobserved temperature and humidity conditions.

### Empirical patterns of virus decay

Our data suggest that SARS-CoV-2 environmental persistence could vary meaningfully across the range of temperatures and humidities encountered in daily life, with posterior median [95 % credible interval] half-lives as long as 27 h [20, 39] (10 °C, 40 % RH) and as short as 1.5h [1.1, 2.1] (27 °C, 65 % RH), once droplets reach quasi-equilibrium with the ambient air conditions (Fig. 1b, Appendix Table A1).

**Figure 1.**
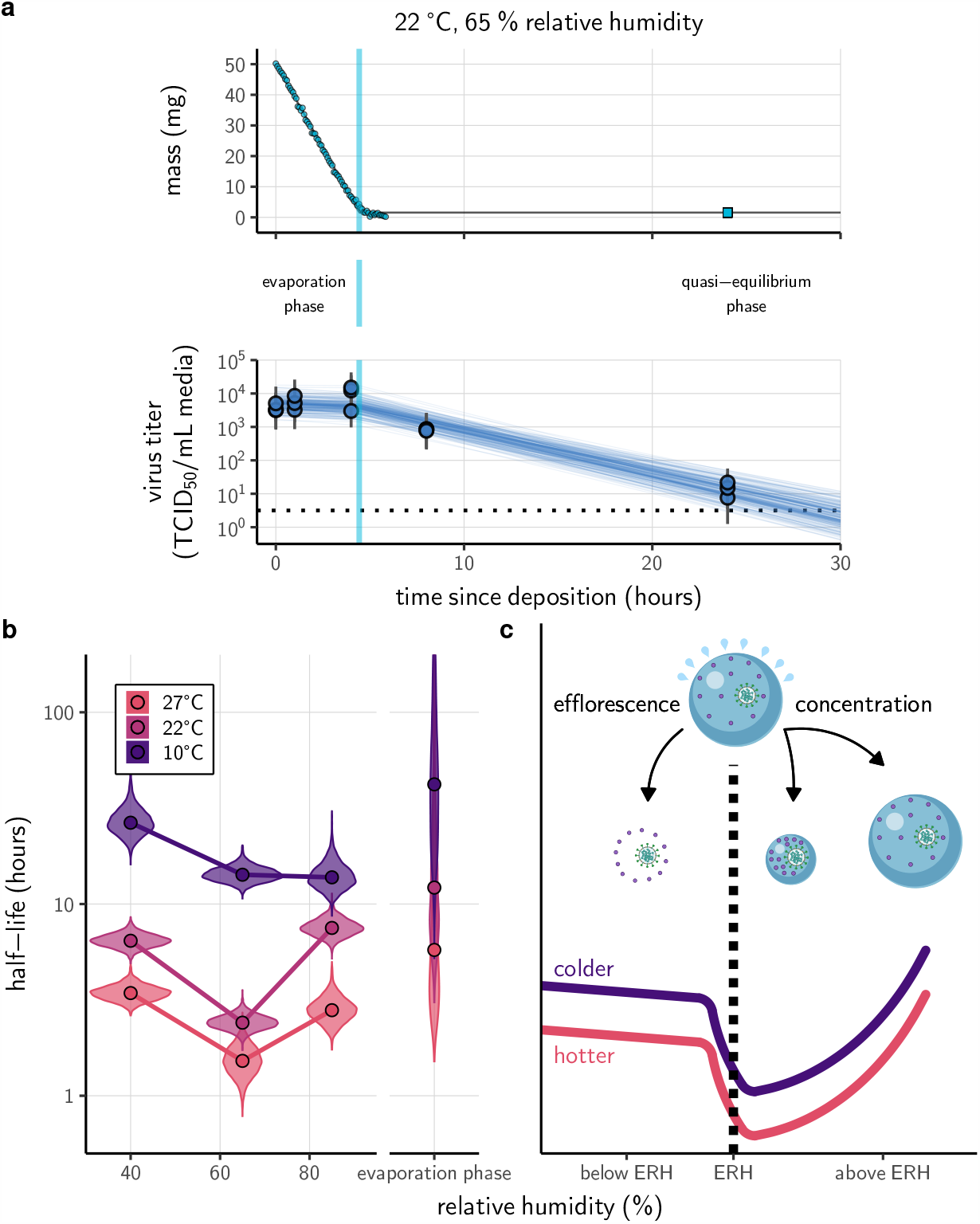
Inactivation kinetics and estimated half-life of SARS-CoV-2 on an inert surface as a function of temperature and relative humidity (RH). (**a**) Example of medium evaporation and virus inactivation as a function of time since deposition; experiments at 22 °C and 65 % RH shown. Inactivation proceeds in two phases: an evaporation phase during which water mass is lost from the sample to evaporation and a quasi-equilibrium phase once the sample mass has plateaued. Light blue vertical line shows posterior median estimated time that quasi-equilibrium was reached. Top plot: medium evaporation. Dots show measured masses. Square shows measured final (quasi-equilibrium) mass; plotted at 24 h for readability. Lines are 10 random draws from the posterior for the evaporation rate; horizontal section of line reflects the reaching of quasi-equilibrium (measured final mass). Bottom plot: virus inactivation. Points show posterior median estimated titers in log_10_TCID_50_/mL for each sample; lines show 95 % credible intervals. Black dotted line shows the approximate single-replicate limit of detection (LOD) of the assay: 10^0.5^ TCID_50_/mL media. Three samples collected at each time-point. Lines are 10 random draws per measurement from the posterior distribution for the inactivation rates estimated by the simple regression model (see Methods). (**c**) Measured virus half-lives. Violin plots show posterior distribution of estimated half-lives, plotted on a logarithmic scale. Dots show posterior median value. Color indicates temperature. Measurements at 40 %, 65 %, and 85 % RH reflect decay kinetics once the deposited solution has reached quasi-equilibrium with the ambient air. Estimated half-lives for the evaporation phase that occurs prior to quasi-equilibrium are plotted to the right, since conditions during this phase are mainly dilute, and thus analogous to high RH quasi-equilibrium conditions. (**b**) Schematic of hypothesized effects of temperature and relative humidity on duration of virus viability. Virus half-lives are longer at lower temperatures, regardless of humidity, because inactivation reaction kinetics proceed more slowly. Relative humidity affects virus half-life by determining quasi-equilibrium solute concentration in the droplet containing the virus. Above the efflorescence relative humidity (ERH), solutes are concentrated by evaporation. The lower the ambient humidity, the more water evaporates, the more concentration occurs, and the faster inactivation reactions proceed. Below the ERH, solutes effloresce, forming crystals. Half-lives are thus not particularly sensitive to changes in sub-ERH relative humidity, and half-lives even slightly below the ERH may be substantially longer than half-lives slightly above it.

Minimal virus decay occurred during the evaporation phase (Fig. 1a, Appendix Fig. A2), when excess water was present. Estimated half-lives were long but exact values were highly uncertain, as the small amount of absolute virus inactivation during the brief evaporation phases, combined with the noise involved in sampling and titration, limits our inferential capacity. Posterior median evaporation phase half-lives were 42 h [11, 330] at 10 °C, 12 h [4.5, 160] at 22 °C, and 5.8h [2.1, 130] at 27 °C (Appendix Table A1).

Overall, virus decay became markedly faster as temperature increased for all humidities, with decay at 27 °C roughly five to ten times faster than decay at 10 °C. Across temperatures, virus decay was relatively rapid at 65 % RH and tended to be slower either at lower (40 %) or higher (85 %) humidities or when excess water was present during the evaporation phase (Fig. 1b).

### Mechanistic model for temperature and humidity effects

Many viruses, including SARS-CoV-2, exhibit exponential decay on surfaces and in aerosols [40, 15, 6]. We drew upon known principles of droplet chemistry and its potential effects on virus inactivation chemistry (Fig. 1c) to create a minimal mechanistic model incorporating the effects of both temperature and relative humidity on exponential decay rates.

We model temperature dependence with the Arrhenius equation, which describes a reaction rate *k* as a function of an activation energy *E*_*a*_, an asymptotic high-temperature reaction rate *A*, the universal gas constant *R*, and the absolute temperature *T*:

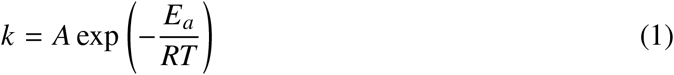

Prior work has found Arrhenius-like temperature dependence for virus inactivation on surfaces and in aerosols for many viruses [1], including human coronaviruses [65].

Mechanistic principles of virus inactivation as a function of humidity have been more elusive, but recent work has suggested that relative humidity affects virus inactivation by controlling evaporation and thus governing the solute concentrations in a droplet containing virions [40, 37]. In more humid environments, evaporation is slower and more water remains when quasi-equilibrium is reached. In less humid environments, evaporation is faster and little or no water remains (Fig. 1c).

When released from infected hosts, virions are found in host bodily fluids, whereas viral inactivation experiments are typically conducted in cell culture medium containing amino-acids and electrolytes, in particular sodium chloride (NaCl) [10, 16]. Prior work has found that higher quasi-equilibrium solute concentrations are associated with faster virus inactivation rates [64, 63]. The simplest explanation for this is that the measured solute concentration is a direct proxy for the concentration of the reactants governing the inactivation reaction. Thus ambient humidity affects the reaction rate by setting the quasi-equilibrium concentrations of the reactants that induce inactivation of the virus.

The exact quasi-equilibrium state reached will depend on the solutes present, since different solutes depress vapor pressure to different degrees. In electrolyte solutions like bodily fluids or cell culture media, efflorescence is also important. Below a threshold ambient humidity—the efflorescence relative humidity (ERH)—electrolytes effloresce out of solution, forming a crystal (Fig. 1c). Below the ERH, the reaction no longer occurs in solution, and so inactivation may be slower. The notable U-shape of virus inactivation as a function of relative humidity, observed in our data (Fig. 1a) and elsewhere in the literature [63, 5, 52, 60], including for coronaviruses [9, 55], could be explained by this regime shift around the ERH (Fig. 1c).

To quantify these effects, we model virus inactivation at quasi-equilibrium on inert surfaces as a chemical reaction with first-order reaction kinetics; that is, the quantity of virus is the limiting reactant of the rate-determining step, and the concentrations of other reactants are assumed to be approximately constant over time. At constant temperature and humidity, the quantity of virus should then exhibit exponential decay. During the evaporation phase prior to quasi-equilibrium, reactants are less concentrated and decay is expected to be slower, as observed from our data (Fig. 1a,b). If small initial droplet sizes are used—as in real-world depositions (predominantly *<* 10 µL [28, 27, 58]) and in some experiments—evaporative quasi-equilibration should be near instant, and so inactivation should follow the kinetics at quasi-equilibrium. Larger droplets, such as those used in our experiments, will take more time to equilibrate (depending on temperature and humidity), allowing us to distinguish the quasi-equilibrium phase from the evaporation phase.

We partition inactivation at quasi-equilibrium into two humidity regimes, effloresced and solution, according to whether the ambient RH is below the ERH (effloresced) or above (solution). In either case, we approximate virus inactivation as a first-order reaction with rate *k*_eff_ or *k*_sol_, respectively. Based on observations of NaCl solutions at room temperature and atmospheric pressure [44], we use an ERH of 45 %. This means that 40 % RH experiments are in the effloresced regime and 65 % and 85 % RH experiments are in the solution regime.

We model the effloresced and solution inactivation rates *k*_eff_ and *k*_sol_ using two Arrhenius equations with a shared activation energy *E*_*0*_ but distinct asymptotic high-temperature reaction rates *A*_eff_ and *A*_sol_. In solution conditions, we further modulate *k*_sol_ by a quasi-equilibrium “concentration factor 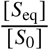: how concentrated the solution has become at quasi-equilibrium [*S*_eq_] relative to its initial state [*S*_0_]. Given our assumption of first-order kinetics, an n-fold increase in the non-virion reactant concentrations should translate directly into an n-fold increase in the inactivation rate.

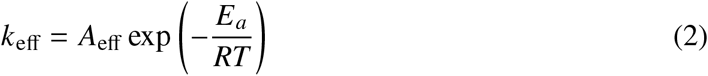

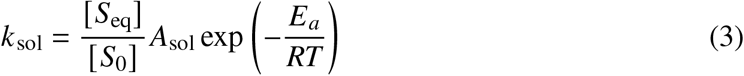

We estimated *E*_*a*_, *A*_eff_, and *A*_sol_ from our data, constraining all to be positive. We treated evaporation phase data as governed by :_sol_, with a dynamic value of the concentration factor 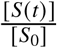 (Appendix section 4.4). We computed the quasi-equilibrium concentration factor 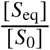 in two ways: using measurements from our evaporation experiments (measured concentration fit) and with a theoretically motivated curve fit to our virological data (modeled concentration fit, Appendix Fig. A9). See Appendix section 5.5.3 for details.

We also considered a 4-parameter variant of the model with distinct activation energies below the ERH 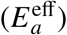 and above 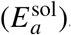, placing the same prior on each. This accounts for the possibility that the rate-determining step of the inactivation reaction might be distinct in the two regimes. The estimated activation energies were near-identical below and above the ERH (Fig. A8): the posterior median percentage difference between two *E*_*a*_ values was less than 2% (−1.2 %, 95 % cred. int. [−24 %, 17 %]) for the measured concentration fit. This suggests that the rate-determining reaction step—and thus the activation energy—is the same in both regimes. Accordingly, we report estimates from the 3-parameter model with a shared *E*_*a*_. We provide additional details and interpretation of our mechanistic inactivation modeling in the Appendix, sections 3, 5.5.

### Model fitting and prediction of unobserved conditions

Our dataset comprises 9 experimental conditions, each with 7 time-points that span the evaporation and quasi-equilibrium phases. We sought to explain the virus inactivation rates across this entire dataset using our mechanistic model with just 3 free parameters: the activation energy *E*_*a*_ and the asymptotic high-temperature reaction rates under effloresced and solution conditions, *A*_eff_ and *A*_sol_. The mechanistic function used and the constraint on the parameters to be positive means that inactivation rate must increase with temperature and with increasing solute concentration. Remarkably, the fit of the mechanistic model (Fig. 2, Appendix Figs. A3, A4, A6) is virtually indistinguishable from the fit of the simple regression, which estimates independent exponential decay rates for each condition (Appendix Figs. A2, A5, see Appendix section 5.4.1). Parameter estimates are given in the Appendix (Fig. A10, Tables A3, A4).

**Figure 2.**
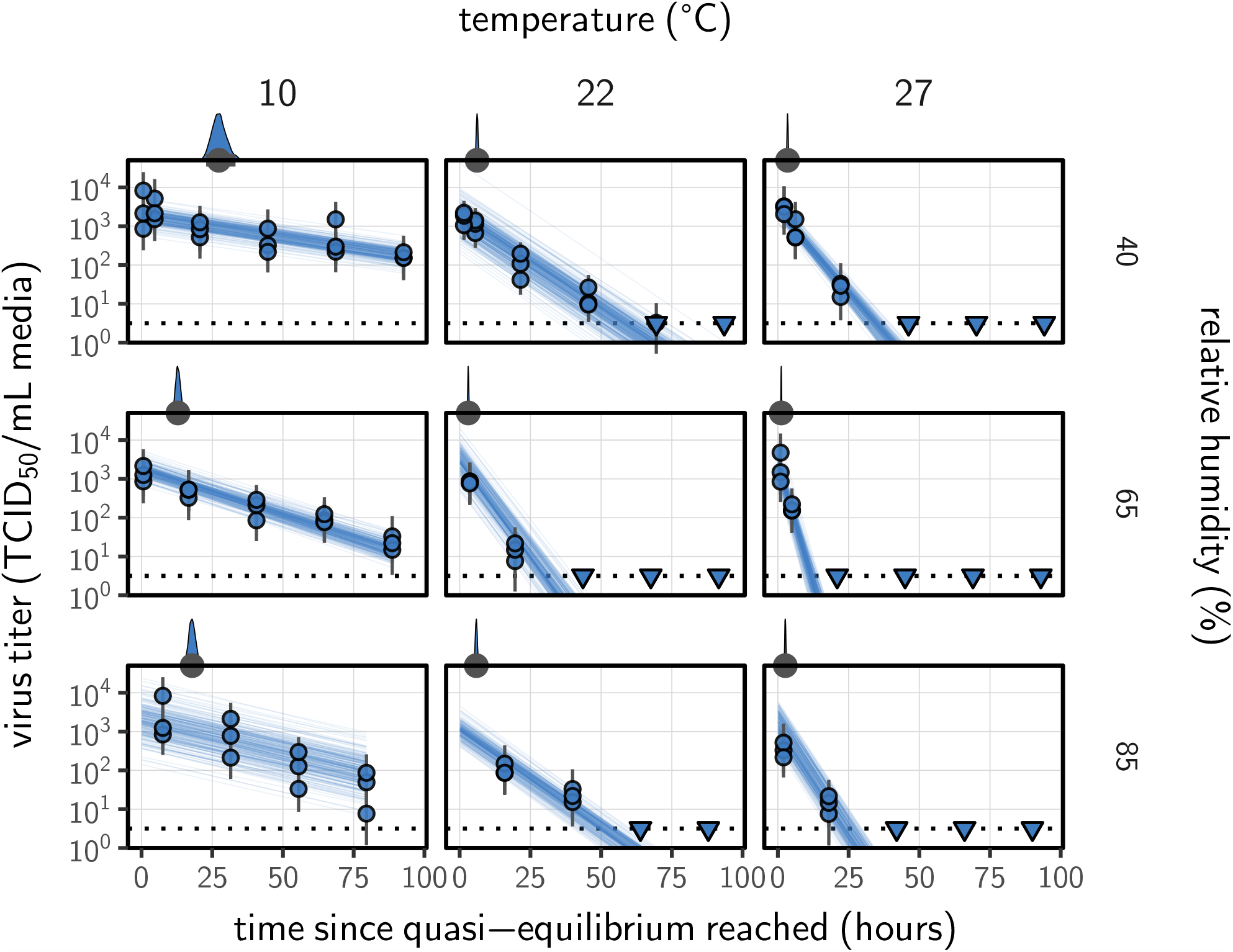
Estimated titers and mechanistic model fit for SARS-CoV-2 stability on polypropylene at quasi-equilibrium. Points show posterior median estimated titers in log_10_TCID_50_/mL for each sample; lines show 95 % credible intervals. Time-points with no positive wells for any replicate are plotted as triangles at the approximate single-replicate limit of detection (LOD) of the assay—denoted by a black dotted line at 10^0.5^ TCID_50_/mL media—to indicate that a range of sub-LOD values are plausible. Three samples collected at each time-point. x-axis shows time since quasi-equilibrium was reached, as measured in evaporation experiments. Lines are random draws (10 per sample) from the joint posterior distribution of the initial sample virus concentration and the mechanistic model predicted decay rate; the distribution of lines gives an estimate of the uncertainty in the decay rate and the variability of the initial titer for each experiment. Density plots above each box show posterior distribution of virus half-life according to the model for the given condition; point under the density shows the posterior median half-life and line shows a 95 % credible interval. Parameters from the measured concentration model fit.

We used the mechanistic model to predict SARS-CoV-2 half-life for unobserved temperature and humidity conditions from 0 to 40 °C, and from 0 to 100 % RH. We chose these ranges to reflect environments encountered by human beings in daily life. We did not extrapolate to temperatures below 0 °C since inactivation kinetics may be different when fluid containing the virus freezes. The exact freezing points of suspension medium and human fluids at sea level will depend on solute concentration, but will typically be below the 0 °C freezing point of pure water.

Median predicted SARS-CoV-2 half-life varies by more than three orders of magnitude, from less than half an hour at 40 °C just above the modeled approximate ERH, to more than a month at 0 °C and 100 % RH (Fig. 3a and c). We find good qualitative agreement between model predictions and model-free estimates from our data, including long half-lives prior to quasi-equilibrium. The U-shaped effect of humidity on virus half-life is readily explained by the regime-shift at the ERH (Fig. 3a). In particular, half-lives become extremely long at cold temperatures and in very dilute solutions, which are expected at high RH (Fig. 3a,b). Of note, the worst agreement between predictions and model-free estimates is found at 10 °C and 85 % RH (Fig. 3b). This is partially explained by the fact that the quasi-equilibrium concentration reached under those conditions was higher than our model prediction of concentration from RH (Appendix Fig. A9). Accordingly, the half-life prediction for 10 °C and 85 % RH based on measured concentration (Fig. 3b) is superior to the prediction based on modeled concentration (Fig. 3a).

**Figure 3.**
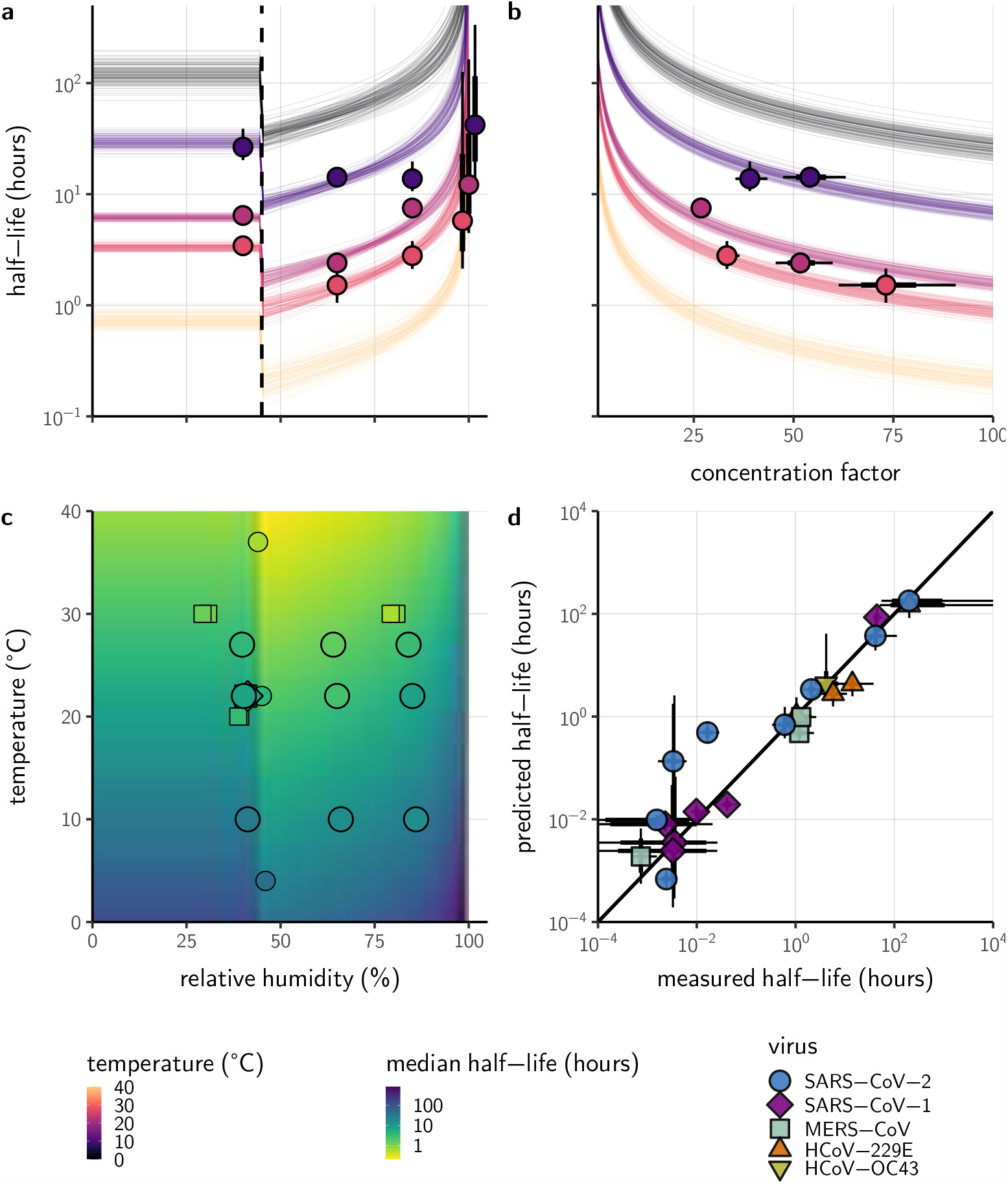
Extrapolation of human coronavirus half-life from the mechanistic model to unobserved temperatures and humidities and prediction of data from the literature. (**a**) Predicted half-life as a function of relative humidity, according to the modeled concentration fit. Points show posterior median for measured half-lives, estimated without the mechanistic model (i.e. independent estimation of a fixed exponential decay rate for each temperature/humidity combination), lines show a 68 % (thick) and 95 % (thin) credible interval. Dashed line shows the ERH. Estimated evaporation phase half-lives plotted at the right. Colored lines show predicted half-lives as a function of humidity at five temperatures: 0 °C, 10 °C, 22 °C, 27 °C, and 40 °C. 100 random draws from the posterior distribution are shown at each temperature to visualize uncertainty. Line and point colors indicate temperature. (**b**) Predicted half-life above the ERH as a function of quasi-equilibrium concentration factor, according to the measured concentration fit. Points and lines as in **a**, but only solution (above ERH) conditions shown. (**c**) Heatmap showing posterior median predicted half-lives from the modeled concentration fit as a function of temperature and relative humidity. Posterior median estimated half-lives for human coronaviruses from our study and from the literature plotted on top (see also Appendix Table A2 and Fig. A7). Shape indicates virus; measurements from our own group are shown slightly larger with a slightly thicker outline. (**d**) Comparison of model-free estimates (x-axis) to model predictions (y-axis) for human coronavirus half-lives. Points show posterior median for measured (horizontal) or predicted (vertical) half-lives and lines show a 68 % (thick) and 95 % (thin) credible interval. Shape indicates virus; only data from the literature are shown.

As a stronger test of our model’s validity, we used our estimated *E*_*a*_ and *A* values to make out-of-sample predictions of the half-lives of five human coronaviruses reported from independent studies: four betacoronaviruses (SARS-CoV-2, SARS-CoV-1, MERS-CoV and HCoV-OC43) and one alphacoronavirus (HCoV-229E). We compiled data on the environmental stability of those viruses under conditions ranging from 4 to 95 °C, from 30 to 80 % RH, and on a range of surfaces or bulk media, and computed empirical—model-free—estimates of virus half-lives (Appendix Tables A5– A2).

Where both temperature and RH were available, we compared these model-free estimates to predictions based on the mechanistic model parameterized with our SARS-CoV-2 data (Fig. 3c, Appendix Fig. A7). We found striking agreement for half-life estimates both above and below the ERH, and for temperatures ranging from 4 to 37 °C.

To include a broader range of conditions in our out-of-sample model testing, we used our model to predict half-lives observed in all comparable studies by extrapolating from a reference half-life in each study. Predicted half-lives matched observations well across five orders of magnitude (Fig. 3d), despite spanning five virus species and despite important heterogeneities in the data collection process (Appendix section 6). The two conspicuous outliers, where SARS-CoV-2 half-lives were measured to be substantially shorter than our prediction, correspond to samples exposed to high heat in closed vials [13, 12] which is known to accelerate virus inactivation [21].

## Discussion

Combining novel data, mathematical modeling, and a meta-analysis of existing literature, we have developed a unified, mechanistic framework to quantify the joint effects of temperature and humidity on virus stability. In particular, our model provides a mechanism for the non-linear and non-monotonic relationship between relative humidity and virus stability previously observed for numerous enveloped viruses [64, 9, 55], but not previously reported for SARS-CoV-2. Our work documents and explains the strong dependence of SARS-CoV-2 stability on environmental temperature and relative humidity, and accurately predicts half-lives for five coronavirus species in conditions from 4 to 95 °C, and from 30 to 80 % RH and in bulk solution.

Our findings have direct implications for the epidemiology and control of SARS-CoV-2 and other enveloped viruses. The majority of SARS-CoV-2 clusters have been linked to indoor settings [36], suggesting that virus stability in indoor environmental conditions may be an important determinant of superspreading risk. Our results provide a mechanistic explanation for the many observed SARS-CoV-2 superspreading events in cool indoor environments such as food processing plants [17, 23, 49] and hockey rinks [3, 43], where the typical air temperature is around 10 °C, or in dry indoor environments such as long-distance flights [32, 26]. Conversely, our results imply that the relative rarity of outdoor SARS-CoV-2 transmission clusters is not readily explained by temperature and humidity effects, since these conditions outdoors during temperate winters should be favorable for the virus. Instead, increased ventilation [51] and UV light inactivation [53] may be more important than the effects of temperature and humidity outdoors. In contrast, typical climate-controlled conditions indoors (moderate temperature and low humidity) are favorable for virus stability, and specialized conditions such as those found in food processing plants even more so. Our results highlight the importance of proper personal protective equipment and improved ventilation for protecting workers, particularly in cold indoor settings, and the general transmission risks associated with indoor gatherings.

The effects of temperature and humidity we observe in our data and model are relevant both to fomite and to airborne transmission. Prior work has shown that virus decay as a function of RH is similar in droplets on surfaces and suspended aerosols [37, 34]. Numerous studies of smaller deposited droplets [52] or aerosols [5, 63, 25] have reported similar qualitative patterns to those we report, with increased decay rates at high temperatures and a U-shaped effect of RH. Furthermore, surface stability can matter for aerosol transmission risk, since small particles containing infectious virions can be re-suspended from surfaces and inhaled [2]. Re-suspension is further enhanced by procedures such as high-pressure washing, which is common in food processing plants. While the relative contributions of aerosol and fomite transmission to the epidemiology of SARS-CoV-2 continue to be investigated [47, 7], our results indicate that cold situations present elevated transmission risks for either mode, especially if air is either dry or very humid. It has been speculated, for instance, that chilled or frozen foods might allow for rare but impactful long-range fomite transmission [19]. Our results show that this is conceivable, as there is good empirical and mechanistic support for prolonged virus viability at very low temperatures.

Environmental stability is not the only mechanism by which temperature and humidity affect respiratory virus transmission. Very hot or cold conditions outdoors can lead people to spend more time indoors, where transmission risks are heightened due to poor ventilation. Low-humidity environments can dry out human airways and thus impair defenses against respiratory viruses [35]. Ambient humidity also determines the size distribution of aerosols in the environment, again by affecting evaporation rates. Smaller aerosols settle to the ground more slowly [40], which could facilitate transmission.

At low RH, humidity effects on inactivation, immunity, and settling may compound each other: all increase transmission risk. At high RH, reduced inactivation could promote transmission, but improved immune defenses and faster settling could hinder it, so the net effect on transmission is less clear.

Still, temperate winters increase transmission of many respiratory viruses [39]. Individuals spend increased time indoors in heated buildings. Ventilation is often poor, as windows are kept closed to make heating efficient. Air in heated buildings is typically very dry; this improves virus stability and weakens immune defenses. Policymakers should consider ventilating and humidifying essential indoor spaces to reduce transmission risk. Other mitigation measures such as indoor masking may likewise be even more crucial during winter. Indoor spaces in which individuals cannot be masked, such as bars and restaurants, remain particular cause for concern.

Several analyses have projected that SARS-CoV-2 transmission will likewise be faster in temperate winters [46, 33, 4]. Major seasonal or climate-mediated mitigation of SARS-CoV-2 spread was not evident during the northern hemisphere’s spring and summer [8, 48]. This was expected, since population susceptibility and epidemic control measures can be more important than seasonality in an early pandemic context [4]. Thus the fact that temperate summers did not eliminate transmission should not lead to false confidence that temperate winters will not promote it. Recent surges in cases, hospitalizations, and deaths across the northern hemisphere may have been driven in part by behavioral, immunological, or virological seasonality.

Our work has implications for the study of virus environmental stability and seasonality more broadly. Whether absolute or relative humidity is more important for influenza stability has been a matter of debate [54, 40]. The answer has proved elusive because it is difficult to disentangle the effects of humidity from those of temperature. Our mechanistic model permits principled dis-aggregation of those effects, and reveals a strong effect of relative humidity even after accounting for the effects of temperature.

There may thus exist general principles that govern virus inactivation across enveloped viruses, and perhaps even more broadly. Similar empirical patterns of temperature and humidity dependence to what we measured, and modeled, for SARS-CoV-2 have been observed for other important viruses. In particular, the U-shaped dependence of inactivation on RH has been reported for animal coronaviruses [55, 9], as well as for influenza viruses, paramyxoviruses, rhabdoviruses and retroviruses [63, 5, 52, 60], suggesting the existence of a shared mechanism for the effect of humidity across enveloped RNA viruses. Some enveloped DNA viruses such as herpesviruses and poxviruses [55, 60] and some encapsulated viruses such as polioviruses [14, 55] also show similar empirical behavior. Experiments have found that heat treatment of viruses reduces infectivity principally by degrading surface proteins [62], lending further support to a chemical model of environmental virus inactivation. We discuss additional practical implications for the empirical study of virus environmental stability in the Appendix (section 7).

Despite years of research on virus stability as a function of temperature and humidity and plausible hypotheses about the underlying chemistry, proposed mechanisms have lacked explicit quantitative support. By encoding the underlying chemistry into a mathematical model and estimating parameters using modern computational techniques, we provide such support, with critical insights for the control of an ongoing pandemic. Our empirical results provide mechanistic insight into transmission risks associated with cold and climate controlled indoor settings, while our modeling work allows for explicit quantitative comparison of the aerosol and fomite risks in different environments, and suggests that simple, general mechanisms govern the viability of enveloped viruses: hotter, more concentrated solutions are favorable to chemical reactions—and therefore unfavorable to viruses.

## Methods

### Laboratory experiments

#### Viruses and titration

We used SARS-CoV-2 strain HCoV-19 nCoV-WA1-2020 (MN985325.1) [24] for this study. We quantified viable virus by end-point titration on Vero E6 cells as described previously [18, 15], and inferred posterior distributions for titers and exponential decay rates directly from raw titration data using Bayesian statistical models (see Statistical analyses and mathematical modeling below).

#### Virus stability experiment

We measured virus stability on polypropylene (ePlastics, reference PRONAT.030X24X47S/M) as previously described [15]. We prepared a solution of Dulbecco’s Modified Eagle Medium (DMEM, a common cell culture medium) supplemented with 2 mM L-glutamine, 2% fetal bovine serum and 100 units/mL penicillin/streptomycin, and containing 105 TCID_50_/mL SARS-CoV-2. Polypropylene disks were autoclaved for decontamination prior to the experiment. We then placed 50 µL aliquots of this SARS-CoV-2 suspension onto the polypropylene disks under nine environmental conditions: three RH (40 %, 65 %, and 85 %) at each of three temperatures (10 °C, 22 °C, and 27 °C).These controlled environmental conditions were produced in incubators (MMM Group CLIMACELL and Caron model 6040) with protection from UV-B or UV-C exposure. We prepared 216 disks corresponding to three replicates per eight post-deposition time-points (0, 1, 4, 8, and 24 hours, then daily for 4 days) for the nine conditions. At each time-point, samples were collected by rinsing the disks with 1 mL of DMEM and stored at −80 °C until titration.

#### Evaporation experiment

We measured the evaporation kinetics of suspension medium under the same temperature and humidity conditions as the virus stability experiments. We placed 50 µL aliquots of supplemented DMEM onto polypropylene disks in a Electro-Tech Systems 5518 environmental chamber. The polypropylene disks were rinsed three times 1M sulfuric acid, ethanol and DI H2O respectively before use. We measured medium mass *<* (*C*) every 5 min for up to 20 h or until a quasi-equilibrium was reached using a micro-balance (Sartorius MSE3.6P-000-DM, readability 0.0010 mg). The chamber of the micro-balance was half-opened to keep air circulating with the environmental chamber. The flow entering the balance chamber decreased the balance accuracy to around 0.010 mg. We measured initial droplet mass (*m*(0)) and final droplet mass (*m*(∞)) under closed-chamber conditions to increase accuracy.

### Statistical analyses and mathematical modeling

We quantified the stability of SARS-CoV-2 under different conditions by estimating the decay rates of viable virus titers. We inferred individual titers using a Bayesian model we have previously described [21]. Briefly, the model treats titration well infection as a Poisson single-hit process. We inferred raw exponential decay rates by modifying a previously-described simple regression model [21] to account for the evaporation phase. See the Appendix (section 5.4) for model description.

We estimated parameters of our mechanistic models by predicting titers based on those models and then applying the same Poisson single-hit observation process to estimate parameters from the data. See the Appendix (section 5.5) for a complete description, including model priors.

We estimated evaporation rates and corresponding drying times by modeling mass loss for each environmental condition *8* as linear in time at a rate *V*_*8*_ until the final mass *<* (1) was reached, See the Appendix (sections 4.2, 5.3) for a full description including model priors.

We drew posterior samples using Stan [57], which implements a No-U-Turn Sampler (a form of Markov Chain Monte Carlo), via its R interface RStan [56].

### Meta-analysis

To test the validity of our model beyond the measured environmental conditions (i.e., beyond 10– 27 °C and 40–85 % RH), we compiled data from 11 published studies on human coronaviruses, including SARS-CoV-2, SARS-CoV-1, MERS-CoV, HCoV-OC43 and HCoV-299E, under 17 temperature-RH conditions. We generated estimates of half-life and uncertainties (Appendix Table A2) and compared those estimates to the half-lives predicted by the mechanistic model parametrized from our SARS-CoV-2 data. As data on evaporation kinetics were not available, we estimated a unique half-life for each experimental condition, covering both the evaporation and quasi-equilibrium phases. As virus decay during the evaporation phase is expected to be minimal, and the evaporation phase to be short, the estimated half-life can be used as a proxy for the quasi-equilibrium half-life. The complete data selection, extraction and analysis process is detailed in the Appendix (section 6).

We also included data from SARS-CoV-1 and MERS-CoV collected by our group during previous studies [15]. Those data were collected at 22 °C and 40 % RH on polypropylene using the protocol described previously [15] and similar to the one used to collect the SARS-CoV-2 data. SARS-CoV-1 strain Tor2 (AY274119.3) [41] and MERS-CoV strain HCoV-EMC/2012 were used for these experiments. We calculated half-lives for evaporation and quasi-equilibrium phases using the same analysis pipeline used for SARS-CoV-2 (Appendix section 5.4). These data were used only for out-of-sample prediction testing. We used the obtained evaporation phase half-lives as proxies for the half-life at 100 % RH, as with SARS-CoV-2. See Appendix section 6.3 for a figure showing model fits (Fig. A32) and a table of estimated half-lives (Table A5).

### Visualization

We created plots in R version using ggplot2 [61], ggdist [30], and tidybayes [31], and created original schematics using BioRender.com.

## Acknowledgments

This research was supported by the Intramural Research Program of the National Institute of Allergy and Infectious Diseases (NIAID), National Institutes of Health (NIH). DHM was supported by the U.S. National Science Foundation (CCF 1917819). JOL-S and AG were supported by the Defense Advanced Research Projects Agency DARPA PREEMPT #D18AC00031 and the UCLA AIDS Institute and Charity Treks, and JOL-S was supported by the U.S. National Science Foundation (DEB-1557022), the Strategic Environmental Research and Development Program (SERDP, RC-2635) of the U.S. QH, LCM, and PJV were supported by the U.S. National Science Foundation (CBET-1705653, CBET-2029911). The content of the article does not necessarily reflect the position or the policy of the U.S. government, and no official endorsement should be inferred.

## Author contributions

VJM and JOL-S conceived the study. KCY performed the inactivation experiments, with support from TB, RJF, MJM, NvD, and VJM. QH performed the evaporation experiments, with support from PJV and LCM. DHM developed the mechanistic model and conducted statistical and theoretical analysis, with support from JOL-S, AG, FWR, and LCM. DHM wrote analysis code and produced figures and schematics. AG reviewed the literature on coronavirus stability and compiled and prepared data for meta-analysis. DHM, AG, and JOL-S drafted the paper, which all authors edited.

## Code and data availability

All code and data needed to reproduce results and figures is archived on Github (https://github.com/dylanhmorris/sars-cov-2-temp-humidity/) and on Zenodo (https://doi.org/10.5281/zenodo.4093264), and licensed for reuse, with appropriate attribution/citation, under a BSD 3-Clause Revised License.

## References

[1] Mark H Adams. “The stability of bacterial viruses in solutions of salts”. In: The Journal of general physiology 32.5 (1949), p. 579.

[2] Sima Asadi et al. “Influenza A Virus Is Transmissible via Aerosolized Fomites”. In: Nature Communications 11 (Aug. 18, 2020), p. 4062. ISSN: 2041-1723. DOI: 10.1038/s41467-020-17888-w.

[3] David Atrubin, Michael Wiese, and Becky Bohinc. “An Outbreak of COVID-19 Associated with a Recreational Hockey Game-Florida, June 2020”. In: Morbidity and Mortality Weekly Report 69.41 (2020), p. 1492.

[4] Rachel E. Baker et al. “Susceptible supply limits the role of climate in the early SARS-CoV-2 pandemic”. In: Science (2020). DOI: 10.1126/science.abc2535.

[5] J. E. Benbough. “Some factors affecting the survival of airborne viruses”. In: The Journal of General Virology 10 (1971), pp. 209–220. DOI: 10.1099/0022-1317-10-3-209.

[6] Jennifer Biryukov et al. “Increasing Temperature and Relative Humidity Accelerates Inactivation of SARS-CoV-2 on Surfaces”. In: Msphere 5.4 (2020).

[7] Jing Cai et al. “Indirect Virus Transmission in Cluster of COVID-19 Cases, Wenzhou, China, 2020”. In: Emerging Infectious Diseases 26 (2020). DOI: 10.3201/eid2606.200412.

[8] Colin J Carlson et al. “Misconceptions about weather and seasonality must not misguide COVID-19 response”. In: Nature Communications 11.1 (2020), pp. 1–4.

[9] Lisa M Casanova et al. “Effects of air temperature and relative humidity on coronavirus survival on surfaces”. In: Applied and environmental microbiology 76.9 (2010), pp. 2712–2717. DOI: 10.1128/AEM.02291-09.

[10] Franco Cavaliere et al. “Airway secretion electrolytes: reflection of water and salt states of the body.” In: Critical care medicine 17.9 (1989), pp. 891–894.

[11] Kwok-Hung Chan et al. “The effects of temperature and relative humidity on the viability of the SARS coronavirus”. In: Advances in Virology 2011 (2011). DOI: 10.1155/2011/734690.

[12] Alex W. H. Chin. personal communication. Aug. 5, 2020.

[13] Alex W. H. Chin et al. “Stability of SARS-CoV-2 in different environmental conditions”. In: The Lancet Microbe 1 (2020), e10. DOI: 10.1016/S2666-5247(20)30003-3.

[14] JC De Jong and KC Winkler. “The inactivation of poliovirus in aerosols”. In: Epidemiology & Infection 66.4 (1968), pp. 557–565.

[15] Neeltje van Doremalen et al. “Aerosol and surface stability of SARS-CoV-2 as compared with SARS-CoV-1”. In: New England Journal of Medicine 382 (2020), pp. 1564–1567. DOI: 10.1056/NEJMc2004973.

[16] R Dulbecco and G Freeman. “Plaque production by the polyoma virus.” In: Virology 8.3 (1959), p. 396.

[17] Jonathan W. Dyal. “COVID-19 Among Workers in Meat and Poultry Processing Facilities — 19 States, April 2020”. In: Morbidity and Mortality Weekly Report 69 (2020). DOI: 10.15585/mmwr.mm6918e3.

[18] Robert Fischer et al. “Assessment of N95 respirator decontamination and re-use for SARS-CoV-2”. In: Emerging Infectious Diseases In press (2020). DOI: 10.1101/2020.04.11.20062018.

[19] Dale Fisher et al. “Seeding of outbreaks of COVID-19 by contaminated fresh and frozen food”. In: bioRxiv (2020).

[20] Yuki Furuse et al. “Clusters of coronavirus disease in communities, Japan, January–April 2020”. In: Emerging infectious diseases 26.9 (2020), p. 2176.

[21] Amandine Gamble et al. “Heat-treated virus inactivation rate depends strongly on treatment procedure”. In: bioRxiv (2020). DOI: 10.1101/2020.08.10.242206.

[22] Laurent Guillier et al. “Modeling the inactivation of viruses from the coronaviridae family in response to temperature and relative humidity in suspensions or on surfaces”. In: Applied and Environmental Microbiology 86.18 (2020). DOI: 10.1128/AEM.01244-20.

[23] Thomas Günther et al. “Investigation of a superspreading event preceding the largest meat processing plant-related SARS-Coronavirus 2 outbreak in Germany”. In: SSRN (2020). DOI: 10.2139/ssrn.3654517.

[24] Michelle L. Holshue et al. “First Case of 2019 Novel Coronavirus in the United States”. In: New England Journal of Medicine 382.10 (2020), pp. 929–936. DOI: 10.1056/NEJMoa2001191.

[25] M. K. Ijaz et al. “Survival Characteristics of Airborne Human Coronavirus 229E”. In: The Journal of General Virology 66 (Pt 12) (Dec. 1985), pp. 2743–2748. ISSN: 0022-1317. DOI: 10.1099/0022-1317-66-12-2743.

[26] Mahesh Jayaweera et al. “Transmission of COVID-19 virus by droplets and aerosols: A critical review on the unresolved dichotomy”. In: Environmental Research (2020), p. 109819.

[27] David Johnson et al. “Aerosol Generation by Modern Flush Toilets”. In: Aerosol Science and Technology 47 (Sept. 1, 2013), pp. 1047–1057. ISSN: 0278-6826. DOI: 10.1080/02786826.2013.814911.

[28] G.R. Johnson et al. “Modality of Human Expired Aerosol Size Distributions”. In: Journal of Aerosol Science 42 (Dec. 2011), pp. 839–851. ISSN: 00218502. DOI: 10.1016/j.jaerosci.2011.07.009.

[29] Morgan P Kain et al. “Chopping the tail: how preventing superspreading can help to maintain COVID-19 control”. In: MedRxiv (2020).

[30] Matthew Kay. ggdist: Visualizations of Distributions and Uncertainty. R package version 2.2.0. 2020. DOI: 10.5281/zenodo.3879620. URL: http://mjskay.github.io/ggdist/.

[31] Matthew Kay. tidybayes: Tidy Data and Geoms for Bayesian Models. R package version 2.1.1. 2020. DOI: 10.5281/zenodo.1308151. URL: http://mjskay.github.io/tidybayes/.

[32] Nguyen Cong Khanh et al. “Transmission of Severe Acute Respiratory Syndrome Coronavirus 2 During Long Flight”. In: Emerging Infectious Diseases (2020). DOI: 10.3201/eid2611.203299.

[33] Stephen M. Kissler et al. “Projecting the transmission dynamics of SARS-CoV-2 through the postpandemic period”. In: Science (2020). ISSN: 0036-8075, 1095-9203. DOI: 10.1126/science.abb5793. URL: https://science.sciencemag.org/content/early/2020/04/14/science.abb5793.

[34] Karen A Kormuth et al. “Influenza virus infectivity is retained in aerosols and droplets independent of relative humidity”. In: The Journal of infectious diseases 218.5 (2018), pp. 739–747.

[35] Eriko Kudo et al. “Low ambient humidity impairs barrier function and innate resistance against influenza infection”. In: Proceedings of the National Academy of Sciences 116.22 (2019), pp. 10905–10910.

[36] Quentin J. Leclerc et al. “What Settings Have Been Linked to SARS-CoV-2 Transmission Clusters?” In: Wellcome Open Research 5 (June 5, 2020), p. 83. ISSN: 2398-502X. DOI: 10.12688/wellcomeopenres.15889.2.

[37] Kaisen Lin and Linsey C Marr. “Humidity-dependent decay of viruses, but not bacteria, in aerosols and droplets follows disinfection kinetics”. In: Environmental Science & Technology 54.2 (2020), pp. 1024–1032. DOI: 10.1021/acs.est.9b04959.

[38] James O Lloyd-Smith et al. “Superspreading and the effect of individual variation on disease emergence”. In: Nature 438.7066 (2005), pp. 355–359.

[39] Eric Lofgren et al. “Influenza Seasonality: Underlying Causes and Modeling Theories”. In: Journal of Virology 81.11 (2007), pp. 5429–5436. ISSN: 0022-538X, 1098-5514. DOI: 10.1128/JVI.01680-06.

[40] Linsey C Marr et al. “Mechanistic insights into the effect of humidity on airborne influenza virus survival, transmission and incidence”. In: Journal of the Royal Society Interface 16.150 (2019), p. 20180298.

[41] Marco A Marra et al. “The genome sequence of the SARS-associated coronavirus”. In: Science 300.5624 (2003), pp. 1399–1404.

[42] M. Jeremiah Matson et al. “Effect of Environmental Conditions on SARS-CoV-2 Stability in Human Nasal Mucus and Sputum”. In: Emerging Infectious Diseases 26 (2020), in press. DOI: 10.3201/eid2609.202267.

[43] Nakia McNabb and Brian Ries. “Vermont coronavirus cluster traced to hockey teams and a broomball league”. In: CNN (Oct. 2020). URL: https://www.cnn.com/2020/10/20/us/vermont-ice-rink-covid-trnd/index.html.

[44] E Mikhailov et al. “Interaction of aerosol particles composed of protein and saltswith water vapor: hygroscopic growth and microstructural rearrangement”. In: (2004).

[45] Arnold S Monto et al. “Coronavirus occurrence and transmission over 8 years in the HIVE cohort of households in Michigan”. In: The Journal of infectious diseases 222.1 (Apr. 2020), pp. 9–16. DOI: 10.1093/infdis/jiaa161.

[46] Richard A Neher et al. “Potential impact of seasonal forcing on a SARS-CoV-2 pandemic”. In: Swiss medical weekly 150.1112 (2020).

[47] Sean Wei Xiang Ong et al. “Air, Surface Environmental, and Personal Protective Equipment Contamination by Severe Acute Respiratory Syndrome Coronavirus 2 (SARS-CoV-2) From a Symptomatic Patient”. In: JAMA (Mar. 4, 2020). DOI: 10.1001/jama.2020.3227.

[48] Canelle Poirier et al. “The Role of Environmental Factors on Transmission Rates of the COVID-19 Outbreak: An Initial Assessment in Two Spatial Scales”. en. In: Scientific Reports 10.1 (Oct. 2020), p. 17002. ISSN: 2045-2322. DOI: 10.1038/s41598-020-74089-7.

[49] Roman Pokora et al. “Investigation of superspreading COVID-19 outbreaks events in meat and poultry processing plants in Germany: A cross-sectional study”. In: arXiv preprint (2020). URL: https://arxiv.org/abs/2011.11153.

[50] JA Posada, J Redrow, and I Celik. “A mathematical model for predicting the viability of airborne viruses”. In: Journal of virological methods 164.1-2 (2010), pp. 88–95.

[51] Kimberly A Prather, Chia C Wang, and Robert T Schooley. “Reducing transmission of SARS-CoV-2”. In: Science (2020).

[52] Aaron J Prussin et al. “Survival of the enveloped virus Phi6 in droplets as a function of relative humidity, absolute humidity, and temperature”. In: Applied and environmental microbiology 84.12 (2018). DOI: 10.1128/AEM.00551-18.

[53] Shanna Ratnesar-Shumate et al. “Simulated sunlight rapidly inactivates SARS-CoV-2 on surfaces”. In: The Journal of Infectious Diseases (2020).

[54] Jeffrey Shaman et al. “Absolute Humidity and the Seasonal Onset of Influenza in the Continental United States”. In: PLoS Biology 8.2 (2010), e1000316. ISSN: 1545-7885. DOI: 10.1371/journal.pbio.1000316.

[55] Joseph R Songer. “Influence of relative humidity on the survival of some airborne viruses”. In: Applied microbiology 15.1 (1967), pp. 35–42.

[56] Stan Development Team. “RStan: the R interface to Stan”. In: R package version 2.1 (2016).

[57] Stan Development Team. The Stan Core Library. Version 2.18.0. 2018. URL: https://mc-stan.org/.

[58] Katy-Anne Thompson et al. “Influenza Aerosols in UK Hospitals during the H1N1 (2009) Pandemic – The Risk of Aerosol Generation during Medical Procedures”. In: PLOS ONE 8 (Feb. 13, 2013), e56278. ISSN: 1932-6203. DOI: 10.1371/journal.pone.0056278.

[59] N Van Doremalen, T Bushmaker, and VJ Munster. “Stability of Middle East Respiratory Syndrome coronavirus (MERS-CoV) under different environmental conditions”. In: Eurosurveillance 18.38 (2013), p. 20590. DOI: 10.2807/1560-7917.ES2013.18.38.20590.

[60] SJ Webb, R Bather, and RW Hodges. “The effect of relative humidity and inositol on air-borne viruses”. In: Canadian Journal of Microbiology 9.1 (1963), pp. 87–92.

[61] Hadley Wickham. ggplot2: Elegant Graphics for Data Analysis. Springer-Verlag New York, 2016. ISBN: 978-3-319-24277-4. URL: https://ggplot2.tidyverse.org.

[62] Krista Rule Wigginton et al. “Virus inactivation mechanisms: impact of disinfectants on virus function and structural integrity”. In: Environmental Science & Technology 46 (2012), pp. 12069–12078. ISSN: 1520-5851. DOI: 10.1021/es3029473.

[63] Wan Yang, Subbiah Elankumaran, and Linsey C Marr. “Relationship between Humidity and Influenza A Viability in Droplets and Implications for Influenza’s Seasonality”. In: PloS ONE 7 (2012). DOI: 10.1371/journal.pone.0046789.

[64] Wan Yang and Linsey C Marr. “Mechanisms by Which Ambient Humidity May Affect Viruses in Aerosols”. In: Applied and Environmental Microbiology 78.19 (2012), pp. 6781–6788. DOI: 10.1128/AEM.01658-12.

[65] Te Faye Yap et al. “A Predictive Model of the Temperature-Dependent Inactivation of Coronaviruses”. In: (2020). DOI: 10.26434/chemrxiv.12152970.v4.

[66] Ali M Zaki et al. “Isolation of a novel coronavirus from a man with pneumonia in Saudi Arabia”. In: New England Journal of Medicine 367.19 (2012), pp. 1814–1820.

