## Appendix for "Mechanistic theory predicts the effects of temperature and humidity on inactivation of SARS-CoV-2 and other enveloped viruses"

December 17, 2020

**7 Methodological implications for experimental studies on virus stability 65**

### 1 Key additional figures

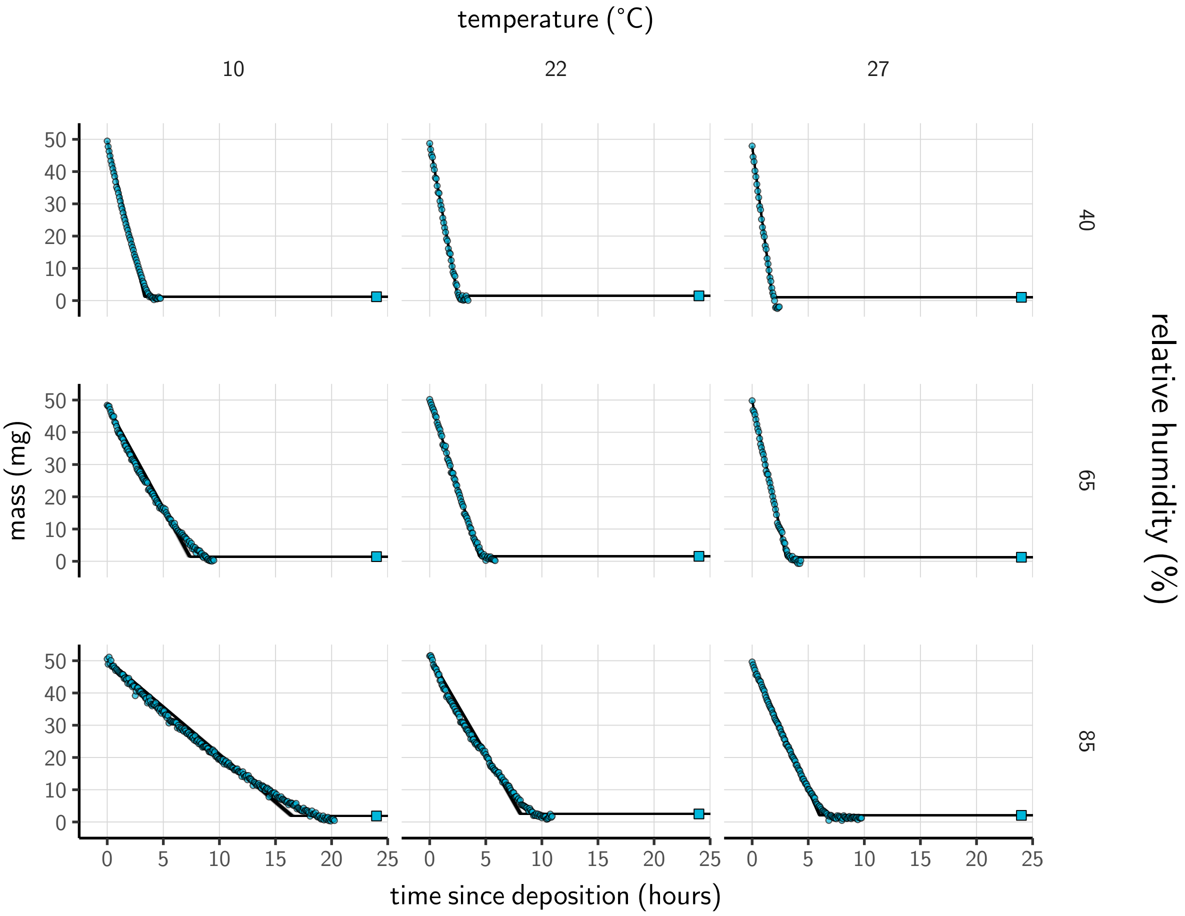

**Figure A1. Evaporation of supplemented Dulbecco's Modified Eagle Medium (DMEM) as a function of temperature and humidity.** Dots show measured masses. Square shows measured final (quasi-equilibrium) mass; actual measurement times for final masses were upon removal of sample from chamber, but for readability they are plotted at 20 h for all experiments. Lines are 100 random draws from the posterior for the evaporation rate; horizontal section of line reflects the reaching of quasi-equilibrium (measured final mass). Transition point between evaporation phase and quasi-equilibrium phase inferred from data (see SI sections 4.2, 5.3). Note that final mass measurement is more accurate than timeseries measurements (see the Evaporation experiment subsection of the Methods)

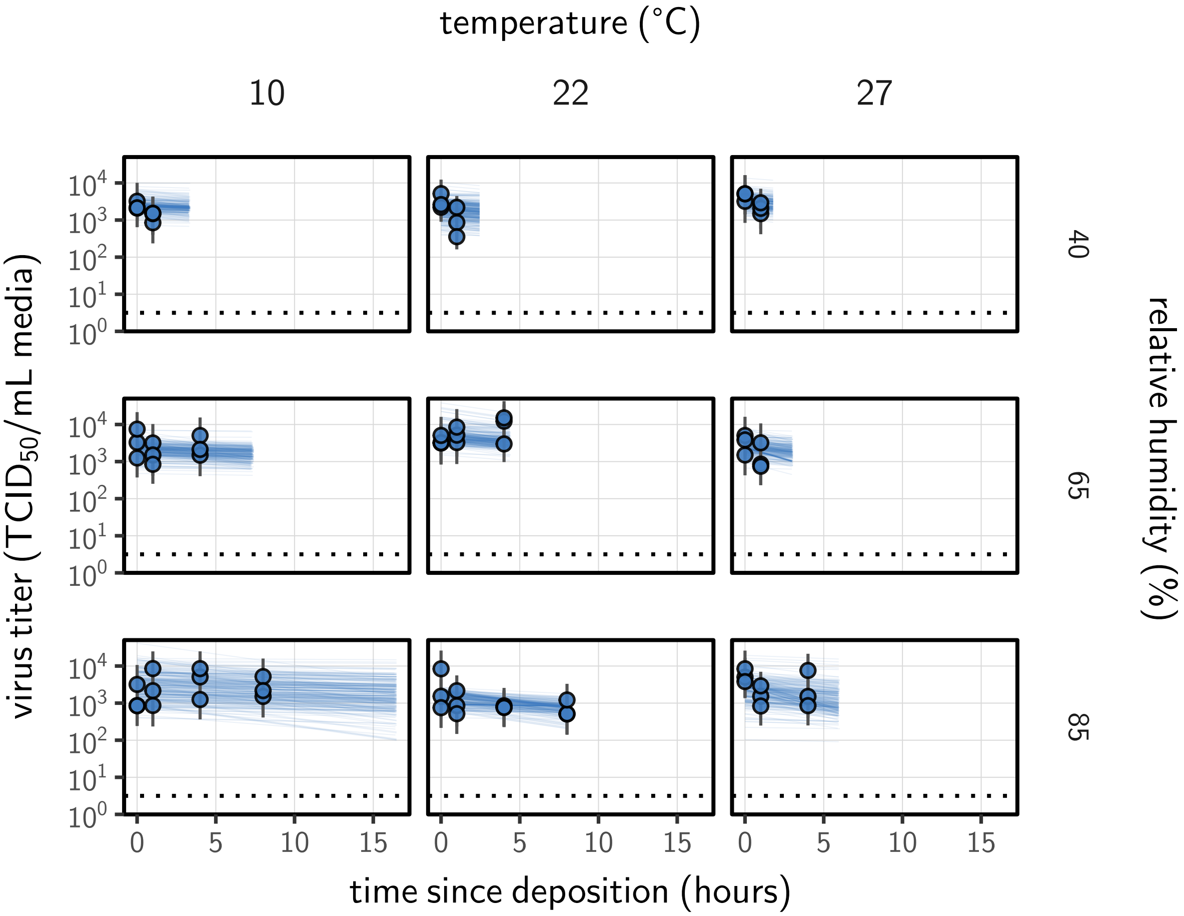

**Figure A2. Fit of the simple regression model to the evaporation phase (pre-drying) SARS-CoV-2 titer data.** Points show posterior median estimated titers in $\log_{10}$TCID$_{50}$/mL for each sample; lines show 100 % credible intervals. Time-points with no positive wells for any replicate are plotted as triangles at the approximate single-replicate limit of detection (LOD) of the assay—denoted by a black dotted line at $10^{0.5}$ TCID$_{50}$/mL media—to indicate that a range of sub-LOD values are plausible. Three samples collected at each time-point. x-axis shows time since sample deposition. Lines are truncated at the estimated time quasi-equilibrium was reached. Lines are random draws (10 per sample) from the joint posterior distribution of the initial sample virus concentration and the estimated decay rate; the distribution of lines gives an estimate of the uncertainty in the decay rate and the variability of the initial titer for each experiment.

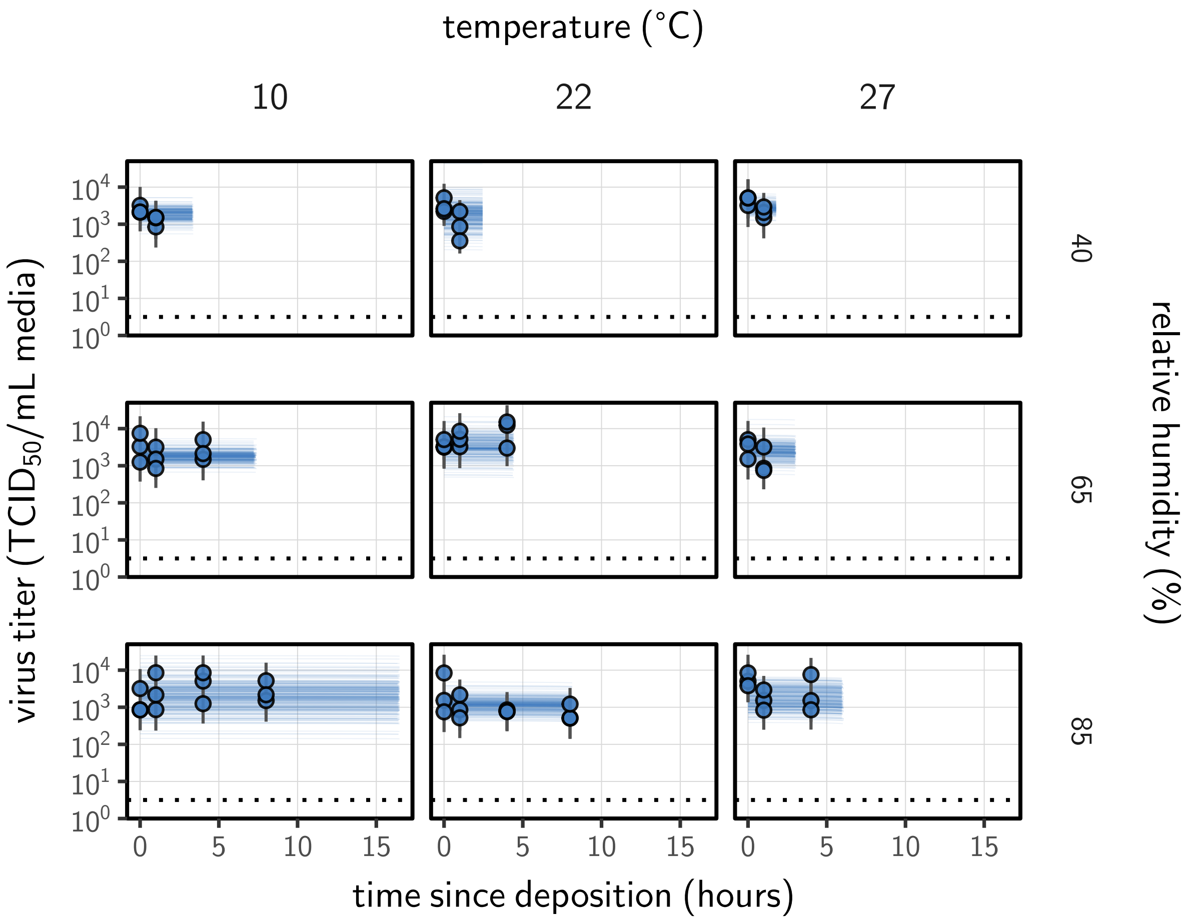

**Figure A3. Estimated titers and measured concentration model fit for SARS-CoV-2 stability on plastic during the evaporation phase.** Points show posterior median estimated titers in $\log_{10}$TCID$_{50}$/mL for each sample; lines show 100 % credible intervals. Time-points with no positive wells for any replicate are plotted as triangles at the approximate single-replicate limit of detection (LOD) of the assay—denoted by a black dotted line at $10^{0.5}$ TCID$_{50}$/mL media—to indicate that a range of sub-LOD values are plausible. Three samples collected at each time-point. x-axis shows time since sample deposition. Lines are truncated at the estimated time quasi-equilibrium was reached. Lines are random draws (10 per sample) from the joint posterior distribution of the initial sample virus concentration and the mechanistic model predicted decay rate; the distribution of lines gives an estimate of the uncertainty in the decay rate and the variability of the initial titer for each experiment.

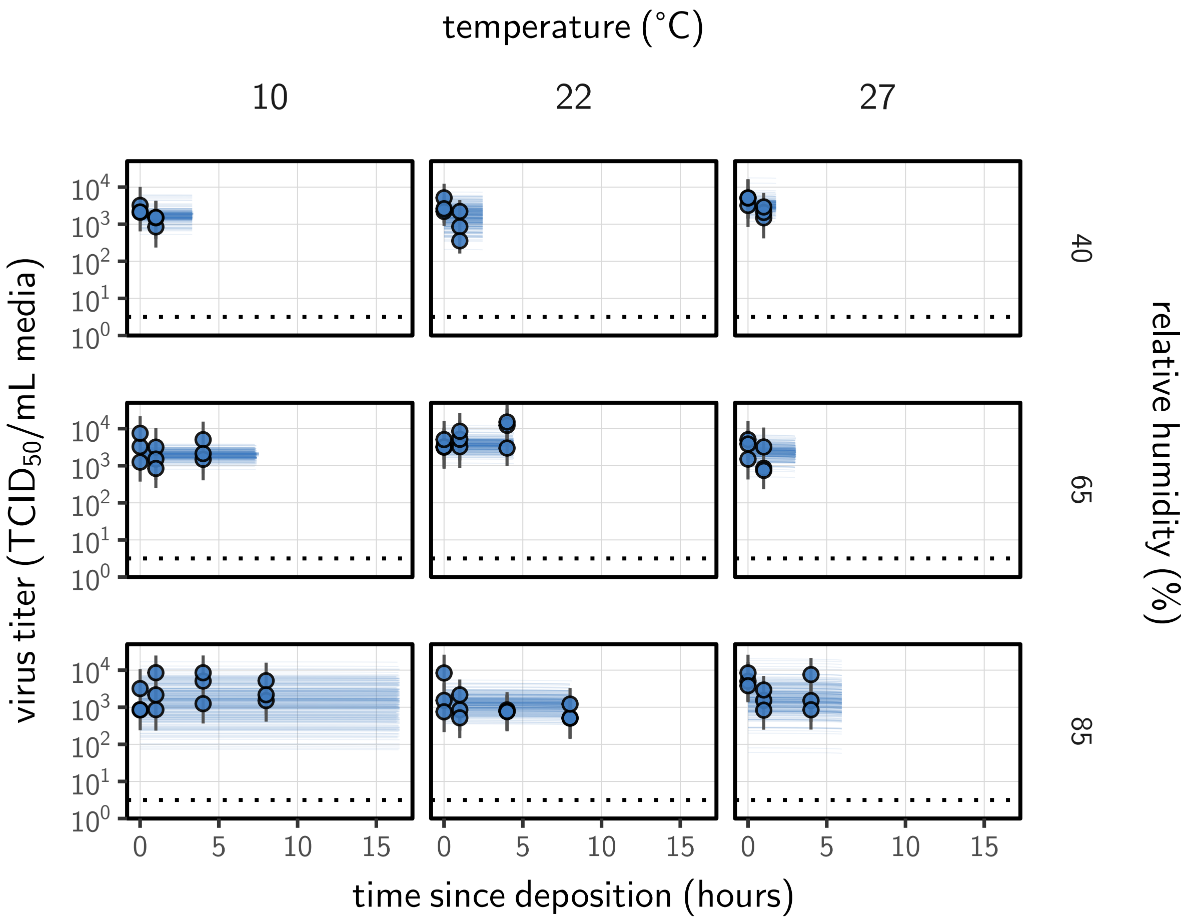

**Figure A4. Estimated titers and modeled concentration model fit for SARS-CoV-2 stability on plastic during the evaporation phase.** Points show posterior median estimated titers in $\log_{10}$TCID$_{50}$/mL for each sample; lines show 100 % credible intervals. Time-points with no positive wells for any replicate are plotted as triangles at the approximate single-replicate limit of detection (LOD) of the assay—denoted by a black dotted line at $10^{0.5}$ TCID$_{50}$/mL media—to indicate that a range of sub-LOD values are plausible. Three samples collected at each time-point. x-axis shows time since sample deposition. Lines are truncated at the estimated time quasi-equilibrium was reached. Lines are random draws (10 per sample) from the joint posterior distribution of the initial sample virus concentration and the mechanistic model predicted decay rate; the distribution of lines gives an estimate of the uncertainty in the decay rate and the variability of the initial titer for each experiment.

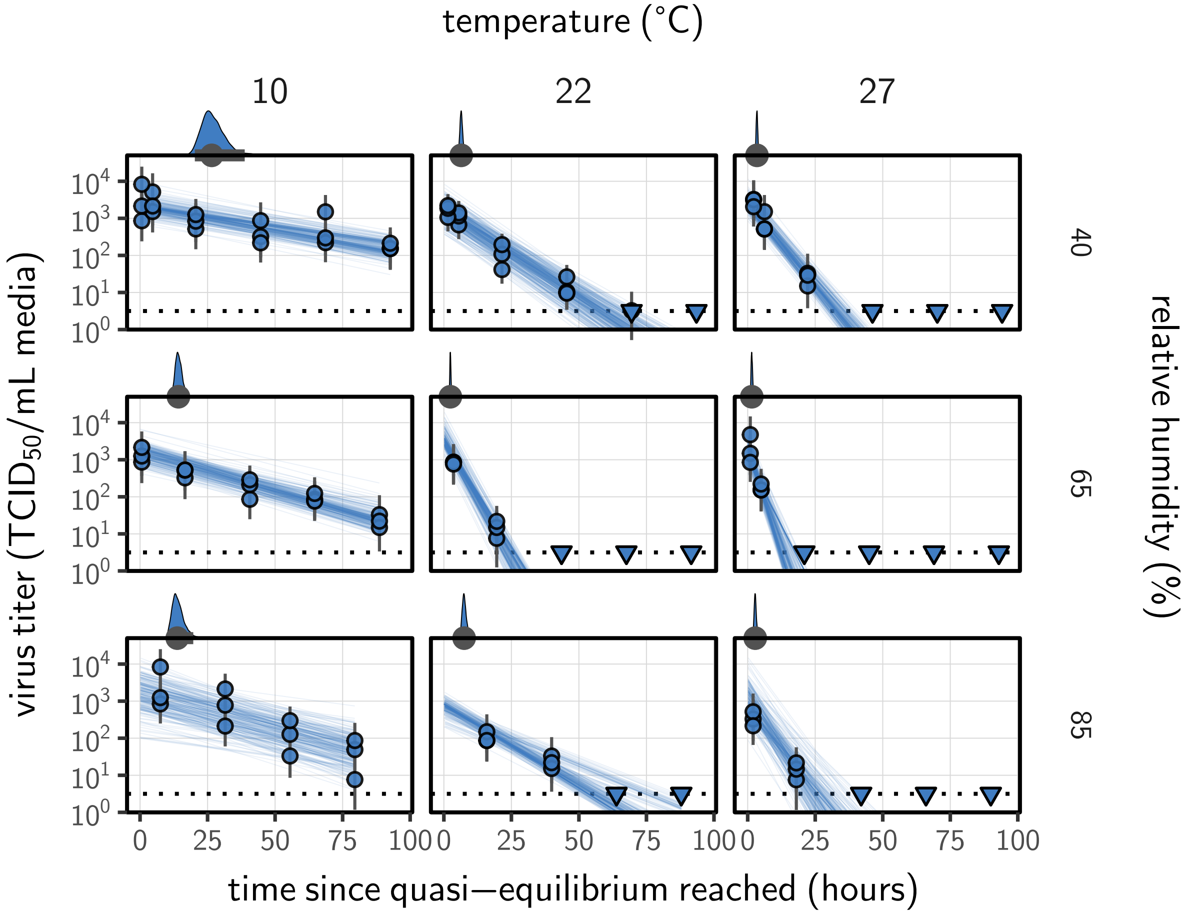

**Figure A5. Fit of the simple regression model to the quasi-equilibrium (post-drying) SARS-CoV-2 titer data.** Points show posterior median estimated titers in $\log_{10}$TCID$_{50}$/mL for each sample; lines show 100 % credible intervals. Time-points with no positive wells for any replicate are plotted as triangles at the approximate single-replicate limit of detection (LOD) of the assay—denoted by a black dotted line at $10^{0.5}$ TCID$_{50}$/mL media—to indicate that a range of sub-LOD values are plausible. Three samples collected at each time-point. x-axis shows time since quasi-equilibrium was reached, as measured in evaporation experiments. Lines are random draws (10 per sample) from the joint posterior distribution of the initial sample virus concentration and the estimated decay rate; the distribution of lines gives an estimate of the uncertainty in the decay rate and the variability of the initial titer for each experiment. Density plots above each box show posterior distribution of virus half-life according to the model for the given condition; point under the density shows the posterior median half-life and line shows a 100 % credible interval.

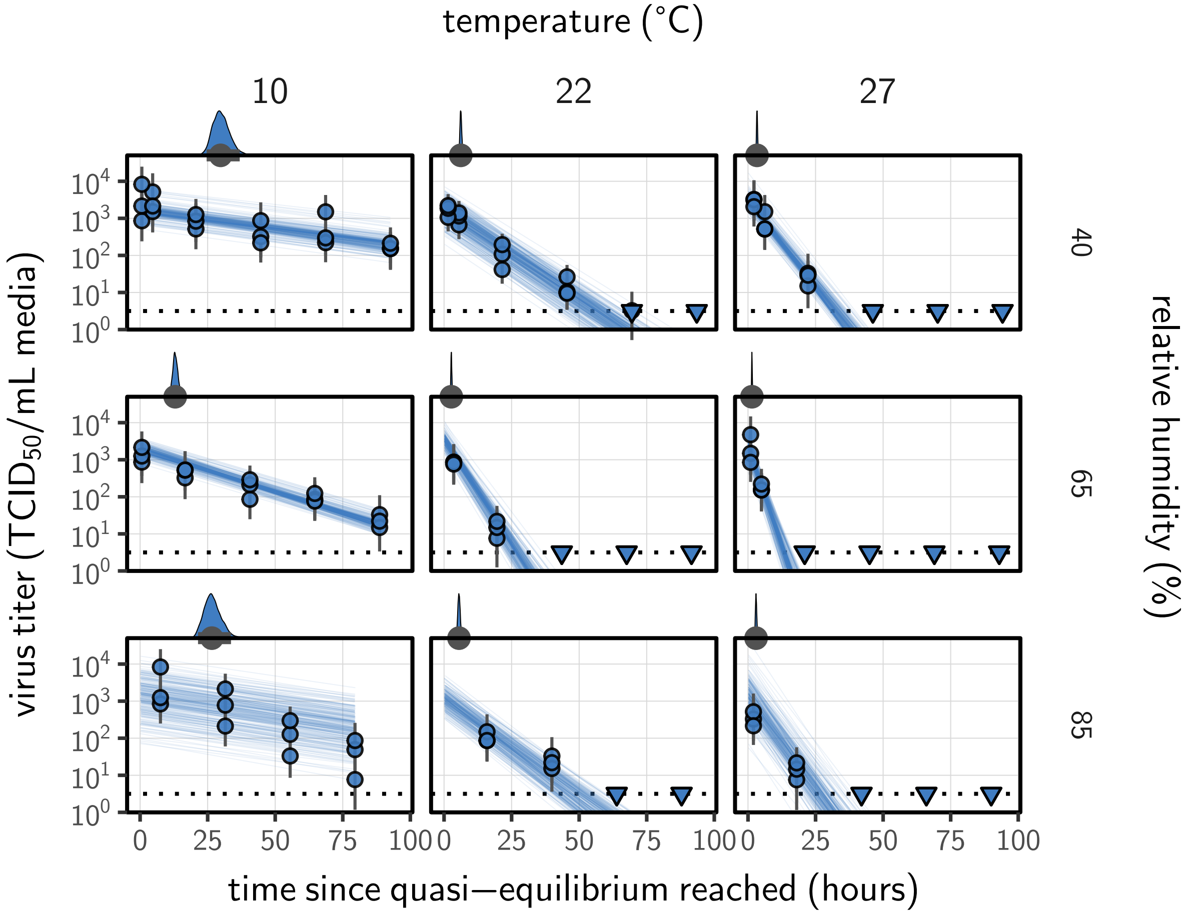

**Figure A6. Estimated titers and modeled concentration model fit for SARS-CoV-2 stability on plastic at quasi-equilibrium.** Points show posterior median estimated titers in $\log_{10}$TCID$_{50}$/mL for each sample; lines show 100 % credible intervals. Time-points with no positive wells for any replicate are plotted as triangles at the approximate single-replicate limit of detection (LOD) of the assay—denoted by a black dotted line at $10^{0.5}$ TCID$_{50}$/mL media—to indicate that a range of sub-LOD values are plausible. Three samples collected at each time-point. x-axis shows time since quasi-equilibrium was reached, as measured in evaporation experiments. Lines are random draws (10 per sample) from the joint posterior distribution of the initial sample virus concentration and the mechanistic model predicted decay rate; the distribution of lines gives an estimate of the uncertainty in the decay rate and the variability of the initial titer for each experiment. Density plots above each box show posterior distribution of virus half-life according to the model for the given condition; point under the density shows the posterior median half-life and line shows a 100 % credible interval.

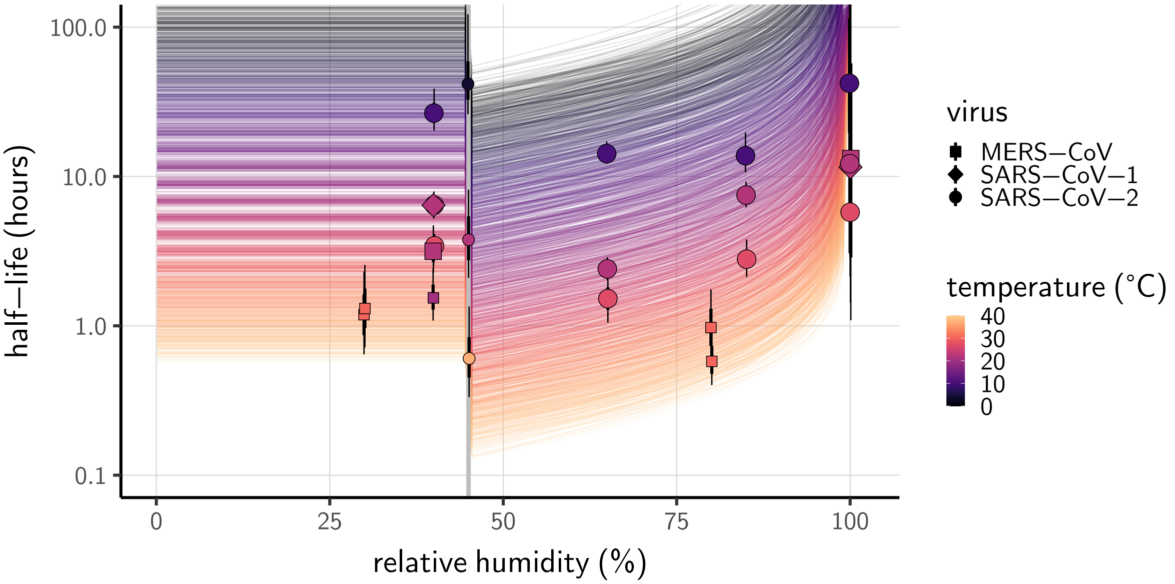

**Figure A7. Modeled concentration fit predictions compared to half-lives estimated directly from data.** Lines show half-life as a function of relative humidity (x-axis value) and temperature (color) according to the modeled concentration fit. 100 random draws from the posterior distribution are plotted for each of 20 evenly spaced temperatures between 0 and 40 °C. Points show posterior median for measured half-lives for human coronaviruses from our study and from the literature (Table A2); lines show a 70 % (thick) and 100 % (thin) credible interval. Half-lives estimates are model-free (i.e. no mechanistic model; fitting of independent exponential decay rates to each condition). Shape indicates virus; measurements from our own group are shown slightly larger. Estimated evaporation phase half-lives plotted at 100 % relative humidity (RH). Grey line shows the efflorescence relative humidity (ERH), 50 %.

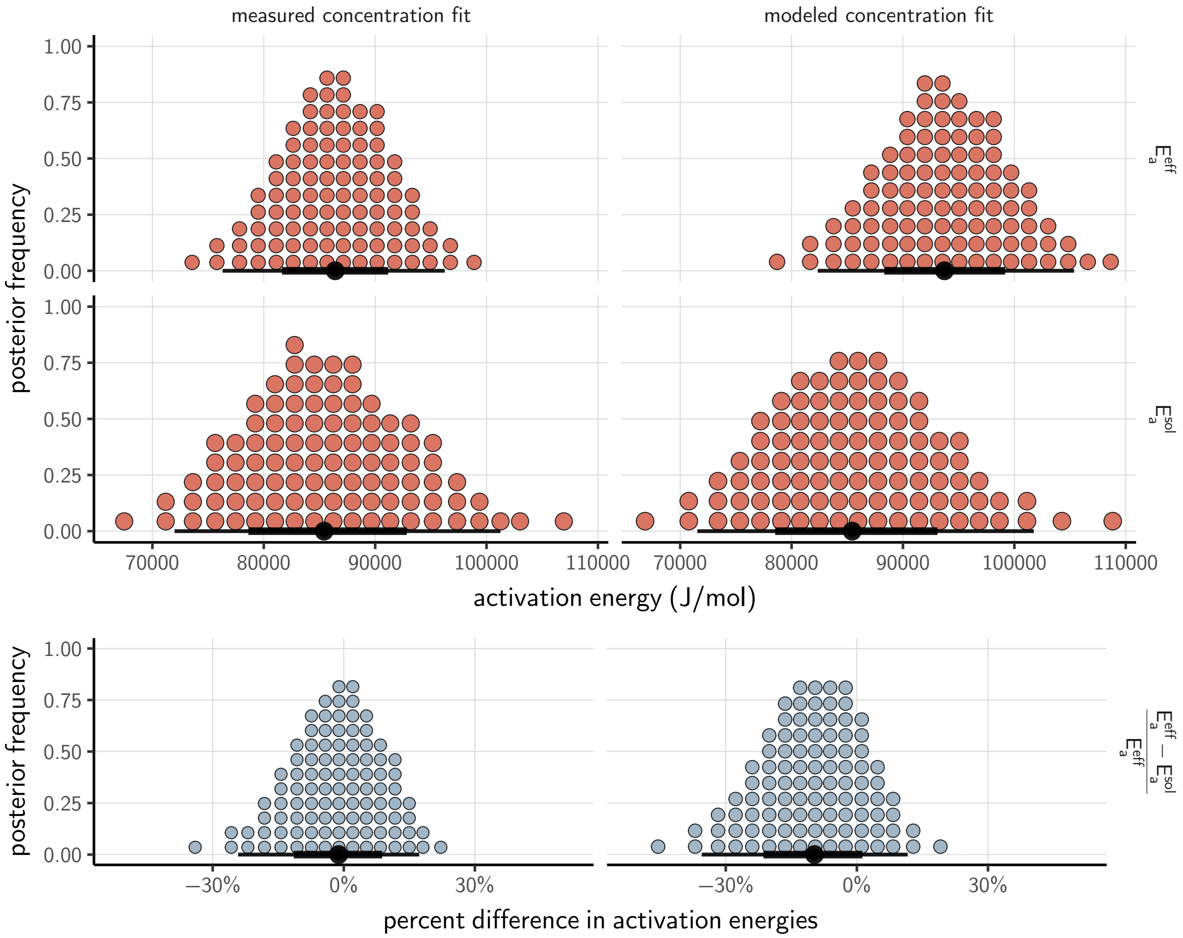

**Figure A8. Posterior distributions for activation energies below ($E_a^{\text{eff}}$) and above ($E_a^{\text{sol}}$) the ERH, and the percentage difference between them $\left(\frac{E_a^{\text{eff}} - E_a^{\text{sol}}}{E_a^{\text{eff}}}\right)$, 4-parameter model version.** Measured concentration fit shown at left, modeled concentration fit shown at right. Distributions are visualized as quantile dotplots [75]; 100 representative dots are shown for each parameter. Black circle below shows posterior median, bars show 68 % (thick) and 95 % (thin) credible intervals.

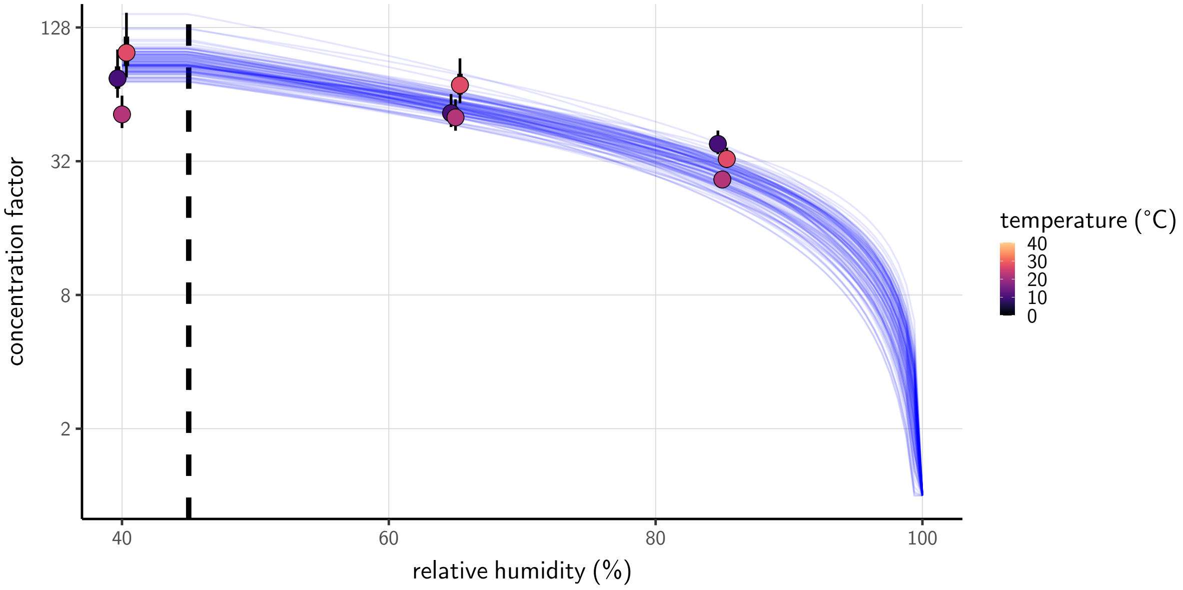

**Figure A9. Concentration factor at quasi-equilibrium a function of relative humidity, modeled concentration fit.** Points show estimates for quasi-equilibrium concentration factor based on empirically measured masses from the evaporation experiments (Fig. A1) and the estimated initial solute mass fraction from the modeled concentration fit (see Main Text, Methods). Estimates shown for each temperature (point color) and ambient RH (x-axis value). Vertical lines around the points show a 70 % (thick) and 100 % (thin) credible interval. Blue curves show model predictions for concentration factor given parameters $\alpha_c$, $\alpha_s$ (Appendix section 4.1, equation 12), and the initial solute mass fraction, all estimated from the measured concentration fit. Each is an independent draw from the joint posterior distribution of the parameters, thus giving a sense of the distribution of possible curves. Vertical dashed line shows the efflorescence relative humidity, 50 %.

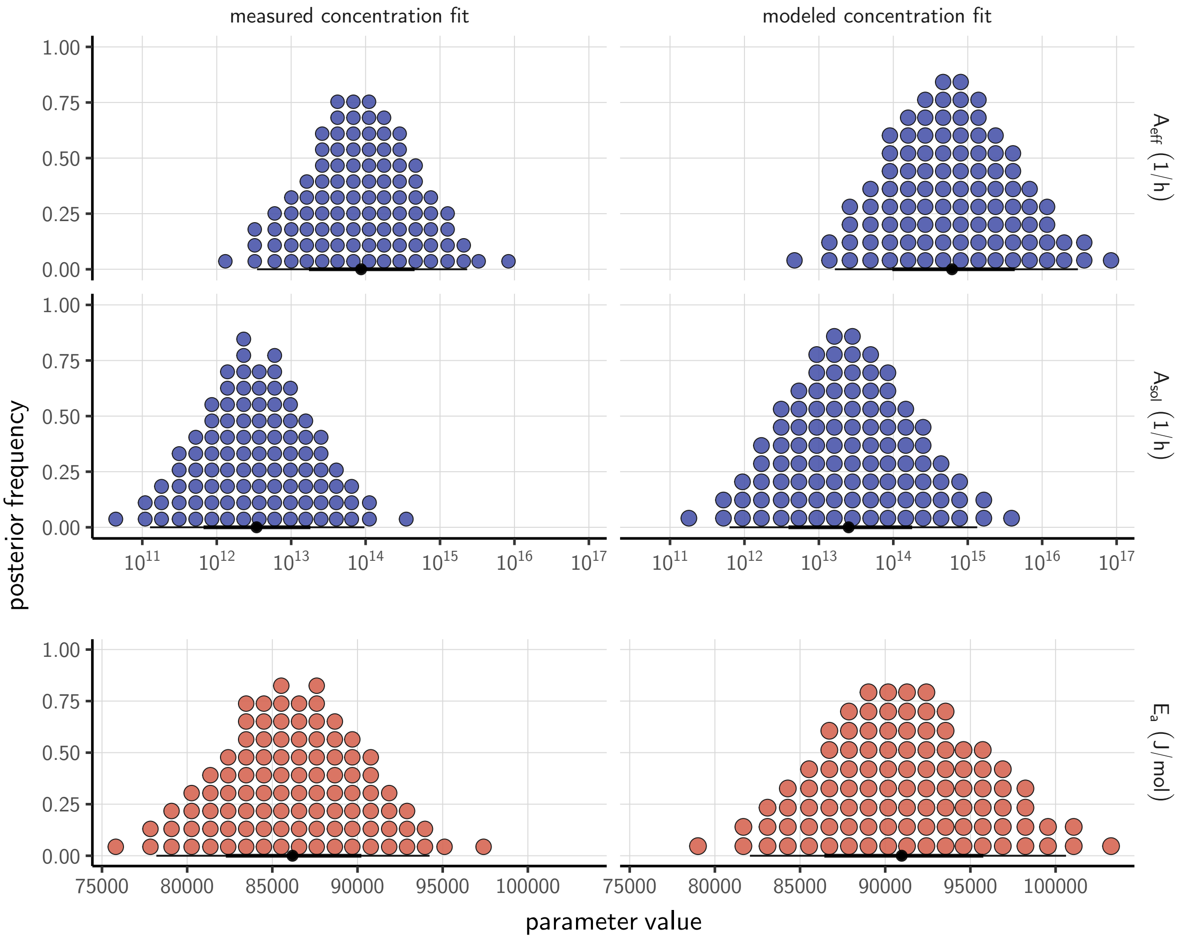

**Figure A10. Posterior distributions for key mechanistic model parameters.** Measured concentration fit shown at left, modeled concentration fit shown at right. Distributions are visualized as quantile dotplots [75]; 100 representative dots are shown for each parameter. Black circle below shows posterior median, bars show 68 % (thick) and 95 % (thin) credible intervals. For $A_{\text{eff}}$ and $A_{\text{sol}}$, parameter values are plotted on a logarithmic scale.

### 2 Key additional tables

**Table A1. Estimated half-lives in hours of SARS-CoV-2 on polypropylene as a function of temperature (T) and relative humidity (RH).** Estimated half-lives are reported as posterior median and the middle 95% credible interval.

| | T (°C) | RH (%) | median half-life (h) | 2.5 % | 97.5 % |
|---|---|---|---|---|---|
| quasi-equilibrium phase | 10 | 40 | 26.55 | 20.28 | 38.75 |
| | 10 | 65 | 14.22 | 12.17 | 17.16 |
| | 10 | 85 | 13.78 | 10.67 | 19.70 |
| | 22 | 40 | 6.43 | 5.52 | 7.56 |
| | 22 | 65 | 2.41 | 2.03 | 2.88 |
| | 22 | 85 | 7.50 | 6.22 | 9.24 |
| | 27 | 40 | 3.43 | 2.91 | 4.12 |
| | 27 | 65 | 1.52 | 1.05 | 2.14 |
| | 27 | 85 | 2.79 | 2.12 | 3.78 |
| evaporation phase | 10 | | 42.08 | 10.97 | 334.34 |
| | 22 | | 12.18 | 4.47 | 163.58 |
| | 27 | | 5.76 | 2.14 | 125.85 |

**Table A2. Estimated half-lives in hours for data from the literature, as a function of material, temperature (T), and relative humidity (RH).** Estimated half-lives are reported as posterior median and the middle 95% credible interval. CCM: cell culture medium; VTM: virus transport medium; Resp. sec.: respiratory secretions

| study | virus | material | T (°C) | RH (%) | median half-life (h) | 2.5 % | 97.5 % |
|---|---|---|---|---|---|---|---|
| Harbourt et al. 2020 [74] | SARS-CoV-2 | Skin | 4 | 45 | $4.17 \times 10^{1}$ | $2.61 \times 10^{1}$ | $1.22 \times 10^{2}$ |
| Lamarre et al. 1989 [77] | HCoV-229E | Bulk CCM | 4 | | $1.97 \times 10^{2}$ | $5.06 \times 10^{1}$ | $1.05 \times 10^{4}$ |
| Rabenau et al. 2005 [82] | SARS-CoV-1 | Bulk CCM | 4 | | 1.24 | $1.18 \times 10^{-1}$ | $9.73 \times 10^{2}$ |
| Lai et al. 2005 [76] | SARS-CoV-1 | Bulk Resp. sec. | 4 | | $4.42 \times 10^{1}$ | $3.68 \times 10^{1}$ | $5.56 \times 10^{1}$ |
| Chin et al. 2020 [13] | SARS-CoV-2 | Bulk VTM | 4 | | $1.98 \times 10^{2}$ | $5.45 \times 10^{1}$ | $9.09 \times 10^{3}$ |
| Van Doremalen et al. 2013 [59] | MERS-CoV | Plastic | 20 | 40 | 1.54 | 1.08 | 2.42 |
| Van Doremalen et al. 2013 [59] | MERS-CoV | Steel | 20 | 40 | 3.14 | 2.26 | 4.71 |
| Lai et al. 2005 [76] | SARS-CoV-1 | Bulk Resp. sec. | 20 | | $1.10 \times 10^{1}$ | 8.45 | $1.61 \times 10^{1}$ |
| Harbourt et al. 2020 [74] | SARS-CoV-2 | Skin | 22 | 45 | 3.77 | 2.10 | 8.17 |
| Lamarre et al. 1989 [77] | HCoV-229E | Bulk CCM | 22 | | $1.51 \times 10^{1}$ | 8.82 | $2.55 \times 10^{1}$ |
| Chin et al. 2020 [13] | SARS-CoV-2 | Bulk VTM | 22 | | $1.83 \times 10^{1}$ | $1.35 \times 10^{1}$ | $2.62 \times 10^{1}$ |
| Van Doremalen et al. 2013 [59] | MERS-CoV | Plastic | 30 | 30 | 1.19 | $6.44 \times 10^{-1}$ | 2.31 |
| Van Doremalen et al. 2013 [59] | MERS-CoV | Steel | 30 | 30 | 1.31 | $7.20 \times 10^{-1}$ | 2.56 |
| Van Doremalen et al. 2013 [59] | MERS-CoV | Plastic | 30 | 80 | $9.73 \times 10^{-1}$ | $5.66 \times 10^{-1}$ | 1.75 |
| Van Doremalen et al. 2013 [59] | MERS-CoV | Steel | 30 | 80 | $5.78 \times 10^{-1}$ | $4.01 \times 10^{-1}$ | $9.96 \times 10^{-1}$ |
| Bucknall et al. 1972 [70] | HCoV-229E | Bulk CCM | 33 | | 1.61 | 1.01 | 4.02 |
| Bucknall et al. 1972 [70] | HCoV-OC43 | Bulk CCM | 33 | | 6.70 | 3.71 | $6.24 \times 10^{1}$ |
| Lamarre et al. 1989 [77] | HCoV-229E | Bulk CCM | 33 | | $1.42 \times 10^{1}$ | 8.68 | $2.27 \times 10^{1}$ |
| Harbourt et al. 2020 [74] | SARS-CoV-2 | Skin | 37 | 45 | $6.05 \times 10^{-1}$ | $3.34 \times 10^{-1}$ | 1.35 |
| Bucknall et al. 1972 [70] | HCoV-229E | Bulk CCM | 37 | | 1.04 | $6.51 \times 10^{-1}$ | 2.06 |
| Bucknall et al. 1972 [70] | HCoV-OC43 | Bulk CCM | 37 | | 4.19 | 2.34 | $4.03 \times 10^{1}$ |
| Lamarre et al. 1989 [77] | HCoV-229E | Bulk CCM | 37 | | 5.77 | 2.84 | $1.10 \times 10^{1}$ |
| Chin et al. 2020 [13] | SARS-CoV-2 | Bulk VTM | 37 | | 2.09 | 1.45 | 3.13 |
| Batéjat et al. 2020 [67] | SARS-CoV-2 | Bulk CCM | 56 | | $2.25 \times 10^{-2}$ | $1.66 \times 10^{-2}$ | $2.78 \times 10^{-2}$ |
| Darnell et al. 2004 [71] | SARS-CoV-1 | Bulk CCM | 56 | | $4.50 \times 10^{-2}$ | $3.41 \times 10^{-2}$ | $6.31 \times 10^{-2}$ |
| Leclercq et al. 2014 [78] | MERS-CoV | Bulk CCM | 56 | | $4.31 \times 10^{-3}$ | $1.29 \times 10^{-3}$ | $1.51 \times 10^{-2}$ |
| Rabenau et al. 2005 [82] | SARS-CoV-1 | Bulk CCM | 56 | | $3.38 \times 10^{-3}$ | $9.53 \times 10^{-6}$ | $2.63 \times 10^{-2}$ |
| Chin et al. 2020 [13] | SARS-CoV-2 | Bulk VTM | 56 | | $1.64 \times 10^{-2}$ | $1.09 \times 10^{-2}$ | $2.89 \times 10^{-2}$ |
| Pagat et al. 2007 [80] | SARS-CoV-1 | Bulk CCM | 60 | | $3.48 \times 10^{-2}$ | $2.59 \times 10^{-2}$ | $5.09 \times 10^{-2}$ |
| Rabenau et al. 2005 [82] | SARS-CoV-1 | Bulk CCM | 60 | | $3.08 \times 10^{-3}$ | $1.18 \times 10^{-5}$ | $2.62 \times 10^{-2}$ |
| Batéjat et al. 2020 [67] | SARS-CoV-2 | Bulk CCM | 65 | | $1.56 \times 10^{-3}$ | $6.84 \times 10^{-6}$ | $1.17 \times 10^{-2}$ |
| Darnell et al. 2004 [71] | SARS-CoV-1 | Bulk CCM | 65 | | $4.13 \times 10^{-2}$ | $3.11 \times 10^{-2}$ | $6.26 \times 10^{-2}$ |
| Leclercq et al. 2014 [78] | MERS-CoV | Bulk CCM | 65 | | $7.34 \times 10^{-4}$ | $5.15 \times 10^{-4}$ | $1.45 \times 10^{-3}$ |
| Batéjat et al. 2020 [67] | SARS-CoV-2 | Bulk Resp. sec. | 65 | | $8.38 \times 10^{-3}$ | $6.01 \times 10^{-3}$ | $1.15 \times 10^{-2}$ |
| Pagat et al. 2007 [80] | SARS-CoV-1 | Bulk CCM | 70 | | $9.82 \times 10^{-3}$ | $7.71 \times 10^{-3}$ | $1.40 \times 10^{-2}$ |
| Chin et al. 2020 [13] | SARS-CoV-2 | Bulk VTM | 70 | | $3.34 \times 10^{-3}$ | $1.66 \times 10^{-3}$ | $6.13 \times 10^{-3}$ |
| Darnell et al. 2004 [71] | SARS-CoV-1 | Bulk CCM | 75 | | $2.24 \times 10^{-3}$ | $9.48 \times 10^{-6}$ | $2.13 \times 10^{-2}$ |
| Batéjat et al. 2020 [67] | SARS-CoV-2 | Bulk Resp. sec. | 95 | | $2.42 \times 10^{-3}$ | $1.61 \times 10^{-3}$ | $3.62 \times 10^{-3}$ |

**Table A3. Parameter estimates for the mechanistic model of SARS-CoV-2 inactivation as a function of temperature and humidity, measured concentration fit.** Estimates are reported as posterior median and the middle 95% credible interval.

| parameter | median | 2.5 % | 97.5 % | unit |
|---|---|---|---|---|
| $A_{\mathrm{eff}}$ | $8.70 \times 10^{13}$ | $3.48 \times 10^{12}$ | $2.31 \times 10^{15}$ | $\mathrm{h}^{-1}$ |
| $A_{\mathrm{sol}}$ | $3.43 \times 10^{12}$ | $1.27 \times 10^{11}$ | $9.64 \times 10^{13}$ | $\mathrm{h}^{-1}$ |
| $E_a$ | $8.62 \times 10^{4}$ | $7.82 \times 10^{4}$ | $9.42 \times 10^{4}$ | $\mathrm{J\,mol}^{-1}$ |

**Table A4. Parameter estimates for the mechanistic model of SARS-CoV-2 inactivation as a function of temperature and humidity, modeled concentration fit.** Estimates are reported as posterior median and the middle 95% credible interval.

| parameter | median | 2.5 % | 97.5 % | unit |
|---|---|---|---|---|
| $A_{\mathrm{eff}}$ | $6.15 \times 10^{14}$ | $1.64 \times 10^{13}$ | $3.02 \times 10^{16}$ | $\mathrm{h}^{-1}$ |
| $A_{\mathrm{sol}}$ | $2.51 \times 10^{13}$ | $6.31 \times 10^{11}$ | $1.34 \times 10^{15}$ | $\mathrm{h}^{-1}$ |
| $E_a$ | $9.10 \times 10^{4}$ | $8.21 \times 10^{4}$ | $1.01 \times 10^{5}$ | $\mathrm{J\,mol}^{-1}$ |

### 3 Mechanistic inactivation model interpretation

#### 3.1 Interpretation of the single activation energy

We observe in the main text that a single activation energy $E_a$ explains the data well across the effloresced and solution regimes (Fig. A8).

Moreover, our estimate is consistent with activation energies observed for other RNA viruses [85]. Our median [95 % credible interval] measured concentration fit $E_a$ estimate, 86 182.88 J mol$^{-1}$ [78 190.02, 94 229.79], falls squarely within the range of literature estimates (approximately 60 000 to 240 000 J mol$^{-1}$) [85].

These observations raise the question of whether the actual inactivating reaction is identical in the effloresced and solution regimes, in different media, and for different viruses. But at least for a given virus or family of viruses, it is possible for virus inactivation reactions to have the same activation energy even if different media or different environments imply a different inactivating reactant. Plausible routes of chemical virus inactivation include conformational changes in virion proteins and disruption of the virus capsid [62]. These may occur via a two-step reaction:

$$\text{viable virion} + \text{external reactant} \leftrightarrow \text{intermediate product} \longrightarrow \text{inactivated virion} \tag{4}$$

If the second step is rate-limiting, then the overall reaction kinetics are first order and the measured activation energy will reflect the $E_a$ for that step. This energy could easily depend only on the virus proteins and not on the external reactant.

#### 3.2 Two-step reactions can produce first-order kinetics proportional to concentration

Provided the external reactant concentration $[r]$ is not meaningfully depleted, even a two step inactivation reaction of this form would still imply a linear dependence of inactivation rate on concentration of external reactant, and thus a linear dependence on solution concentration as postulated in our model (equation 3). Below we describe a minimal two-step reaction mechanism that is consistent with these observations.

We denote the concentration of viable virus by $[v_v]$, the concentration of inactivated virus by

$[v_i]$, and the concentration of intermediate product by $[x]$. We denote the rate constants for the forward and backward first-step reactions by $k_1^+$ and $k_1^-$ and the rate constant for the second-step reaction by $k_2^+$. We have:

$$\begin{aligned}
\frac{\mathrm{d}[v_v]}{\mathrm{d}t} &= -k_1^+[r][v_v] + k_1^-[x]\,\mathrm{d}t \\
\frac{\mathrm{d}[x]}{\mathrm{d}t} &= k_1^+[r][v_v] - k_1^-[x] - k_2^+[x]\,\mathrm{d}t \\
\frac{\mathrm{d}[v_i]}{\mathrm{d}t} &= k_2^+[x]\,\mathrm{d}t
\end{aligned} \tag{5}$$

By assumption, the first step in the reaction is fast relative to the second. The intermediate product $[x]$ should therefore reach a quasi-equilibrium value $[\bar{x}]$. We solve for it by setting $\frac{\mathrm{d}[x]}{\mathrm{d}t} = 0$ and neglecting the smaller $-k_2^+$ term:

$$[\bar{x}] = [r]\frac{k_1^+}{k_1^-}[v_v] \tag{6}$$

Substituting $[\bar{x}]$ for $[x]$ into $\frac{\mathrm{d}[v_i]}{\mathrm{d}t}$, it follows that virus inactivation obeys first-order kinetics proportional to the external reactant concentration $[r]$:

$$\frac{\mathrm{d}[v_i]}{\mathrm{d}t} = [r]k_2^+\frac{k_1^+}{k_1^-}[v_v]\,\mathrm{d}t \tag{7}$$

#### 3.3 Interpretation of the asymptotic reaction rates

We also observe that the pre-exponential factor (asymptotic high temperature reaction rate) is somewhat but not substantially greater in the effloresced regime than in the solution regime ($A_{\mathrm{eff}} > A_{\mathrm{sol}}$). Since $A_{\mathrm{sol}}$ is modulated by $\frac{[S_{\mathrm{eq}}]}{[S_0]}$, this implies that reaction rates in the effloresced crystals (which we assume occur at the same rate for all sub-ERH ambient humidities) are faster than reactions at 100 % RH, but not as fast as at humidities slightly above the ERH, such as 65 % (Figs. A10).

This empirical result is plausible. Below the ERH, reactants are in closer proximity, but also less mobile: modeled as a quasi-solution, there is a higher reactant concentration but also a lower diffusion coefficient. It is thus plausible that the effective rate of potentially reactive collisions for a given temperature could be greater than the rate in dilute solution at 100 % RH, but substantially lower than the rate in more concentrated solution at 65 % RH.

### 4 Mechanistic modeling of evaporation and concentration

To measure the solute concentration factor over time and to determine when droplets reached evaporative quasi-equilibrium (i.e., evaporation became slow enough that concentration factor could be treated as a constant), we quantified the evaporation of the suspension medium on polypropylene plastic (without virus) at the tested temperature and humidity combinations (Main Text Methods; Fig. A1).

To extrapolate to unobserved relative humidities, we estimated the quasi-equilibrium solute concentration factor $\frac{[S_{eq}]}{[S_0]}$ as a function of relative humidity $h$.

#### 4.1 Solute concentration factor

The concentration factor as a function of time $\frac{[S(t)]}{[S_0]}$ is equal to the ratio of the initial mass of water $w(0)$ (before evaporation begins) to the current mass of water $w(t)$. We measured total masses $m(t)$, not masses of water, but assuming that the mass of solutes, $s$, is conserved:

$$\frac{[S(t)]}{[S_0]} = \frac{w(0)}{w(t)} = \frac{m(0) - s}{m(t) - s} \tag{8}$$

and so:

$$\frac{[S_{eq}]}{[S_0]} = \frac{w(0)}{w(\infty)} = \frac{m(0) - s}{m(\infty) - s} \tag{9}$$

In order to predict decay rates at unobserved relative humidities, we fit a semi-mechanistic function to the measured concentration factors to predict $\frac{[S_{eq}]}{[S_0]}$ as a function of fractional relative humidity $h$. We begin with the observation (see section 4.5 for a derivation) that if $X(\infty)$ is the molar fraction of solutes in the solution at quasi-equilibrium and $X(0)$ is the initial molar fraction of solutes, then:

$$\frac{[S_{\mathrm{eq}}]}{[S_0]} = \frac{1 - X(0)}{X(0)} \frac{X(\infty)}{1 - X(\infty)} \tag{10}$$

We denote the initial ratio of the molar fractions by $r(0) = \frac{X(0)}{1-X(0)}$.

The final molar ratio $r(\infty) = \frac{X(\infty)}{1-X(\infty)}$ depends on the fractional relative humidity $h$. We approximate this relationship by a flexible two-parameter function:

$$r(\infty) = \frac{X(\infty)}{1 - X(\infty)} = \left(\frac{-\ln(h)}{\alpha_s}\right)^{\frac{1}{\alpha_c}} \tag{11}$$

Combining yields:

$$\frac{[S_{\mathrm{eq}}]}{[S_0]} = \frac{1}{r(0)} \left(\frac{-\ln(h)}{\alpha_s}\right)^{\frac{1}{\alpha_c}} \tag{12}$$

The estimated parameters $\alpha_c, \alpha_s > 0$ reflect deviations of our solute mixture from ideal behavior ($\alpha_c = \alpha_s = 1$). We derived this approximate expression from chemical theory; see section 4.6 for the derivation. Ideal chemical behavior would imply a linear relationship between $X(\infty)$ and $h$ near $h = 1$: $X(\infty) = 1 - h$. This works well for dilute solutions. But it predicts too high a concentration factor at low relative humidities, since it neglects the increasingly strong effects of solutes in preventing evaporation as those solutes become more concentrated. To extrapolate in a worthwhile way, then, we need at least a minimal model of non-ideal evaporative behavior in a concentrated solution. Our simple function fits our data well (Fig. 12).

#### 4.2 Modeling of medium evaporation

In our evaporation experiments, we observed an approximately linear decrease in water mass $w(t)$ over time (Fig. A1), followed by a leveling off at an approximately constant value (quasi-equilibrium). We therefore approximated the evaporation process with a piece-wise linear function:

$$m(t) = \begin{cases} m(0) - \beta t & \beta t < m(0) - m(\infty) \\ m(\infty) & \text{otherwise} \end{cases} \tag{13}$$

As noted in section 4.1, we assumed that solute mass $s$ was conserved, so $w(t) = m(t) - s$. It follows that:

$$w(t) = \begin{cases} m(0) - s - \beta t & \beta t < m(0) - m(\infty) \\ m(\infty) - s & \text{otherwise} \end{cases} \tag{14}$$

This implies that the concentration factor as a function of time is given by:

$$\frac{[S(t)]}{[S_0]} = \frac{w(0)}{w(t)} = \frac{m(0) - s}{m(0) - s - \beta t} \tag{15}$$

Defining $B = \frac{\beta}{w(0)} = \frac{\beta}{m(0)-s}$ yields a normalized form:

$$\frac{[S(t)]}{[S_0]} = \frac{1}{1 - Bt} \tag{16}$$

#### 4.3 Evaporation and quasi-equilibrium phases

In our estimation models, we partitioned virus inactivation into two phases: evaporation and quasi-equilibrium (see Methods). We denote the time to quasi-equilibrium for experiment $i$ by $\tau_i$.

We determined $\tau_i$ for each inactivation experimental condition based on on the evaporative mass loss rate $\beta_i$ in the corresponding evaporation experiment.

For the simple regression model and the measured concentration fit of the mechanistic model, we defined $\tau_i$ as the time to reach the measured final total mass $m_i(\infty)$ from the measured initial total mass $m_i(0)$ given the inferred evaporative mass loss rate $\beta_i$:

$$\tau_i = \frac{m_i(0) - m_i(\infty)}{\beta_i} \tag{17}$$

For the modeled concentration fit of the mechanistic model, we partition the phases not based on $\tau_i$ but rather based on the time $\bar{\tau}_i$ to reach the predicted quasi-equilibrium concentration factor $\frac{[S_{\mathrm{eq}}]}{[S_0]}_i$ given the inferred $B_i = \frac{\beta_i}{w_i(0)}$:

$$\bar{\tau}_i = \frac{1 - \frac{1}{\frac{[S_{\mathrm{eq}}]}{[S_0]}_i}}{B_i} \tag{18}$$

Note that this relation also holds for the measured concentration version. Letting $\frac{[S_{\mathrm{eq}}]}{[S_0]} = \frac{m(0)-s}{m(\infty)-s}$, equation 18 simplifies to equation 17.

#### 4.4 Modeling of virus decay dynamics during the evaporation phase

Prior to evaporative quasi-equilibrium or complete efflorescence, virions are in wet conditions, with non-negligible evaporation ongoing. The degree of concentration of that solution $\frac{[S(t)]}{[S_0]}$ changes as a function of time as the solvent (here, suspension medium) evaporates, until a quasi-equilibrium state is reached at $\frac{[S(t)]}{[S_0]} = \frac{[S_{\mathrm{eq}}]}{[S_0]}$.

Per equation 8, the concentration factor as a function of time is equal to $\frac{w(0)}{w(t)}$.

The inactivation rate during that evaporation phase, which we denote by $k_{\mathrm{ev}}$, is then a function of time $k_{\mathrm{ev}}(t)$:

$$k_{\mathrm{ev}}(t) = \frac{w(0)}{w(t)} A_{\mathrm{sol}} \exp\left(-\frac{E_a}{RT}\right) \tag{19}$$

Letting $v(t)$ denote the quantity of viable virus, inactivation kinetics will then proceed according to the differential equation:

$$\frac{\mathrm{d}v}{\mathrm{d}t} = -k_{\mathrm{ev}}(t) v \,\mathrm{d}t \tag{20}$$

We define $k_0 = k_{\mathrm{sol}}(0) = A_{\mathrm{sol}} \exp\left(-\frac{E_a}{RT}\right)$ and apply our linear evaporation model from equation

15:

$$\frac{\mathrm{d}v}{\mathrm{d}t} = -k_{\mathrm{ev}}(t)v\,\mathrm{d}t = -\frac{k_0}{1 - \left(\frac{\beta}{w(0)}\right)t} v\,\mathrm{d}t = -\frac{k_0}{1 - Bt} v\,\mathrm{d}t \tag{21}$$

Solving yields:

$$v(t) = v(0)\exp\left(\frac{k_0}{B}\ln(1 - Bt)\right) = v(0)(1 - Bt)^{(k_0/B)} \tag{22}$$

subject to the constraint that $Bt < 1$, which is always satisfied for $t \leq \tau$, under the assumption that some non-zero amount of water remains at quasi-equilibrium.

Since virus titers are typically measured in $\log_{10}$ units, it is useful to have this expression in those terms:

$$\log_{10} v(t) = \log_{10}(v_0) + \frac{k_0}{B}\log_{10}(1 - Bt) \tag{23}$$

#### 4.5 Relationship between concentration factor and solute molar fraction (equation 10)

Under the assumption that mass of solute does not change, all mass change reflects loss or gain of solvent. This mass change translates directly into increased or decreased concentration, and allows us to compute the estimated concentration factor as a function of time, $\frac{[S(t)]}{[S_0]}$, based on our evaporation experiments.

If we have $N_w(t)$ moles of solvent versus an initial value of $N_w(0)$ and a constant number $N_s$ of solute, then following a similar reasoning as in equation 8:

$$\frac{[S(t)]}{[S_0]} = \frac{N_w(0)}{N_w(t)} \tag{24}$$

If $X(t)$ is the mole fraction of solutes in the solution, $N_w(t) = (1 - X(t))N(t)$ and $N_s = X(t)N(t)$

where $N(t) = N_w(t) + N_s$. It follows that the ratio of moles is the ratio of the mole fractions:

$$\frac{N_w(t)}{N_s} = \frac{1 - X(t)}{X(t)} \tag{25}$$

Since $N_s$ does not change:

$$\frac{N_w(0)}{N_s} = \frac{(1 - X(0))}{X(0)} \tag{26}$$

Hence:

$$\frac{[S(t)]}{[S_0]} = \frac{N_w(0)}{N_w(t)} = \frac{N_w(0)/N_s}{N_w(t)/N_s} = \frac{\frac{1-X(0)}{X(0)}}{\frac{1-X(t)}{X(t)}} = \left(\frac{1 - X(0)}{X(0)}\right) \frac{X(t)}{1 - X(t)} \tag{27}$$

and therefore:

$$\frac{[S_{\text{eq}}]}{[S_0]} = \left(\frac{1 - X(0)}{X(0)}\right) \frac{X(\infty)}{1 - X(\infty)} \tag{28}$$

#### 4.6 Derivation of approximate functional form for the quasi-equilibrium solute concentration (equation 11)

To compute $\frac{[S_{\text{eq}}]}{[S_0]}$ as a function of fractional relative humidity $h$, we need an expression for the ratio of the quasi-equilibrium solute mole fraction $X(\infty)$ to the quasi-equilibrium solvent mole fraction $1 - X(\infty)$ as a function of $h$.

An evaporating aqueous solution reaches equilibrium with the ambient air when the ambient relative humidity is equal to the water activity $a_w$ in the solution:

$$h = a_w \tag{29}$$

For an ideal solution, the water activity would be given by:

$$a_w = 1 - X(\infty) \tag{30}$$

where $X(\infty)$ is the mole fraction of solutes (Raoult's law). In a real solution, this expression must be modified to account for non-ideal behavior.

If there are $n$ species of solute ions and/or molecules present with molar fractions $X_j$, we express this non-ideality in terms of the practical osmotic coefficient $\phi(X_1, ...X_n)$ [68], which is in general a function of the $X_j$:

$$a_w = \exp\left(-\phi \frac{\sum_{j=1}^{n} X_j}{1 - \sum_{j=1}^{n} X_j}\right) = \exp\left(-\phi \frac{X(\infty)}{1 - X(\infty)}\right) \tag{31}$$

Since our medium has a consistent solute formulation and we assume that solutes are conserved, we can treat $\phi$ as a function of the total solute molar fraction $X(t)$. We use the following flexible functional form for $\phi$:

$$\phi = \alpha_s \left(\frac{X}{1 - X}\right)^{\alpha_c - 1} \tag{32}$$

With $\alpha_c, \alpha_s > 0$. We define these constrained parameters in terms of unconstrained parameters $c_c$ and $c_s$:

$$\begin{aligned} \alpha_c &= \exp(-c_c) \\ \alpha_s &= \exp(-c_s) \end{aligned} \tag{33}$$

It follows that:

$$a_w = \exp\left[-\alpha_s \left(\frac{X}{1 - X}\right)^{\alpha_c}\right] \tag{34}$$

This is a flexible two-parameter function with a number of desirable properties.

- $X = 0$ implies $a_w = 1$ and $X = 1$ implies $a_w = 0$, as should be the case.
- When $c_c, c_s = 0$, the relationship approximates the linear behavior observed in the ideal case, and we have $\phi = 1$ regardless of $X$, reflecting this ideality.
- $c_c < 0$ implies a concave relationship between mole fraction and activity near $a_w = 1$, $c_c > 0$ implies a convex relationship there, and $c_c = 0$ a linear relationship.

- Varying $c_s$ controls the steepness of the relationship near $a_w = 1$ while preserving concavity in that region; larger values imply a steeper relationship.
- Empirical $\phi(X)$ functions for important solute components of DMEM, such as NaCl, are monotonically increasing in $X$ over the range of expected equilibrium mole fractions [44], and thus should be readily approximated by our function.

Using the property that evaporative equilibrium occurs when $a_w = h$, we approximate the ratio $r(\infty)$ of solute to solvent mole fractions at quasi-equilibrium 11) by:

$$r(\infty) = \frac{X(\infty)}{1 - X(\infty)} = \left(\frac{-\ln(h)}{\alpha_s}\right)^{\frac{1}{\alpha_c}}$$

This function readily approximates a number of realistic shapes [44, 83] for the relationship between $X$ and $h$, particularly on the interval of interest, between 100% relative humidity and the efflorescence relative humidity (ERH) ($1 \geq h \geq \text{ERH} \approx 0.45$).

This function has simpler approximations to the humidity-molar-ratio relationship as special cases. For instance, $\alpha_c = 1$ implies that $\phi$ does not vary with solute mole fraction $X$ (as happens in ideal solutions).

The main downside of this function is that our $\phi(X)$ is constrained to be be monotonic. It is thus impossible for the relationship between $h$ and $\frac{X}{1-X}$ to have more than one concavity change the range $[0, 1]$. But this is unlikely to be important given that we are mainly interested (and fitting to) the range from the ERH to 100 % relative humidity. In fact, an always-concave function readily explains our evaporation data in that range (Fig. A9).

Plugging equation 11 into equation 10 yields the expression for $\frac{[S_{\text{eq}}]}{[S_0]}$ in terms of the initial solute mole fraction ratio $r(0) = \frac{X(0)}{1-X(0)}$ and the ambient relative humidity $h$ given in equation 12:

$$\frac{[S_{\text{eq}}]}{[S_0]} = \frac{1}{r(0)}\left(\frac{-\ln(h)}{\alpha_s}\right)^{\frac{1}{\alpha_c}}$$

Notice that while quasi-equilibrium concentration factors will depend on both $\alpha_c$ and $\alpha_s$, the ratio of two quasi-equilibrium concentration factors from the same baseline (i.e. $\frac{[S^a_{\text{eq}}]/[S_0]}{[S^b_{\text{eq}}]/[S_0]}$ for two different ambient humiditites $h_a$ and $h_b$) will depend only on $\alpha_c$:

$$\frac{[S_{\text{eq}}^a]/[S_0]}{[S_{\text{eq}}^b]/[S_0]} = \left(\frac{\ln(h_a)}{\ln(h_b)}\right)^{\frac{1}{\alpha_c}} \tag{35}$$

Using equation 35 in conjunction with equation 3, one can predict a half-life at one temperature-relative humidity pair from a half-life measured at another, provided all else is equal. We use such an approach to make relative predictions in our meta-analysis (Fig. 3d). See section 6.4.2 for details.

### 5 Bayesian estimation models

#### 5.1 Model notation

In the model notation that follows, the symbol ∼ denotes that a random variable is distributed according to the given distribution. Normal distributions are parametrized as:

$$\text{Normal}(\text{mean, standard deviation})$$

Positive-constrained normal distributions ("Half-Normal") are parametrized as:

$$\text{Half-Normal}(\text{mode, standard deviation})$$

For each inactivation experiment (set of temperature humidity conditions for a given virus), there is a corresponding medium evaporation experiment, which in which we measured the evaporation of suspension medium at that same temperature and humidity.

#### 5.2 Titer inference

##### 5.2.1 Titer inference model

We inferred individual titers directly from titration well data using a Poisson single-hit model.

We then modeled individual positive and negative wells for sample $i$ according to a Poisson single-hit model [69]. That is, the number of virions that successfully infect cells within a given well is Poisson distributed with mean:

$$\ln(2)10^{v_i} \tag{36}$$

This expression for the mean derives from the fact that our units are $\text{TCID}_{50}$; the probability of a positive well at $v_i = 0$, i.e. 1 $\text{TCID}_{50}$, is equal to $1 - \exp(-\ln(2) \times 1) = 0.5$.

Let $y_{idk}$ be a binary variable indicating whether the $k^{\text{th}}$ well at dilution factor $d$ (where $d$ is expressed as $\log_{10}$ dilution factor) for sample $i$ was positive (so $y_{idk} = 1$ if that well was positive and 0 if it was negative). Under a single-hit process, a well will be positive as long as at least one virion successfully infects a cell.

It follows from equation 36 that the conditional probability of observing $y_{idk} = 1$ given a true underlying $\log_{10}$ titer $v_i$ is given by:

$$\mathcal{L}(y_{idk} = 1 \mid v_i) = 1 - \exp\left(-\ln(2) \times 10^{(v_i - d)}\right) \tag{37}$$

This is simply the probability that a Poisson random variable with mean $\ln(2)10^{(v_i-d)}$ is greater than 0, and $v_i - d$ is the expected concentration of virions, measured in $\log_{10}\text{TCID}_{50}$, in the dilute sample. Similarly, the conditional probability of observing $y_{idk} = 0$ given a true underlying $\log_{10}$ titer $v_i$ is:

$$\mathcal{L}(y_{idk} = 0 \mid v_i) = \exp\left(-\ln(2) \times 10^{(v_i - d)}\right) \tag{38}$$

which is the probability that the Poisson random variable is equal to 0.

This gives us our likelihood function, assuming independence of outcomes across wells. Titrated doses introduced to each cell-culture well were of volume 0.1 mL, so we incremented inferred titers by 1 to convert to units of $\log_{10}\text{TCID}_{50}/\text{mL}$.

##### 5.2.2 Titer inference model prior distributions

We assigned a weakly informative Normal prior to the $\log_{10}$ titers $v_i$ ($v_i$ is the titer for sample $i$ measured in $\log_{10}\text{TCID}_{50}/0.1\text{mL}$, since wells were inoculated with 0.1mL), similar to that used in our previous work [18]:

$$v_i \sim \text{Normal}(2.5, 4) \tag{39}$$

###### 5.2.3 Titer inference model predictive checks

We assessed the appropriateness of this prior distribution choice using prior predictive checks. The prior checks suggested that prior distributions were agnostic over the titer values of interest (Figs. A11, A12):

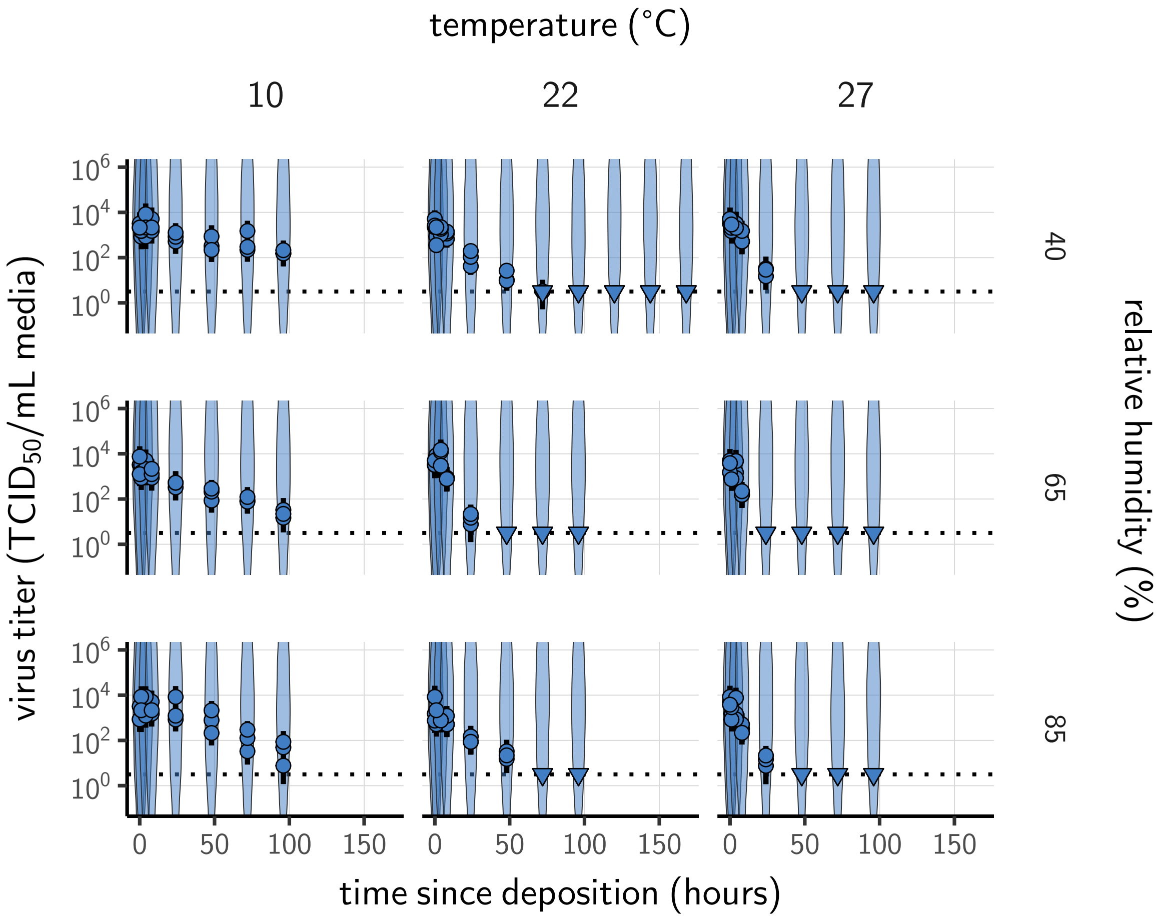

**Figure A11. Prior predictive check for SARS-CoV-2 titer inference.** Violin plots show distribution of simulated titers sampled from the prior predictive distribution. Points show posterior median estimated titers in $\log_{10}$TCID$_{50}$/mL for each sample; lines show 95 % credible intervals. Time-points with no positive wells for any replicate are plotted as triangles at the approximate single-replicate limit of detection (LOD) of the assay—denoted by a black dotted line at $10^{0.5}$ TCID$_{50}$/mL media—to indicate that a range of sub-LOD values are plausible. Three samples collected at each time-point. x-axis shows time since sample deposition. Wide coverage of violins relative to datapoints shows that priors are agnostic over the titer values of interest.

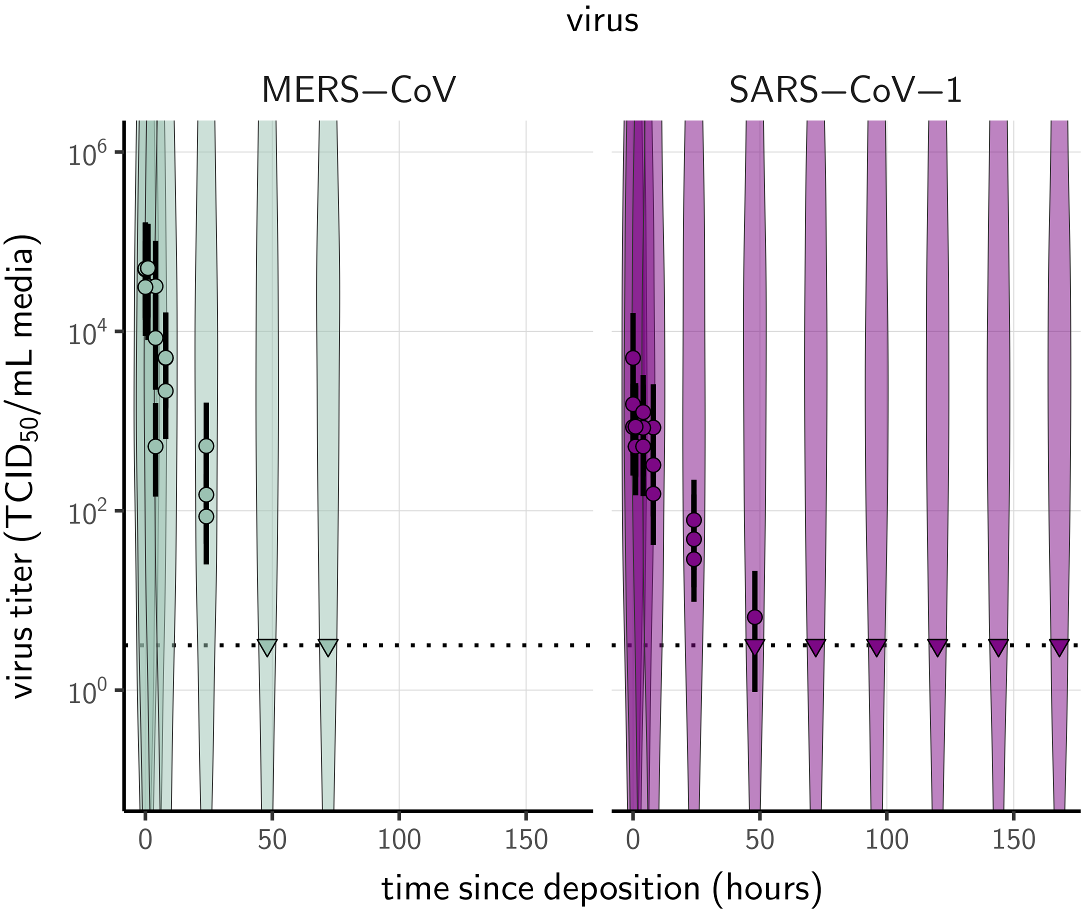

**Figure A12. Prior predictive check for titer inference for SARS-CoV-1 and MERS-CoV.** Violin plots show distribution of simulated titers sampled from the prior predictive distribution. Points show posterior median estimated titers in $\log_{10}\text{TCID}_{50}/\text{mL}$ for each sample; lines show 95 % credible intervals. Time-points with no positive wells for any replicate are plotted as triangles at the approximate single-replicate limit of detection (LOD) of the assay—denoted by a black dotted line at $10^{0.5}$ $\text{TCID}_{50}/\text{mL}$ media—to indicate that a range of sub-LOD values are plausible. Three samples collected at each time-point. x-axis shows time since sample deposition. Wide coverage of violins relative to datapoints shows that priors are agnostic over the titer values of interest.

#### 5.3 Evaporation model

Following 4.2, equation 13, we modeled the expected mass $\bar{m}_i(t)$ for each evaporation experiment $i$ according to the equation:

$$\bar{m}_i(t) = \begin{cases} m_i(0) - \beta_i t & \beta_i t < m_i(0) - m_i(\infty) \\ m_i(\infty) & \text{otherwise} \end{cases} \tag{40}$$

We modeled that the observed masses $m_i(t)$ as normally distributed about the predicted masses $\bar{m}_i(t)$ with an estimated, experiment-specific standard deviation $\sigma_{ei}$:

$$m_i(t) \sim \text{Normal}(\bar{m}_i(t), \sigma_{ei}) \tag{41}$$

To make evaporation prior distributions more interpretable, we placed our prior not on the evaporative mass loss rate $\beta_i$ but rather on the time to reach quasi-equilibrium $\tau_i$ which is related to $\beta_i$ by equation 17:

$$\tau_i = \frac{m_i(0) - m_i(\infty)}{\beta_i}$$

We placed weakly informative Half-Normal priors on the times to quasi-equilibrium $\tau_i$ (measured in hours) and on the measurement standard deviations $\sigma_{ei}$:

$$\tau_i \sim \text{Half-Normal}(10, 10) \tag{42}$$

$$\sigma_{ei} \sim \text{Half-Normal}(0, 1) \tag{43}$$

#### 5.4 Empirical virus decay

##### 5.4.1 Simple regression model

The duration of virus detectability depends not only on environmental conditions and treatment method but also initial inoculum and sampling noise. We therefore estimated the exponential decay rates of viable virus (and thus virus half-lives) using a simple Bayesian regression approach analogous to that described in Fischer et al. 2020 [18]. This modeling approach allowed us to account for differences in initial inoculum levels across samples as well as other sources of experimental noise. The model yields estimates of posterior distributions of viral decay rates and half-lives in the various experimental conditions – that is, estimates of the range of plausible values for these parameters given our data, with an estimate of the overall uncertainty [73].

Our data consist of nine different experimental conditions corresponding to the combinations of three temperatures (10 °C, 22 °C, and 27 °C) and three relative humidity levels (40 %, 65 %, and 85 %). For each treatment, three samples were collected at 0, 1, 4, 8, 24, 72 and 96 hours after deposition. We also used this model for our group's SARS-CoV-1 and MERS-CoV data (in the meta-analysis), which had one experimental condition each: 22 °C and 40 %RH, observed over multiple timepoints. We accounted for evaporation with the same 22 °C, 40 % RH suspension medium evaporation data used for SARS-CoV-2 at that temperature and humidity (as all the virus inactivation experiments were conducted using the same medium).

We modeled each sample $j$ for experimental condition $i$ as starting with some true initial $\log_{10}$ titer $v_{ij0}$. At the time $t_{ij}$ that it is sampled, it has titer $v_{ij}$. As described in section 4.3, we partitioned each experiment $i$ into a evaporation phase and a quasi-equilibrium phase according to an estimated quasi-equilibration time $\tau_i$.

We modeled loss of viable virus at quasi-equilibrium as exponential decay at an experiment-specific rate $\lambda_i$. To avoid making assumptions about the correctness of our evaporation phase inactivation model from section 4.4, we approximated loss of viable virus during the evaporation phase as exponential decay with one decay rate for each temperature condition (which applies to all associated humidity conditions). That is, the evaporation phase decay rate for experiment $i$ is $l_{T(i)}$, where $T(i)$ denotes the temperature for experiment $i$.

It follows that the quantity $v_{ij}$ of virus sampled at time $t_{ij}$ is given by:

$$v_{ij} = \begin{cases} v_{ij0} - l_{T(i)} t_{ij} & t_{ij} \leq \tau_i \\ v_{ij0} - l_{T(i)} \tau_i - \lambda_i (t_{ij} - \tau_i) & t_{ij} > \tau_i \end{cases} \tag{44}$$

We used the direct-from-well data likelihood function described above, except that instead of estimating individual titers independently, we estimated $\lambda_i$ and $l_{T(i)}$ under the assumption that our observed well data $y_{idk}$ reflected the corresponding predicted titers $v_{ij}$.

To check the robustness of our results to our assumptions about the evaporation phase, we also fit a model only to the quasi-equilibrium phase data, with time measured since quasi-equilibrium was reached. In that model, the intercepts $v_{ij0}$ thus reflect the estimated titer at the time quasi-equilibrium was reached:

$$v_{ij0} - \lambda_i(t_{ij} - \tau_i) \tag{45}$$

We modeled each experiment $i$ as having a mean initial $\log_{10}$ titer $\bar{v}_{i0}$. We modeled the individual $v_{ij0}$ as normally distributed about $\bar{v}_{i0}$ with an estimated, experiment-specific standard deviation $\sigma_i$:

$$v_{ij0} \sim \text{Normal}(\bar{v}_{i0}, \sigma_i) \tag{46}$$

##### 5.4.2 Simple regression model prior distributions

We placed a Normal prior on the mean initial $\log_{10}$ titers $\bar{v}_{i0}$ to reflect the known inocula, similar to.

$$\bar{v}_{i0} \sim \text{Normal}(2.5, 1) \tag{47}$$

We placed a Half-Normal prior on the standard deviations $\sigma_i$:

$$\sigma_i \sim \text{Half-Normal}(0, 0.5) \tag{48}$$

The allows either for large variation (1 log) about the experiment mean or for substantially less variation, depending on the data. It is similar—though slightly more diffuse—to that used in prior work [21].

To encode prior information about the decay rates in an interpretable way, we placed Normal priors on the log half-lives $\ln(\eta_i)$, where $\eta_i = \frac{\log_{10}(2)}{\lambda_i}$ and $\ln(\theta_{T(i)})$, where $\theta_{T(i)} = \frac{\log_{10}(2)}{l_{T(i)}}$. We made the priors weakly informative (diffuse over the biologically plausible half-lives); we verified this with prior predictive checks.

$$\begin{aligned} \ln(\eta_i) &\sim \text{Normal}(\ln(6), 2) \\ \ln\left(\theta_{T(i)}\right) &\sim \text{Normal}(\ln(24), 1.25) \end{aligned} \tag{49}$$

We used a larger prior mean for the evaporation phase decay rate based on observations of slow decay of SARS-CoV-2 at moderate temperatures in bulk medium [13] and similar results for other viruses [40].

##### 5.4.3 Simple regression model predictive checks

We assessed the appropriateness of prior distribution choices using prior predictive checks and assessed goodness of fit for the estimated model using posterior predictive checks. Prior checks suggested that prior distributions were agnostic over the parameter values of interest, and posterior checks suggested a good fit of the model to the data. The resultant checks are shown below (Fig. A13–A18).

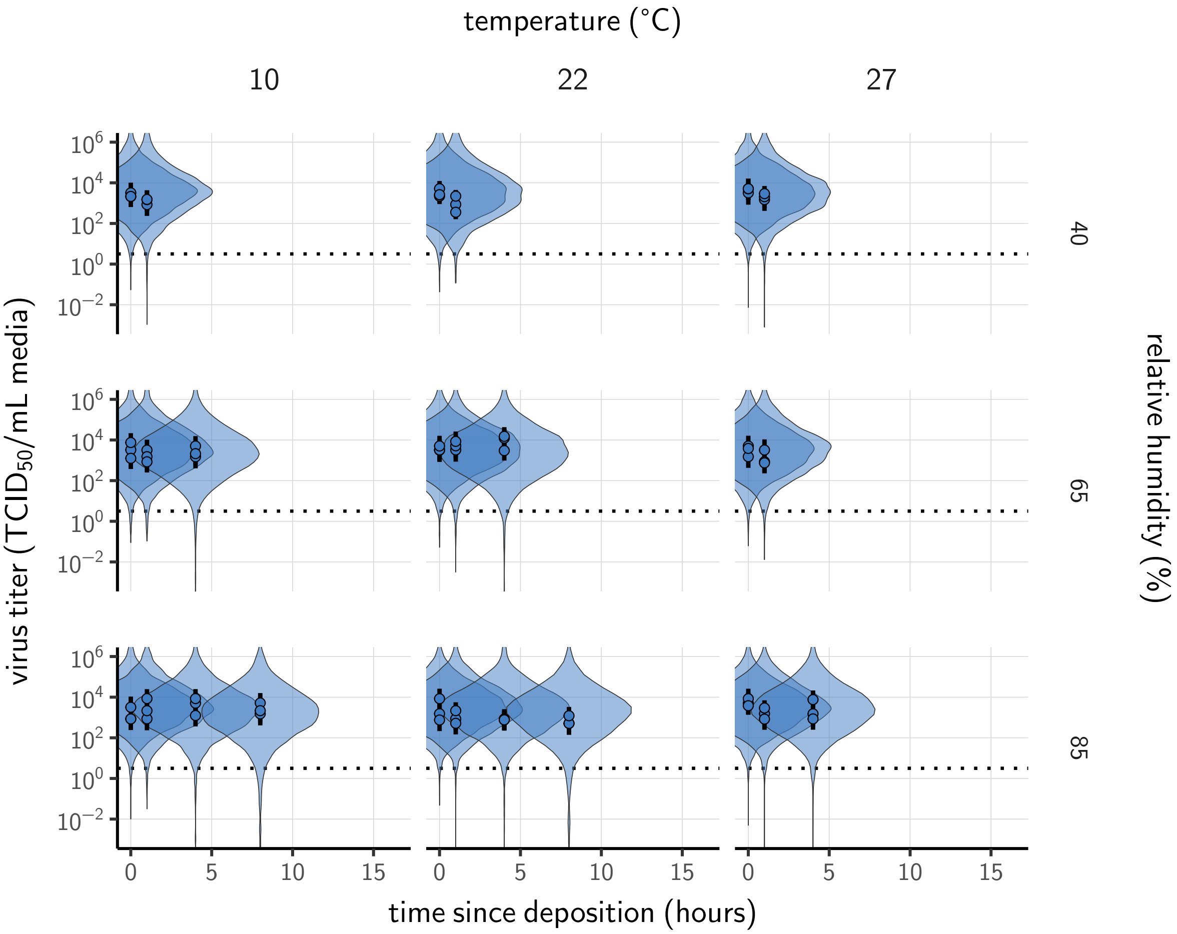

**Figure A13. Prior predictive check for empirical virus decay during the evaporation phase for SARS-CoV-2.** Violin plots show distribution of simulated titers sampled from the prior predictive distribution. Points show posterior median estimated titers in $\log_{10}TCID_{50}/mL$ for each sample; lines show 95 % credible intervals. x-axis shows time since sample deposition. Black dotted line shows the single-replicate limit of detection (LOD) of the assay: $10^{0.5}$ $TCID_{50}/mL$ media. Wide coverage of violins relative to datapoints show that priors are agnostic over the titer values of interest, and that the priors regard both fast and slow decay rates as possible.

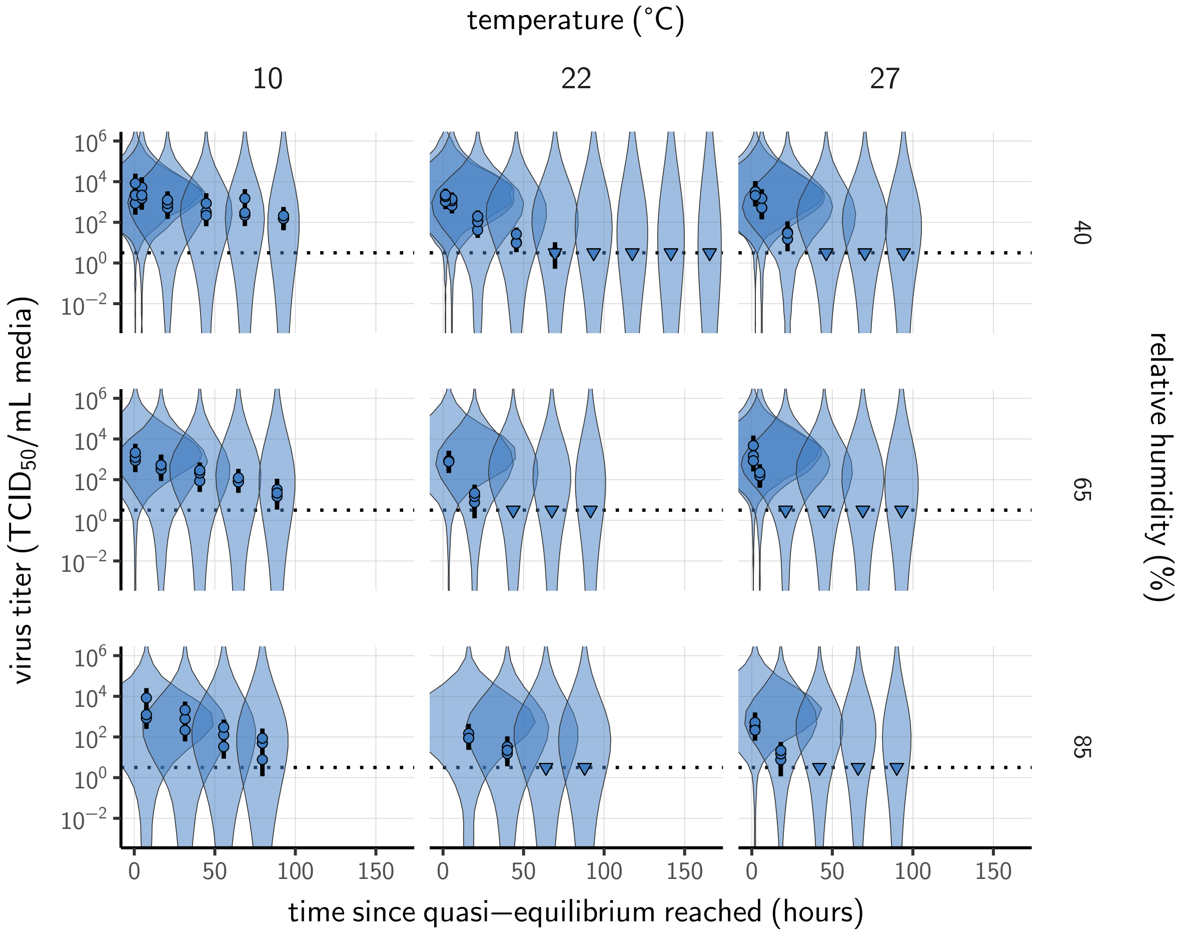

**Figure A14. Prior predictive check for empirical virus decay at quasi-equilibrium for SARS-CoV-2.** Violin plots show distribution of simulated titers sampled from the prior predictive distribution. Points show posterior median estimated titers in $\log_{10}$TCID$_{50}$/mL for each sample; lines show 95 % credible intervals. Time-points with no positive wells for any replicate are plotted as triangles at the approximate single-replicate limit of detection (LOD) of the assay—denoted by a black dotted line at $10^{0.5}$ TCID$_{50}$/mL media—to indicate that a range of sub-LOD values are plausible. Three samples collected at each time-point. x-axis shows time since quasi-equilibrium was reached, as measured in evaporation experiments. Wide coverage of violins relative to datapoints shows that priors are agnostic over the titer values of interest, and that the priors regard both fast and slow decay rates as possible.

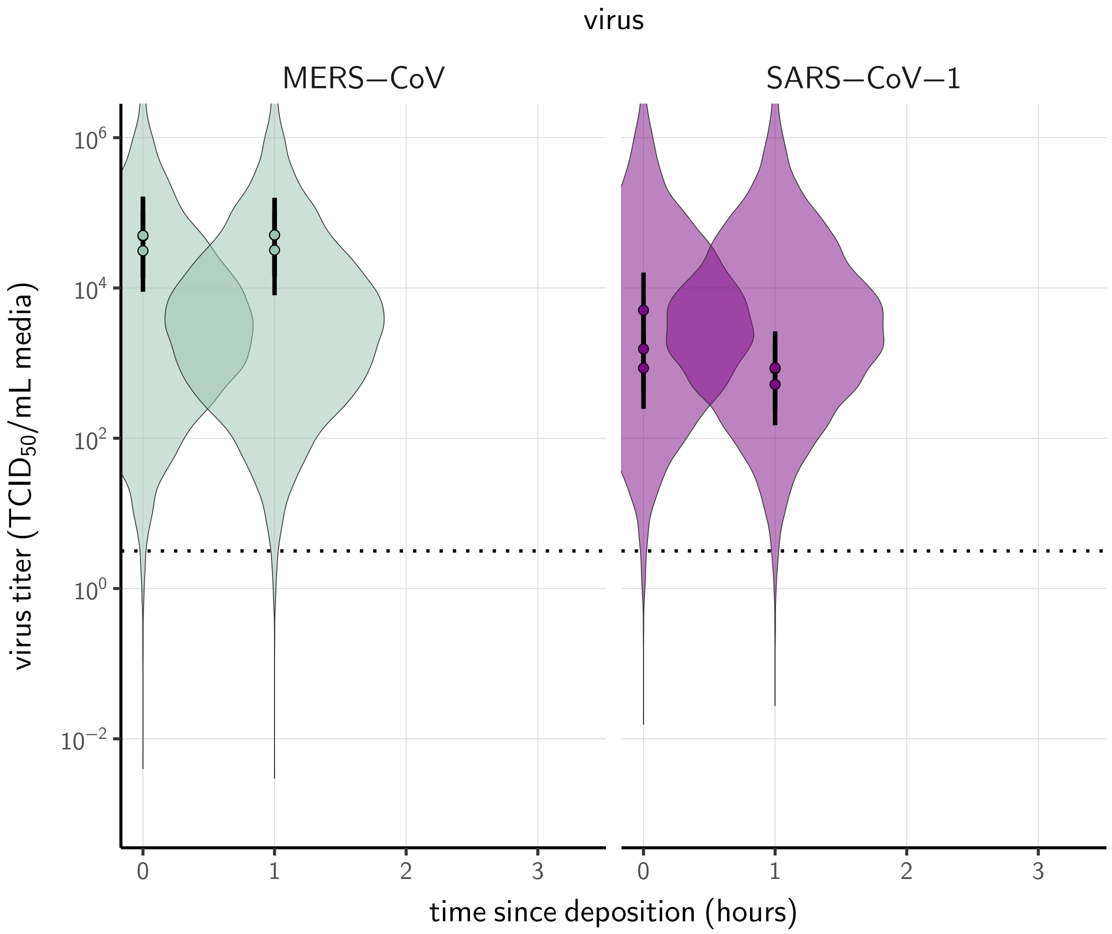

**Figure A15. Prior predictive check for empirical virus decay during the evaporation phase for SARS-CoV-1 and MERS-CoV at 22 °C and 40 % relative humidity.** Violin plots show distribution of simulated titers sampled from the prior predictive distribution. Points show posterior median estimated titers in $\log_{10}$TCID$_{50}$/mL for each sample; lines show 95 % credible intervals. Black dotted line shows the approximate single-replicate limit of detection (LOD) of the assay: $10^{0.5}$ TCID$_{50}$/mL media. Three samples collected at each time-point. x-axis shows time since sample deposition. Lines are truncated at the estimated time quasi-equilibrium was reached. Wide coverage of violins relative to datapoints shows that priors are agnostic over the titer values of interest, and that the priors regard both fast and slow decay rates as possible.

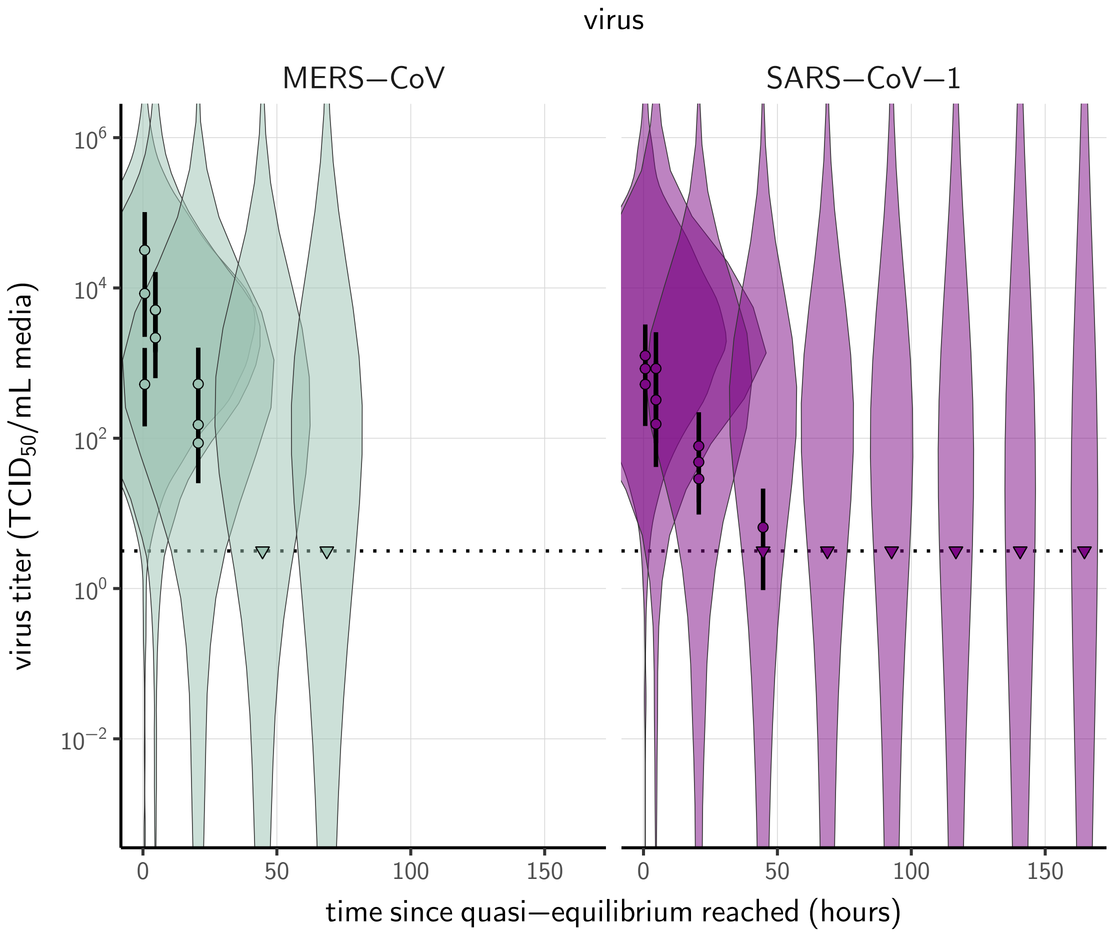

**Figure A16. Prior predictive check for empirical virus decay at quasi-equilibrium for SARS-CoV-1 and MERS-CoV at 22 °C and 40 % relative humidity.** Violin plots show distribution of simulated titers sampled from the prior predictive distribution. Points show posterior median estimated titers in $\log_{10}$TCID$_{50}$/mL for each sample; lines show 95 % credible intervals. Time-points with no positive wells for any replicate are plotted as triangles at the approximate single-replicate limit of detection (LOD) of the assay—denoted by a black dotted line at $10^{0.5}$ TCID$_{50}$/mL media—to indicate that a range of sub-LOD values are plausible. Three samples collected at each time-point. x-axis shows time since quasi-equilibrium was reached, as measured in evaporation experiments. Wide coverage of violins relative to datapoints shows that priors are agnostic over the titer values of interest, and that the priors regard both fast and slow decay rates as possible.

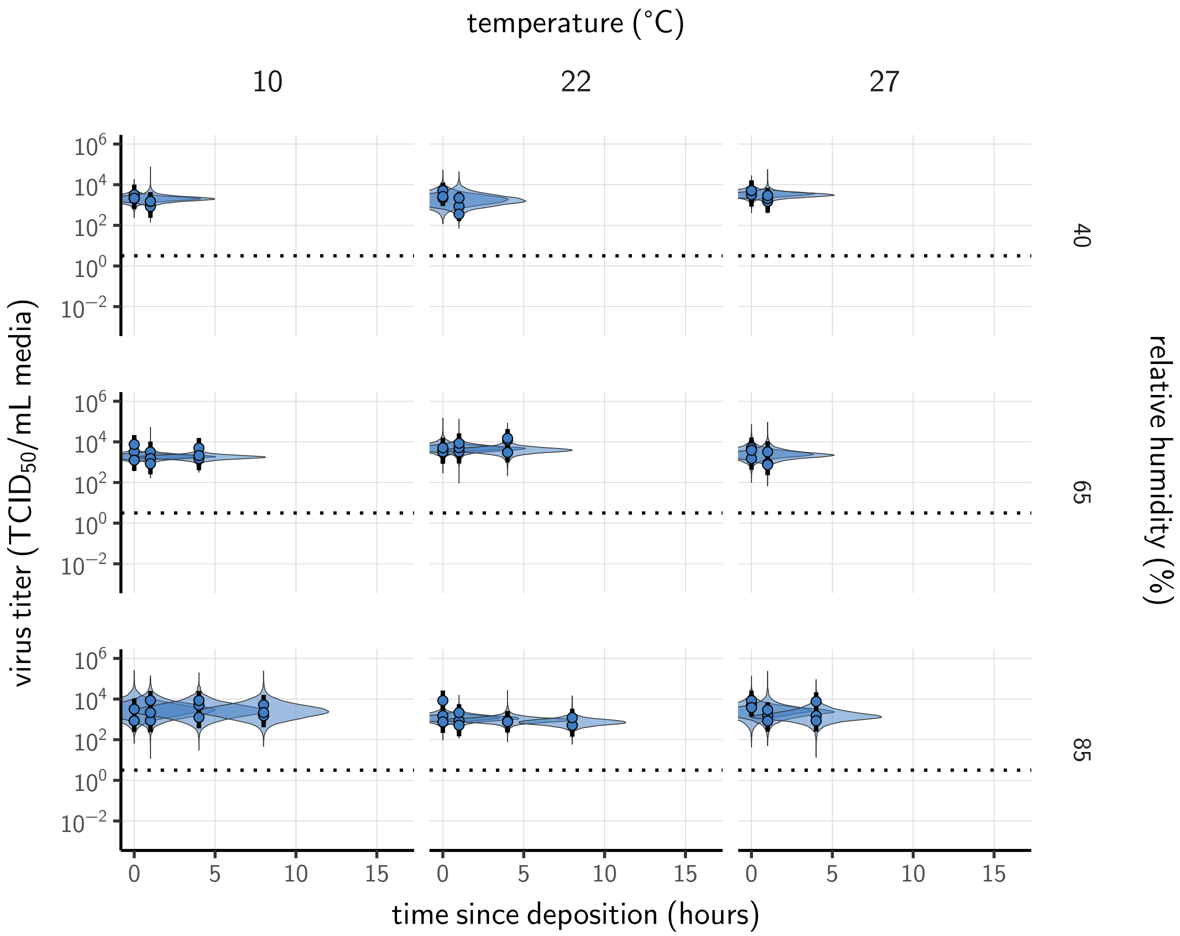

**Figure A17. Posterior predictive check for empirical virus decay during the evaporation phase for SARS-CoV-2.** Violin plots show distribution of simulated titers sampled from the posterior predictive distribution. Points show posterior median estimated titers in $\log_{10}\mathrm{TCID}_{50}/\mathrm{mL}$ for each sample; lines show 95 % credible intervals. Black dotted line shows the approximate single-replicate limit of detection (LOD) of the assay: $10^{0.5}$ $\mathrm{TCID}_{50}/\mathrm{mL}$ media. Three samples collected at each time-point. x-axis shows time since sample deposition. Lines are truncated at the estimated time quasi-equilibrium was reached. Tight correspondence between distribution of posterior simulated titers and independently estimated titers suggests the model fits the data well.

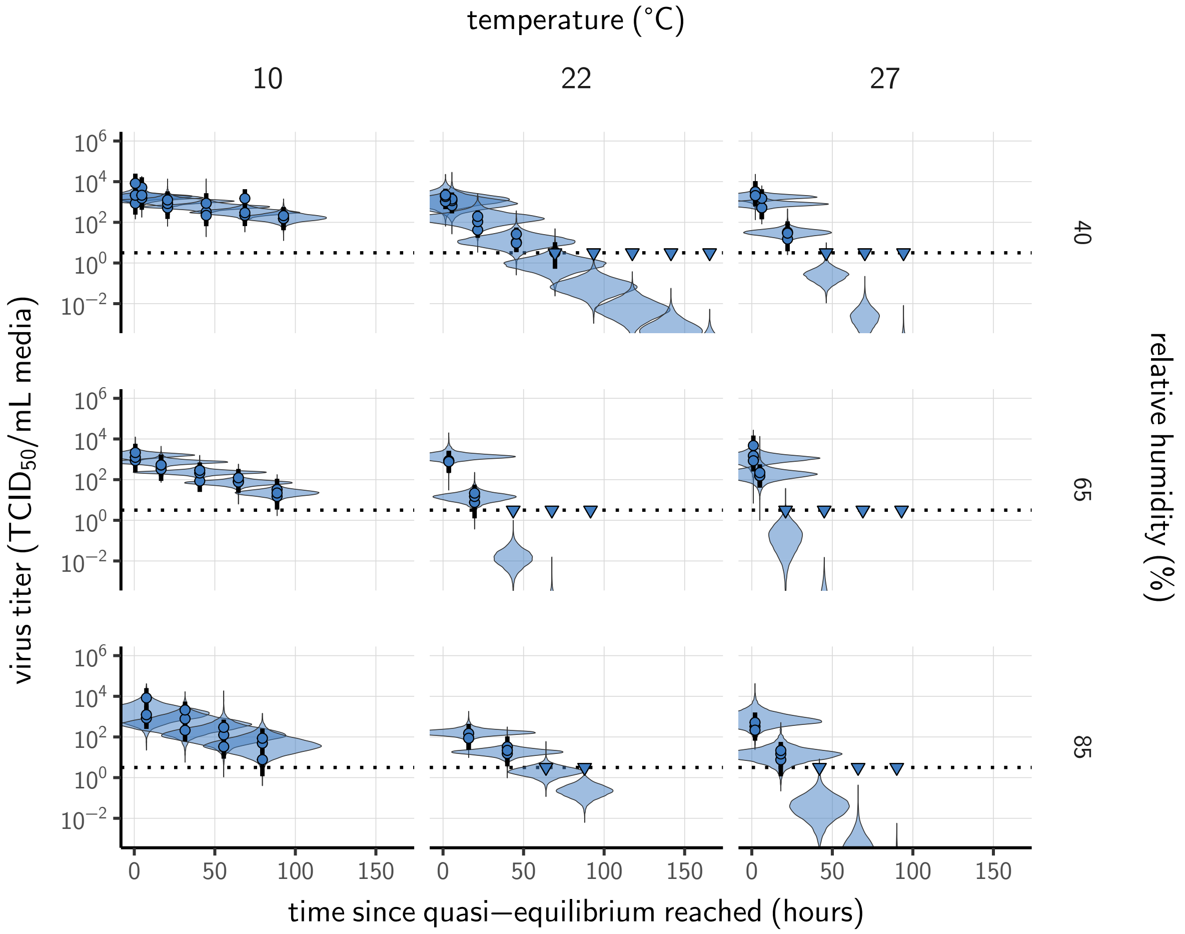

**Figure A18. Posterior predictive check for empirical virus decay at quasi-equilibrium for SARS-CoV-2.** Violin plots show distribution of simulated titers sampled from the posterior predictive distribution. Points show posterior median estimated titers in $\log_{10}$TCID$_{50}$/mL for each sample; lines show 95 % credible intervals. Time-points with no positive wells for any replicate are plotted as triangles at the approximate single-replicate limit of detection (LOD) of the assay—denoted by a black dotted line at $10^{0.5}$ TCID$_{50}$/mL media—to indicate that a range of sub-LOD values are plausible. Three samples collected at each time-point. x-axis shows time since quasi-equilibrium was reached, as measured in evaporation experiments. Tight correspondence between distribution of posterior simulated titers and independently estimated titers suggests the model fits the data well.

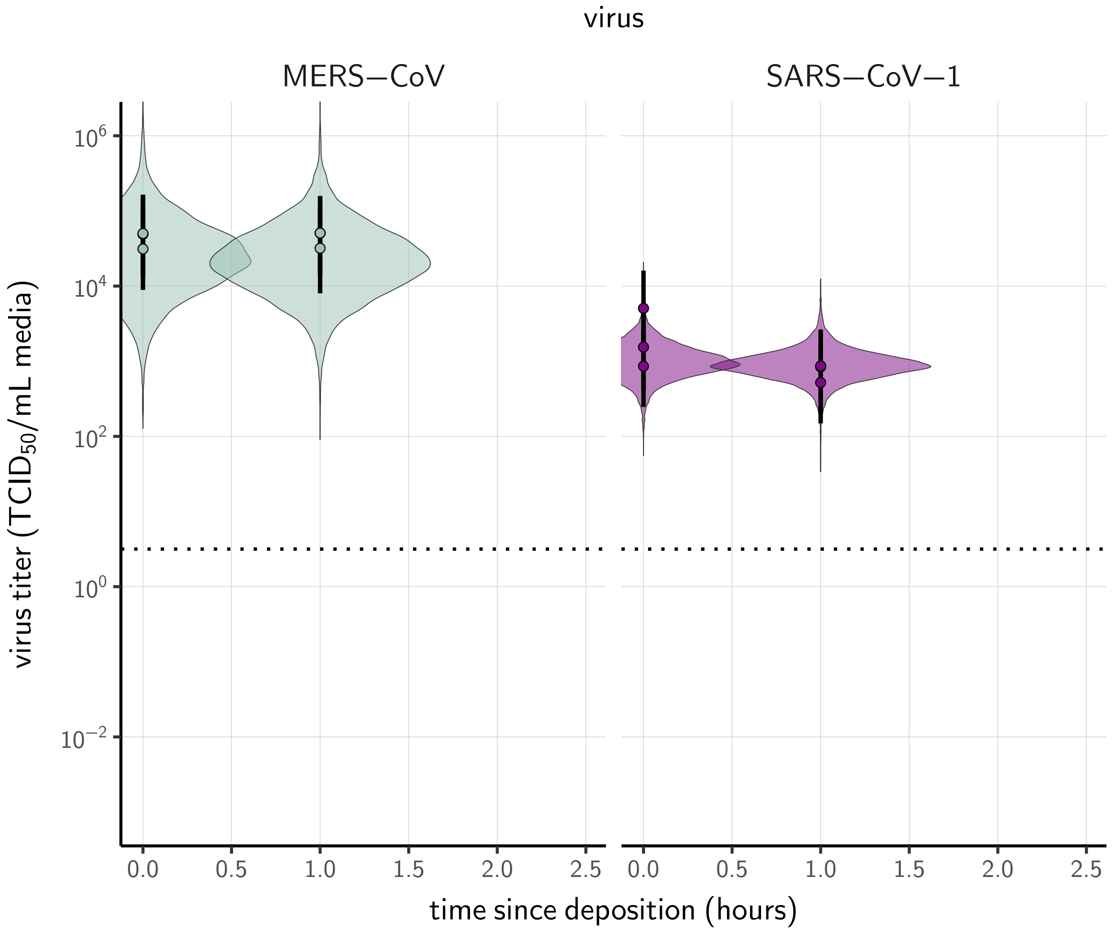

**Figure A19. Posterior predictive check for empirical virus decay during the evaporation phase for SARS-CoV-1 and MERS-CoV at 22 °C and 40 % relative humidity.** Violin plots show distribution of simulated titers sampled from the posterior predictive distribution. Points show posterior median estimated titers in $\log_{10}\text{TCID}_{50}/\text{mL}$ for each sample; lines show 95 % credible intervals. Black dotted line shows the approximate single-replicate limit of detection (LOD) of the assay: $10^{0.5}$ $\text{TCID}_{50}/\text{mL}$ media. Three samples collected at each time-point. x-axis shows time since sample deposition. Lines are truncated at the estimated time quasi-equilibrium was reached. Tight correspondence between distribution of posterior simulated titers and independently estimated titers suggests the model fits the data well.

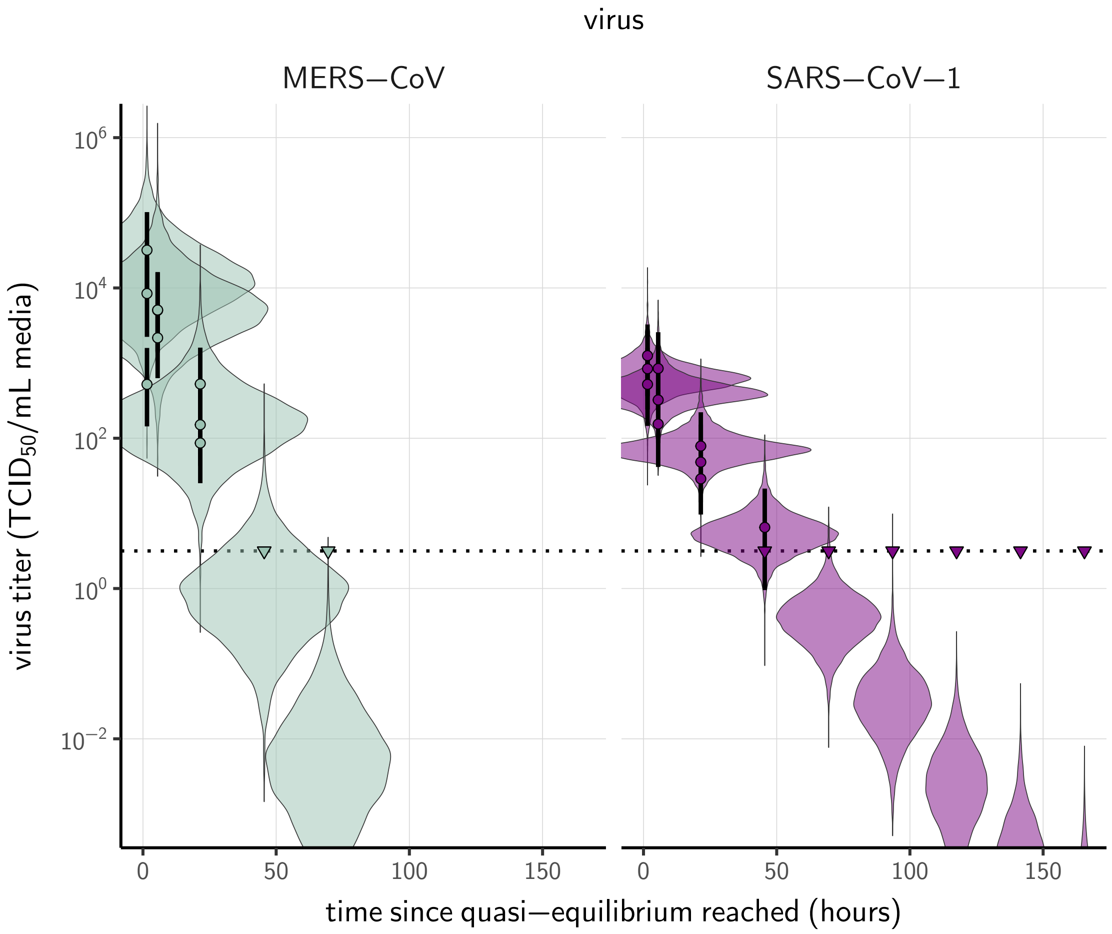

**Figure A20. Posterior predictive check for empirical virus decay at quasi-equilibrium for SARS-CoV-1 and MERS-CoV at 22 °C and 40 % relative humidity.** Violin plots show distribution of simulated titers sampled from the posterior predictive distribution. Points show posterior median estimated titers in $\log_{10}$TCID$_{50}$/mL for each sample; lines show 95 % credible intervals. Time-points with no positive wells for any replicate are plotted as triangles at the approximate single-replicate limit of detection (LOD) of the assay—denoted by a black dotted line at $10^{0.5}$ TCID$_{50}$/mL media—to indicate that a range of sub-LOD values are plausible. Three samples collected at each time-point. x-axis shows time since quasi-equilibrium was reached, as measured in evaporation experiments. Tight correspondence between distribution of posterior simulated titers and independently estimated titers suggests the model fits the data well.

#### 5.5 Mechanistic model of virus decay

##### 5.5.1 Mechanistic model fitting

To fit our mechanistic model (see Main Text, Mechanistic model for temperature and humidity effects), we partitioned experiments according to humidity into two groups: sub-ERH / efflorescence (40 %) and super-ERH / solution (65 %, 85 %). As before, we partitioned each experiment into a evaporation phase and a quasi-equilibrium phase (see section 4.3).

As before, we modeled titers $v_{ij}$ by assuming an initial value $v_{ij0}$ and then modeling decay from that value. We modeled decay during the evaporation phase according to equation 23 and decay during the quasi-equilibrium phase as exponential at a fixed rate $k_i$.

These rates were functions of the temperatures $T_i$ and quasi-equilibrium concentration factors $\frac{[S_{\mathrm{eq}}]}{[S_0]}_i$ according to the mechanistic model.

For all experiments $i$, we modeled decay in solution during the evaporation phase as following equation 23, which follows from the time-varying inactivation rate $k_{\mathrm{ev}}(t)$ given in equation 19:

$$k^i_{\mathrm{ev}}(t) = \frac{w_i(0)}{w_i(t)} A_{\mathrm{sol}} \exp\left(-\frac{E_a}{RT_i}\right) \tag{50}$$

The use of $A_{\mathrm{sol}}$ reflects the assumption that the virus is in solution during the evaporation phase. The $w$ terms model the dynamic concentration factor.

For the quasi-equilibrium phase, we modeled virus decay as exponential at rate $k_{\mathrm{eff}}$ (Main Text equation 2) for efflorescent experiments (at 40 % relative humidity) and as exponential at rate $k_{\mathrm{sol}}$ (Main Text equation 3) for solution experiments (at 65 % or 85 % relative humidity).

That is:

$$k_i = \begin{cases} A_{\mathrm{eff}} \exp\left(-\frac{E_a}{RT_i}\right) & h_i < \mathrm{ERH} \\ \frac{[S_{\mathrm{eq}}]}{[S_0]}_i A_{\mathrm{sol}} \exp\left(-\frac{E_a}{RT_i}\right) & h_i \geq \mathrm{ERH} \end{cases} \tag{51}$$

The resultant titer prediction equation is:

$$v_{ij} = \begin{cases} v_{ij0} + \frac{k_{0i}}{B_i} \log_{10}(1 - B_i\ t_{ij}) & t_{ij} \leq t_i^{\mathrm{eq}} \\ v_{ij0} + \frac{k_{0i}}{B_i} \log_{10}(1 - B_i\ \tau_i) - k_i(t_{ij} - t_i^{\mathrm{eq}}) & t_{ij} > t_i^{\mathrm{eq}} \end{cases} \tag{52}$$

where $k_{0i} = k_{\mathrm{ev}}^i(0)$ and $t_i^{\mathrm{eq}}$ is the modeled time to quasi-equilibrium ($t_i^{\mathrm{eq}} = \tau_i$ for the measured concentration fit and $t_i^{\mathrm{eq}} = \bar{\tau}_i$ for the modeled concentration fit; see section 4.3).

As in the simple regression model, we then used the direct-from-well data likelihood function described above under the assumption that our observed well data $y_{idk}$ reflected the titers $v_{ij}$ predicted by the mechanistic model per equation 52.

We estimated the joint posterior for all parameters. That is, activation energies $E_a$ and asymptotic reaction rates $A$ are estimated in light of evaporative mass loss rates $\beta_i$ and resulting times to quasi-equilibrium $t_i^{\mathrm{eq}}$, and vice versa, for maximally informative propagation of uncertainty.

##### 5.5.2 Concentration factor

In our evaporation experiments, we measured $m_i(0)$ and $m_i(\infty)$, the initial and final total masses, respectively, of the deposited droplet under the temperature and humidity conditions of experiment $i$.

For experiment $i$, we denote the initial mass of water by $w_i(0)$, the final mass of water by $w_i(\infty)$ and the mass of solutes, which which we assume is conserved, by $s_i$. Then:

$$\begin{aligned} m_i(0) &= w_i(0) + s_i \\ m_i(\infty) &= w_i(\infty) + s_i \end{aligned} \tag{53}$$

Denote the initial and final mass fractions of solutes in experiment $i$ by $Y_i(0) = \frac{s_i}{w_i(0)+s_i}$ and $Y_i(\infty) = \frac{s_i}{w_i(\infty)+s_i}$, respectively.

We treated the $Y_i(0)$ as an estimated parameter, assuming that it had the same value across all experiments: $Y_i(0) = Y(0)$.

To estimate the parameters $\alpha_c$ and $\alpha_s$ for $\frac{[S_{\mathrm{eq}}]}{[S_0]}$ as a function of $h$, we needed to predict the observed final total mass, $m(\infty)$ as a function of $h$.

By definition:

$$m_i(\infty) = \frac{s_i}{Y_i(\infty)} \tag{54}$$

We can find $Y_i(\infty)$ by using the fact that $\frac{Y(\infty)}{1-Y(\infty)} = \frac{X(\infty)}{1-X(\infty)} = r(\infty)$, where $X(\infty)$ is the quasi-equilibrium molar fraction of solutes. So $Y_i(\infty) = \frac{r_i(\infty)}{r_i(\infty)+1}$. Since $s_i = Y_i(0)m_i(0)$, it follows that the predicted quasi-equilibrium total mass for experiment $i$, $\bar{m}_i(\infty)$, is:

$$\bar{m}_i(\infty) = \frac{r_i(\infty)+1}{r_i(\infty)} Y_i(0) m_i(0) \tag{55}$$

We modeled $r_i(\infty)$ according to equation 11. Using equation 55, we estimated $Y(0)$ and the parameters $\alpha_c$ and $\alpha_s$ of equation 11 from our data. We modeled the observed log final total masses $\ln(m_i(\infty))$ as normally distributed about the log predicted quasi-equilibrium total masses $\ln(\bar{m}_i(\infty))$ with an estimated standard deviation $\sigma_m$:

$$\ln(m_i(\infty)) \sim \text{Normal}(\ln(\bar{m}_i(\infty)), \sigma_m) \tag{56}$$

We assumed that quasi-equilibrium total mass values measured below the ERH were equivalent to the quasi-equilibrium total mass values at the ERH; this allowed us to use the 40 % RH (sub-ERH) evaporation data points to add additional resolution to the estimation of $\alpha_c$ and $\alpha_s$.

##### 5.5.3 Mechanistic model versions

As described in the Main Text, we fit the mechanistic model in two ways: a measured concentration fit and a modeled concentration fit. The measured concentration fit is the most direct snapshot of inactivation in our data, but it can only predict inactivation rates at RH levels where $\frac{[S_{\text{eq}}]}{[S_0]}$ is known. The modeled concentration fit estimates the mechanistic parameters alongside the relationship between RH and $\frac{[S_{\text{eq}}]}{[S_0]}$ (see section 4.1), which allows for principled extrapolation to unobserved RH values.

**Measured concentration fit.** In the measured concentration fit, we calculated the concentration factor for the $i^{\text{th}}$ experiment, $\frac{[S_{\text{eq}}]}{[S_0]}_i$ according to equation 9 using the measured initial and final total masses $m_i(0)$ and $m_i(\infty)$ and the estimated parameter $Y(0)$:

$$\frac{[S_{\mathrm{eq}}]}{[S_0]}_i = \frac{w_i(0)}{w_i(\infty)} = \frac{m_i(0) - s_i}{m_i(\infty) - s_i} = \frac{m_i(\infty) - Y(0)m_i(0)}{m_i(\infty) - Y(0)m_i(0)} \tag{57}$$

**Modeled concentration fit.** In the modeled concentration fit, we calculated $\frac{[S_{\mathrm{eq}}]}{[S_0]}_i$ from the ambient relative humidity according to equation 12, substituting $\frac{1-Y(0)}{Y(0)}$ for $\frac{1}{r(0)} = \frac{1-X(0)}{X(0)}$, since the two ratios are equal:

$$\frac{[S_{\mathrm{eq}}]}{[S_0]}_i = \frac{1}{r(0)} \frac{X_i(\infty)}{1 - X_i(\infty)} = \left(\frac{1 - Y(0)}{Y(0)}\right)\left(\frac{-\ln(h)}{\alpha_s}\right)^{\frac{1}{\alpha_c}} \tag{58}$$

Note that this means that $\alpha_s$ and $\alpha_c$ for the modeled concentration fit were estimated not only in light of the measured droplet masses but also in light of the measured virus titers, filtered through the mechanistic model of inactivation.

##### 5.5.4 Mechanistic model prior distributions

**Activation energies and asymptotic reaction rates.** To place priors on $E_a$ and $A$ in an interpretable manner, we placed them not on the parameter pairs themselves but rather on the solution and efflorescent half-lives at 20 °C, $\eta_{\mathrm{sol}}(20)$ and $\eta_{\mathrm{eff}}(20)$, and the ratios of virus decay rate at 30 °C to the virus decay rate at 20 °C, $k_{\mathrm{sol}}(30)/k_{\mathrm{sol}}(20)$ and $k_{\mathrm{eff}}(30)/k_{\mathrm{eff}}(20)$.

These quantities fully determine the solution and efflorescence $E_a$ and $A$ values.

Decay rate ratios are related to activation energies by:

$$E_a = \frac{R \ln\left(\frac{k(T_1)}{k(T_2)}\right)}{\frac{1}{T_2} - \frac{1}{T_1}} \tag{59}$$

where the temperatures are given in Kelvin.

The 20 °C half-lives $\eta(20)$ in hours imply associated exponential decay rates in $\log_{10}$ $\mathrm{TCID}_{50}$/mL/h: $k(20) = \frac{\log_{10}(2)}{\eta(20)}$. Given an activation energy $E_a$ and a known decay rate $k(T)$ for a given temperature $T$ in Kelvin, one can calculate the asymptotic rate $A$:

$$\ln(A) = \ln\left(k(T)\right) + \frac{E_a}{RT} \tag{60}$$

Note that for $A_{\mathrm{sol}}$, this is the asymptotic rate at the initial concentration (i.e. when $\frac{[S(t)]}{[S_0]} = \frac{[S_0]}{[S_0]} = 1$).

We placed a Normal prior on the log of the half-life at 20 °C. Since $\eta_{\mathrm{eff}}(20)$ and $\eta_{\mathrm{sol}}(20)$ are the effloresced quasi-equilibrium and unconcentrated solution half-lives, respectively, we used the same prior as that used for the evaporation phase half-life (see section 5.4.2):

$$\begin{aligned} \ln(\eta_{\mathrm{eff}}(20)) &\sim \mathrm{Normal}(\ln(24), 1.25) \\ \ln(\eta_{\mathrm{sol}}(20)) &\sim \mathrm{Normal}(\ln(24), 1.25) \end{aligned} \tag{61}$$

We placed a Half-Normal prior on the natural log of the decay rate ratios:

$$\ln\left(\frac{k(30)}{k(20)}\right) \sim \text{Half-Normal}(0, 1) \tag{62}$$

Note that this means virus inactivation must become more rapid with temperature, another way in which our model's fitted parameters are not truly free, and thus good fits should not necessarily be expected unless the model describes reality.

For fits with distinct $E_a^{\mathrm{sol}}$ and $E_a^{\mathrm{eff}}$, we used the same Half-Normal$(0, 1)$ prior for both $\ln\left(\frac{k_{\mathrm{eff}}(30)}{k_{\mathrm{eff}}(20)}\right)$ and $\ln\left(\frac{k_{\mathrm{sol}}(30)}{k_{\mathrm{sol}}(20)}\right)$.

**Titer intercepts.** We handled the titer intercepts $v_{ij0}$ for the mechanistic model identically to how they were handled in the simple regression model, with identical priors (see equations 46, 47, and 48 in sections 5.4.1 and 5.4.2). We reproduce those equations here for reference:

$$\begin{aligned} v_{ij0} &\sim \mathrm{Normal}(\bar{v}_{i0}, \sigma_i) \\ \bar{v}_{i0} &\sim \mathrm{Normal}(2.5, 1) \\ \sigma_i &\sim \text{Half-Normal}(0, 0.5) \end{aligned}$$

**Concentration factor.** We placed a Normal prior on the log of the initial solute mass fraction $Y_0$, with a mode given by the approximate solute mass fraction for Dulbecco's Modified Eagle Medium (DMEM) reported by the manufacturer (Sigma Aldrich, reference D6546 [72]).

$$\ln(Y_0) \sim \text{Normal}(\ln(0.011), 0.33) \tag{63}$$

We placed Normal priors on the parameters $c_c$ and $c_s$ that model quasi-equilibrium mole fraction ratio as a function of humidity in equation 12:

$$c_c \sim \text{Normal}(0, 0.33)$$
$$c_s \sim \text{Normal}(0, 0.33)$$

Note that this results in lognormal priors on $\alpha_c$ and $\alpha_s$.

We placed a Normal prior on the standard deviation $\sigma_m$ of the observed log quasi-equilibrium mass about its predicted value.

$$\sigma_m \sim \text{Normal}(0, 1) \tag{64}$$

###### 5.5.5 Mechanistic model predictive checks

We assessed the appropriateness of prior distribution choices using prior predictive checks and assessed goodness of fit for the estimated model using posterior predictive checks. Prior checks suggested that prior distributions were agnostic over the parameter values of interest, and posterior checks suggested a good fit of the model to the data. The resultant checks for the measured concentration and modeled concentration versions of the mechanistic model of virus decay are shown below (Fig. A21– A28).

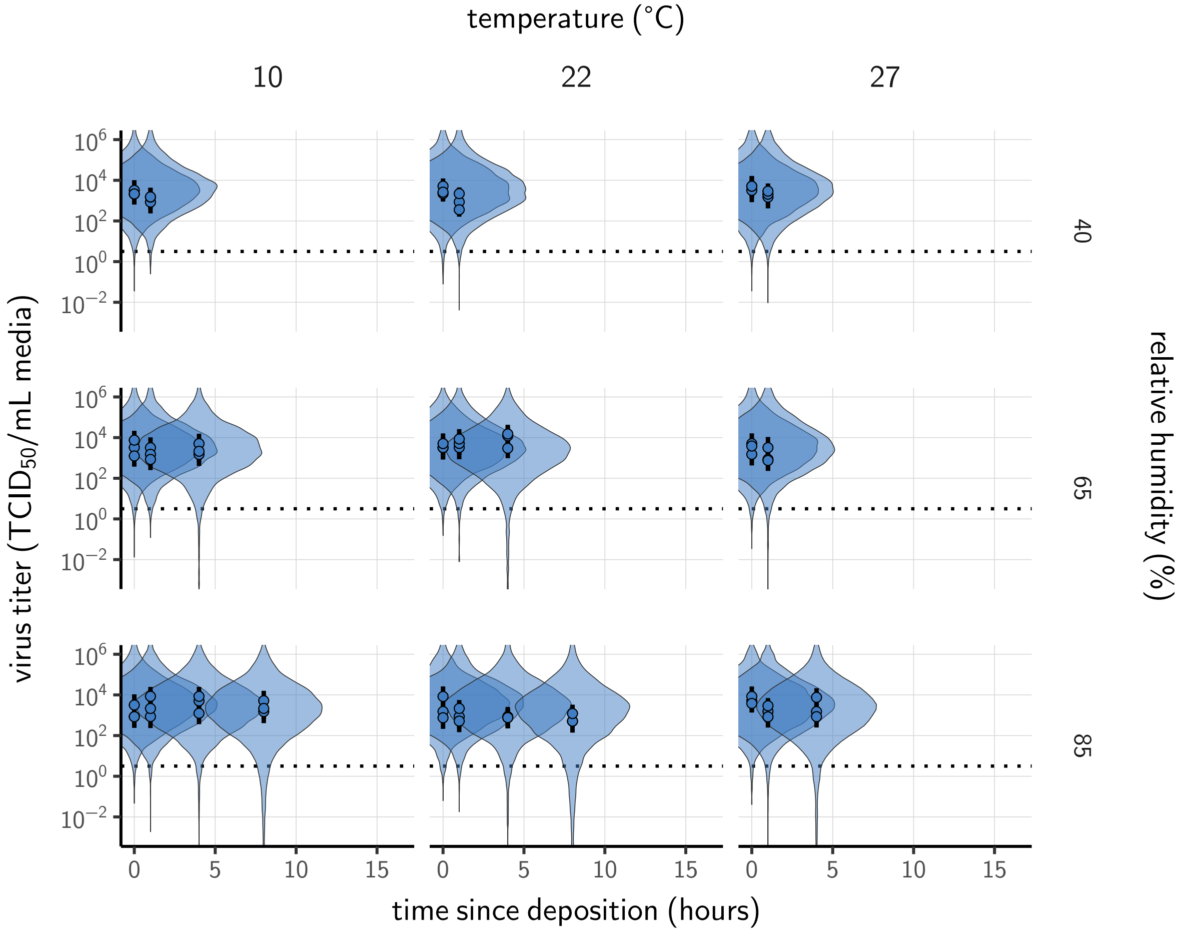

**Figure A21. Prior predictive check for measured concentration fit during the evaporation phase.** Violin plots show distribution of simulated titers sampled from the prior predictive distribution. Points show posterior median estimated titers in $\log_{10}$TCID$_{50}$/mL for each sample; lines show 95 % credible intervals. Black dotted line shows the approximate single-replicate limit of detection (LOD) of the assay: $10^{0.5}$ TCID$_{50}$/mL media. Three samples collected at each time-point. x-axis shows time since sample deposition. Lines are truncated at the estimated time quasi-equilibrium was reached. Wide coverage of violins relative to datapoints shows that priors are agnostic over the titer values of interest, and that the priors regard both fast and slow decay rates as possible.

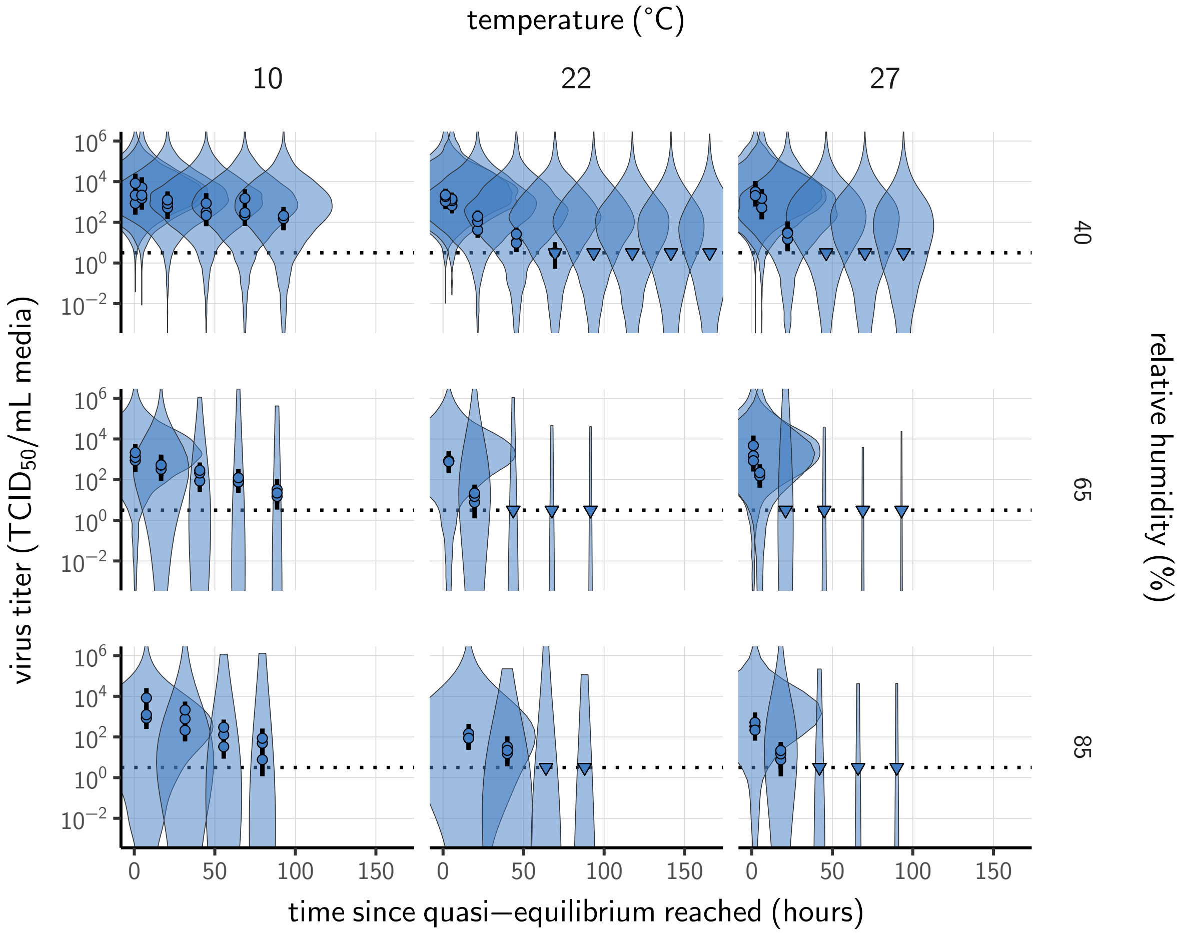

**Figure A22. Prior predictive check for measured concentration fit at quasi-equilibrium.** Violin plots show distribution of simulated titers sampled from the prior predictive distribution. Points show posterior median estimated titers in $\log_{10}$TCID$_{50}$/mL for each sample; lines show 95 % credible intervals. Time-points with no positive wells for any replicate are plotted as triangles at the approximate single-replicate limit of detection (LOD) of the assay—denoted by a black dotted line at $10^{0.5}$ TCID$_{50}$/mL media—to indicate that a range of sub-LOD values are plausible. Three samples collected at each time-point. x-axis shows time since quasi-equilibrium was reached, as measured in evaporation experiments. Wide coverage of violins relative to datapoints shows that priors are agnostic over the titer values of interest, and that the priors regard both fast and slow decay rates as possible.

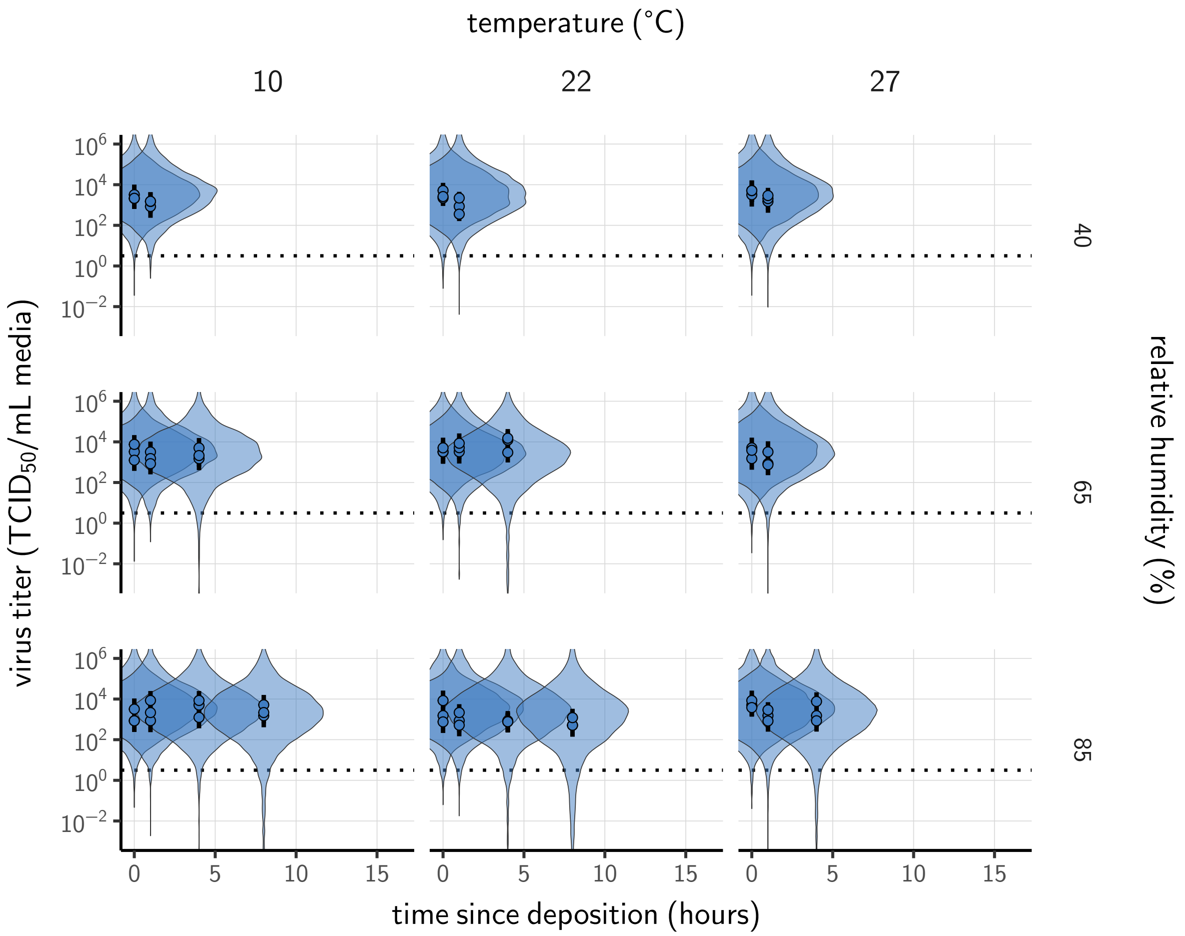

**Figure A23. Prior predictive check for modeled concentration fit during the evaporation phase.** Violin plots show distribution of simulated titers sampled from the prior predictive distribution. Points show posterior median estimated titers in $\log_{10}$TCID$_{50}$/mL for each sample; lines show 95 % credible intervals. Black dotted line shows the approximate single-replicate limit of detection (LOD) of the assay: $10^{0.5}$ TCID$_{50}$/mL media. Three samples collected at each time-point. x-axis shows time since sample deposition. Lines are truncated at the estimated time quasi-equilibrium was reached. Wide coverage of violins relative to datapoints shows that priors are agnostic over the titer values of interest, and that the priors regard both fast and slow decay rates as possible.

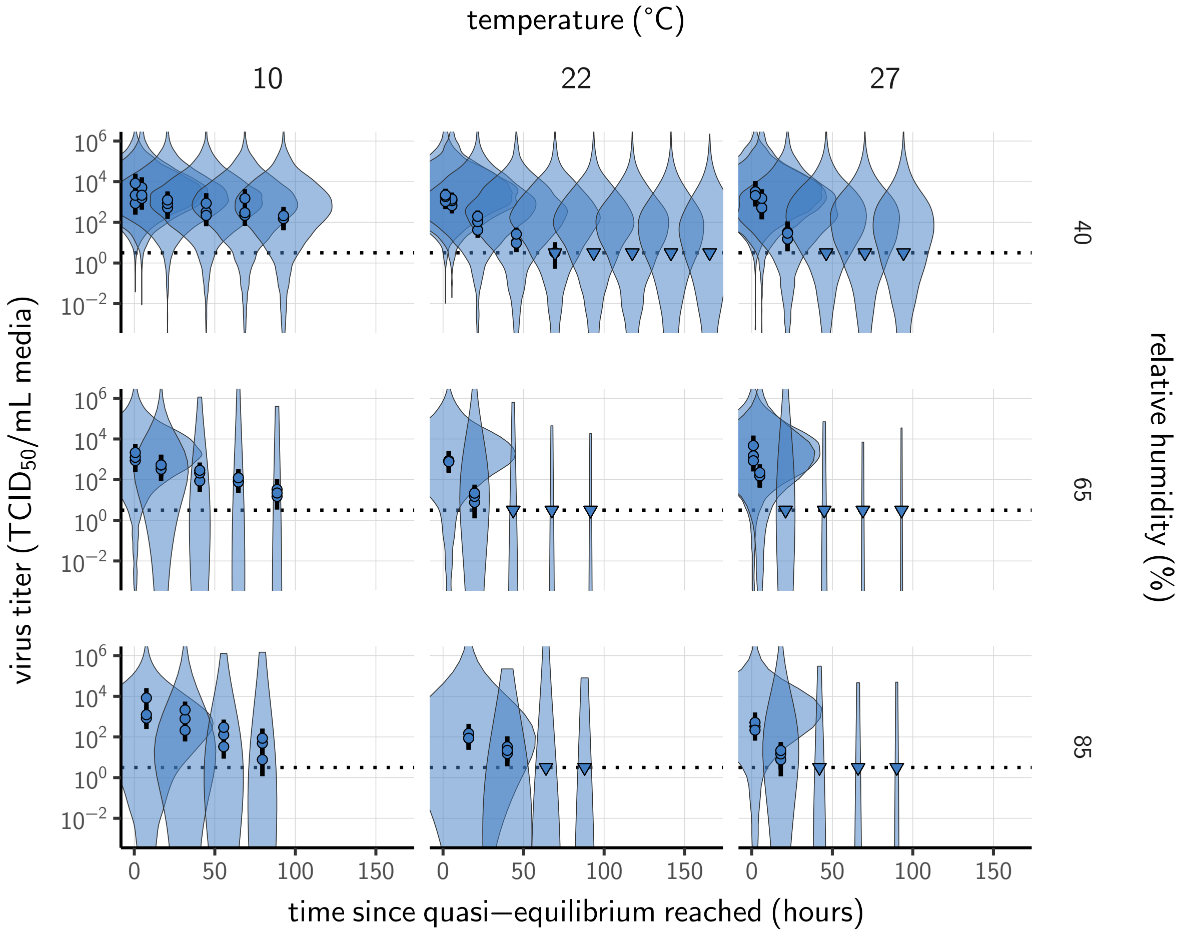

**Figure A24. Prior predictive check for modeled concentration fit at quasi-equilibrium.** Violin plots show distribution of simulated titers sampled from the prior predictive distribution. Points show posterior median estimated titers in $\log_{10}$TCID$_{50}$/mL for each sample; lines show 95 % credible intervals. Time-points with no positive wells for any replicate are plotted as triangles at the approximate single-replicate limit of detection (LOD) of the assay—denoted by a black dotted line at $10^{0.5}$ TCID$_{50}$/mL media—to indicate that a range of sub-LOD values are plausible. Three samples collected at each time-point. x-axis shows time since quasi-equilibrium was reached, as measured in evaporation experiments. Wide coverage of violins relative to datapoints shows that priors are agnostic over the titer values of interest, and that the priors regard both fast and slow decay rates as possible.

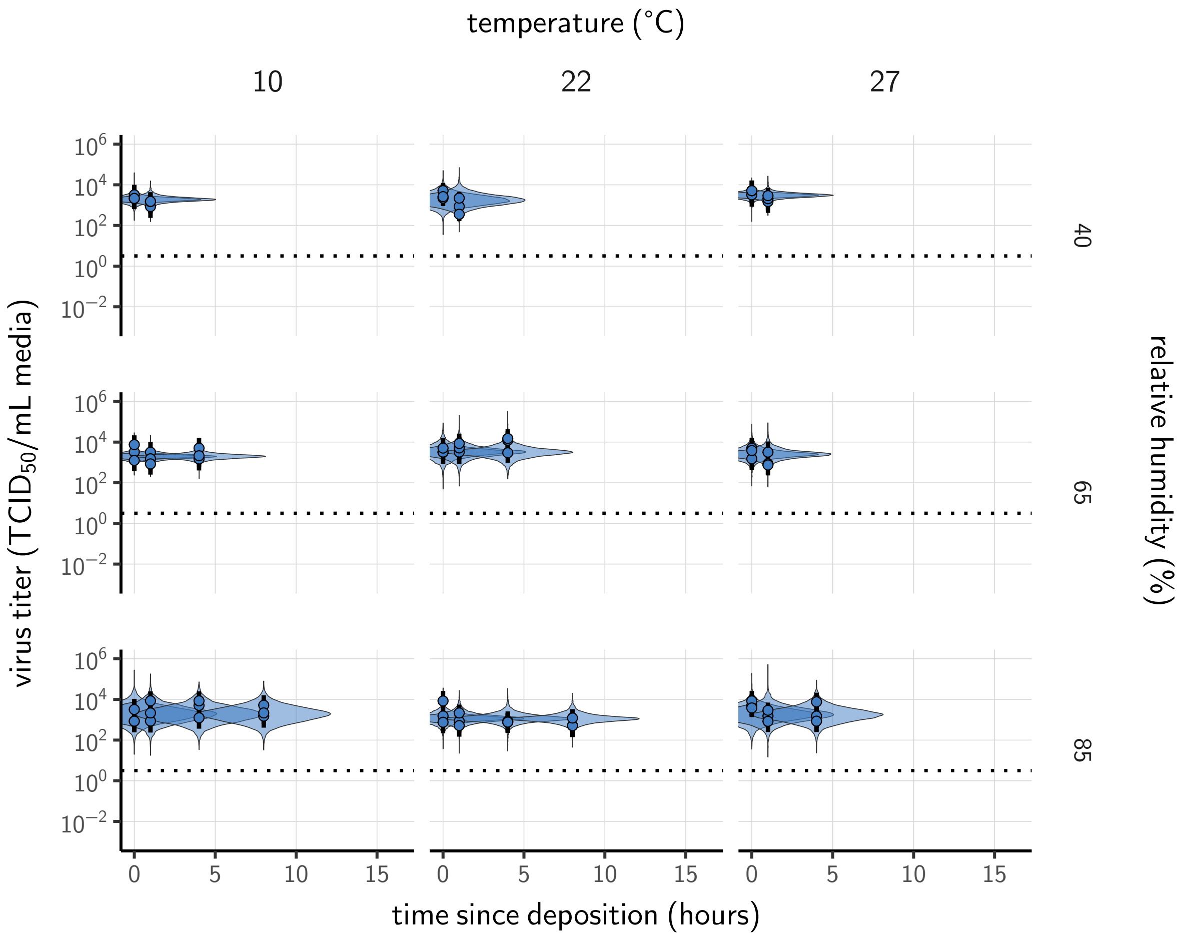

**Figure A25. Posterior predictive check for measured concentration fit during the evaporation phase.** Violin plots show distribution of simulated titers sampled from the posterior predictive distribution. Points show posterior median estimated titers in $\log_{10}$TCID$_{50}$/mL for each sample; lines show 95 % credible intervals. Black dotted line shows the approximate single-replicate limit of detection (LOD) of the assay: $10^{0.5}$ TCID$_{50}$/mL media. Three samples collected at each time-point. x-axis shows time since sample deposition. Lines are truncated at the estimated time quasi-equilibrium was reached. Tight correspondence between distribution of posterior simulated titers and independently estimated titers suggests the model fits the data well.

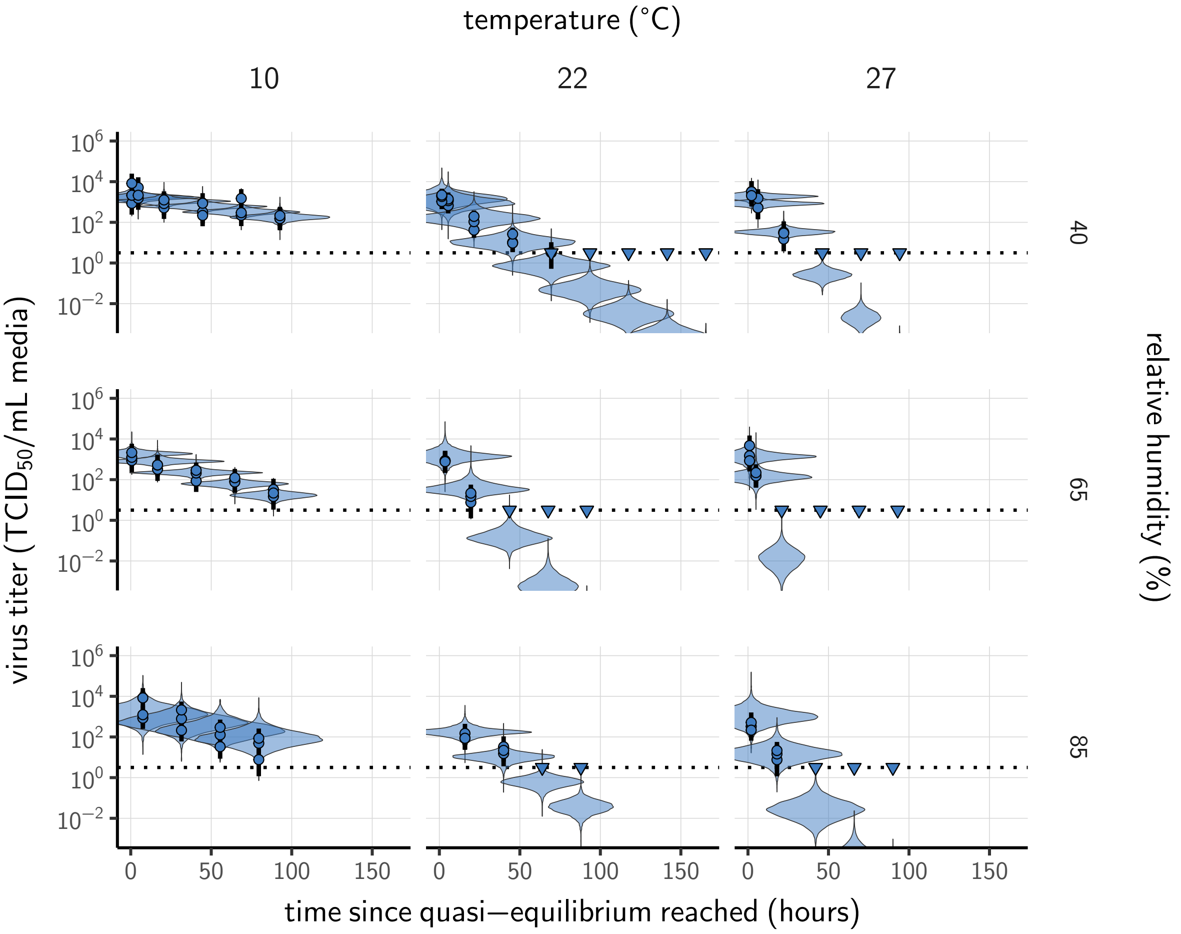

**Figure A26. Posterior predictive check for measured concentration fit at quasi-equilibrium.** Violin plots show distribution of simulated titers sampled from the posterior predictive distribution. Points show posterior median estimated titers in $\log_{10}$TCID$_{50}$/mL for each sample; lines show 95 % credible intervals. Time-points with no positive wells for any replicate are plotted as triangles at the approximate single-replicate limit of detection (LOD) of the assay—denoted by a black dotted line at $10^{0.5}$ TCID$_{50}$/mL media—to indicate that a range of sub-LOD values are plausible. Three samples collected at each time-point. x-axis shows time since quasi-equilibrium was reached, as measured in evaporation experiments. Tight correspondence between distribution of posterior simulated titers and independently estimated titers suggests the model fits the data well.

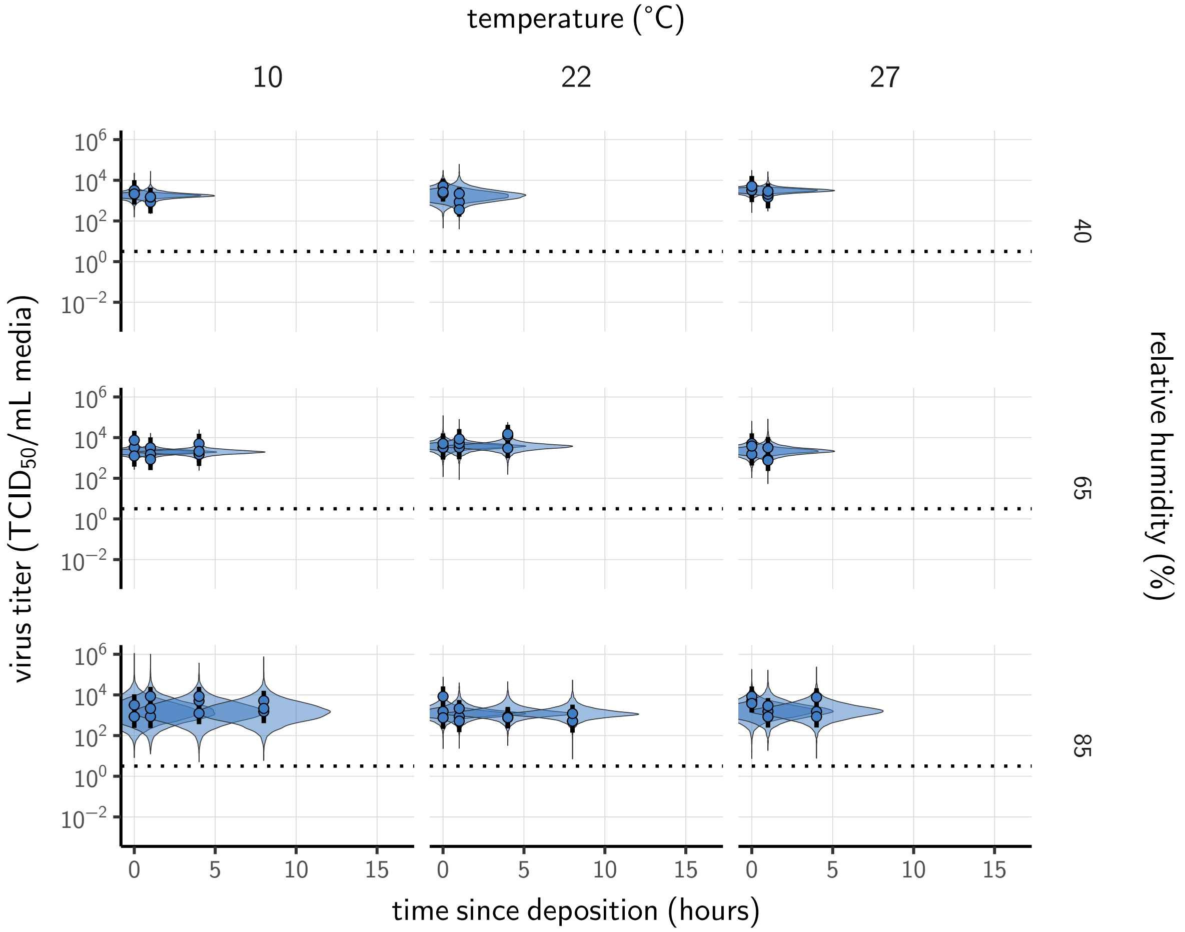

**Figure A27. Posterior predictive check for modeled concentration fit during the evaporation phase.** Violin plots show distribution of simulated titers sampled from the posterior predictive distribution. Points show posterior median estimated titers in $\log_{10}$TCID$_{50}$/mL for each sample; lines show 95 % credible intervals. Black dotted line shows the approximate single-replicate limit of detection (LOD) of the assay: $10^{0.5}$ TCID$_{50}$/mL media. Three samples collected at each time-point. x-axis shows time since sample deposition. Lines are truncated at the estimated time quasi-equilibrium was reached. Tight correspondence between distribution of posterior simulated titers and independently estimated titers suggests the model fits the data well.

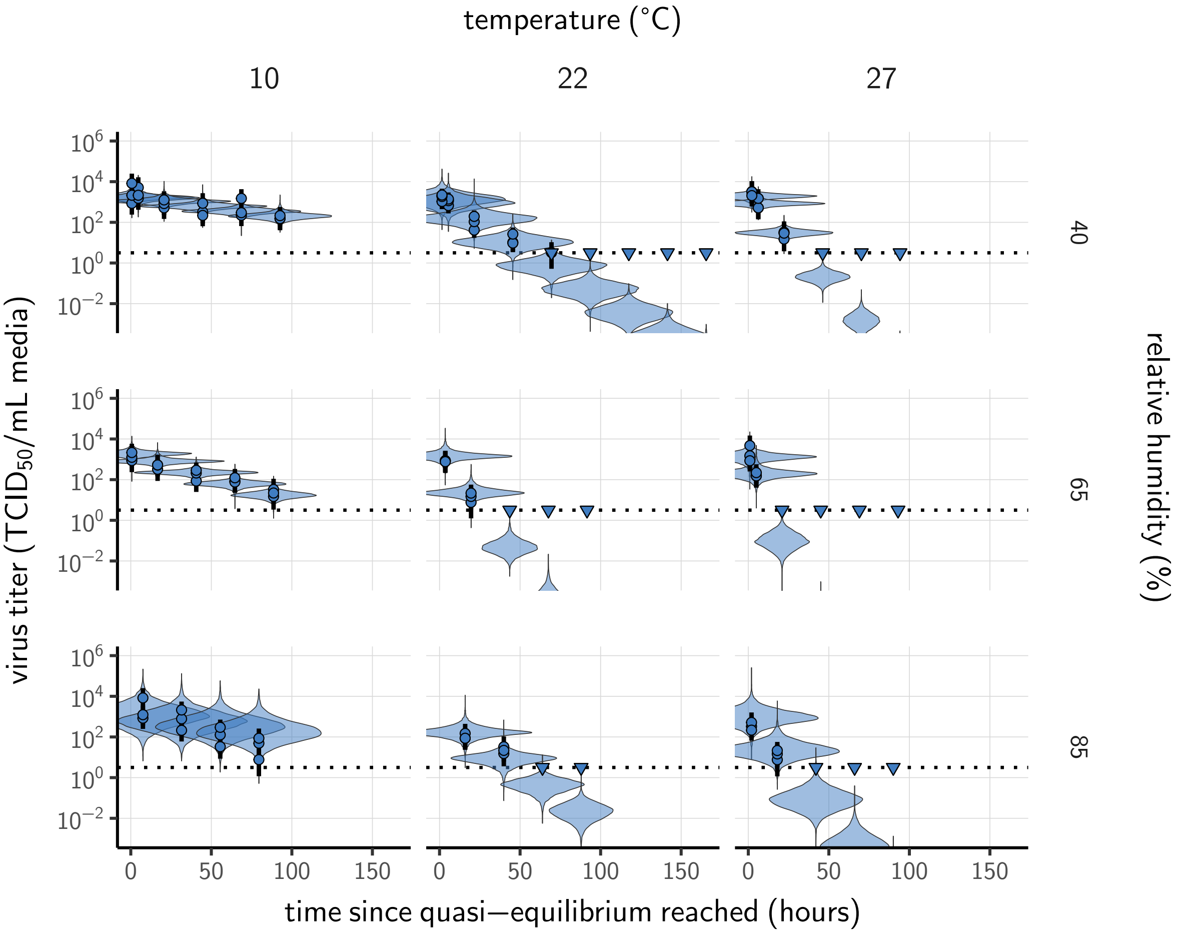

**Figure A28. Posterior predictive check for modeled concentration fit at quasi-equilibrium.** Violin plots show distribution of simulated titers sampled from the posterior predictive distribution. Points show posterior median estimated titers in $\log_{10}$TCID$_{50}$/mL for each sample; lines show 95 % credible intervals. Time-points with no positive wells for any replicate are plotted as triangles at the approximate single-replicate limit of detection (LOD) of the assay—denoted by a black dotted line at $10^{0.5}$ TCID$_{50}$/mL media—to indicate that a range of sub-LOD values are plausible. Three samples collected at each time-point. x-axis shows time since quasi-equilibrium was reached, as measured in evaporation experiments. Tight correspondence between distribution of posterior simulated titers and independently estimated titers suggests the model fits the data well.

### 6 Meta-analysis of human coronavirus half-lives

#### 6.1 Study selection and data extraction

We screened the Web of Science Core Collection database on May 31, 2020, using the following key words: "coronavir* AND (stability OR viability OR inactiv*) AND (temperature OR heat OR humidity)" (83 records). We also considered opportunistically identified pre-prints (up to July 6, 2020) and studies referenced in full-texts assessed for eligibility and potentially reporting datasets of interest (22 records). We then selected publications reporting data of viral stability for human coronaviruses (MERS, SARS-CoV-1, SARS-CoV-2, HCoV-OC43, HCoV-HKU1, HCoV-229E and HCoV-NL63) and for at least two temperature or humidity conditions. Considering the impact of medium composition and contact surface on virus inactivation kinetics [64, 15], we also filtered the selected studies based on these criteria. The complete selection procedure is described in Fig. A29 following the Preferred Reporting Items for Systematic Reviews and Meta-Analyses (PRISMA) [79]. Studies included in our analysis are listed in Table A2.

We compiled data in the form of viral titer or relative infectivity across time, depending on how they were reported in the selected studies. Data were most often reported as mean ± variation (standard deviation or 95% confidence interval) across replicates per time-point and experimental condition. However, as number of replicates and measured variation was not systematically reported, we did not include this information in our analyses. We extracted data from tables and from figures manually using the WebPlotDigitizer application [84]. We also recorded metadata including environmental conditions (temperature and relative humidity), contact surface, and medium composition and volume. The complete dataset is available in the online data and code repository.

Among the selected studies, we sub-selected data to be included in our meta-analysis based on the same criteria. In particular, we restricted the dataset to suspensions composed of respiratory secretions, or cell culture or virus transportation media supplemented only with antibiotics and up to 10% fetal calf serum and 1% glutamine; we also restricted the dataset to stability measurements conducted in bulk medium suspensions, or using droplets deposited on inert surfaces (including steel and polypropylene) or on skin. The final dataset consisted of 38 experimental conditions, covering 17 temperature-humidity combinations and five human coronaviruses (HCoV-229E, HCoV-OC43, MERS-CoV, SARS-CoV-1 and SARS-CoV-2) listed in Table A2.

#### 6.2 Estimation of virus decay in the literature

##### 6.2.1 Estimation model and priors

We converted all data from the literature into $\log_{10}$ fraction of viable virus remaining (Fig. A30–A31). That is, we normalized the reported quantity of viable virus to the earliest measurement—if the authors had not already done so—and expressed time as time elapsed since earliest measurement. We then estimated half-lives independently for each environmental condition $j$ in each study $i$ by fitting a Bayesian exponential decay model with exponential decay rates $\lambda_{ij}$ for each experiment $j$. We treated each reported measurement $y_{ijk}$ (in $\log_{10}$ fraction viable) from experiment $j$ of study $i$ as normally distributed about the predicted $\log_{10}$ fraction viable $\bar{f}_{ijk}$, with an unknown standard deviation $\sigma_{\text{mat}}(i, j)$ estimated independently for each material in study $i$, but shared across all temperature/humidity conditions for that study-material pair.

$$\begin{aligned} y_{ijk} &\sim \text{Normal}(\bar{f}_{ijk}, \sigma_{\text{mat}}(i, j)) \\ \bar{f}_{ijk} &= -\lambda_{ij} t \end{aligned} \tag{65}$$

We placed a diffuse Normal prior on the log half-lives $\eta_{ij} = \frac{\log_{10}(2)}{\lambda_{ij}}$ and a Half-Normal prior on the standard deviations $\sigma_{\text{mat}}(i, j)$:

$$\begin{aligned} \ln(\eta_{ij}) &\sim \text{Normal}(-2, 4) \\ \sigma_{\text{mat}}(i, j) &\sim \text{Half-Normal}(0.6, 0.2) \end{aligned} \tag{66}$$

##### 6.2.2 Estimation model predictive checks

We assessed appropriateness of priors with prior predictive checks (Fig. A30) and goodness-of-fit with posterior predictive checks (Fig. A31). Prior checks suggested that prior distributions were agnostic over the parameter values of interest, and posterior checks suggested a good fit of the model to the data.

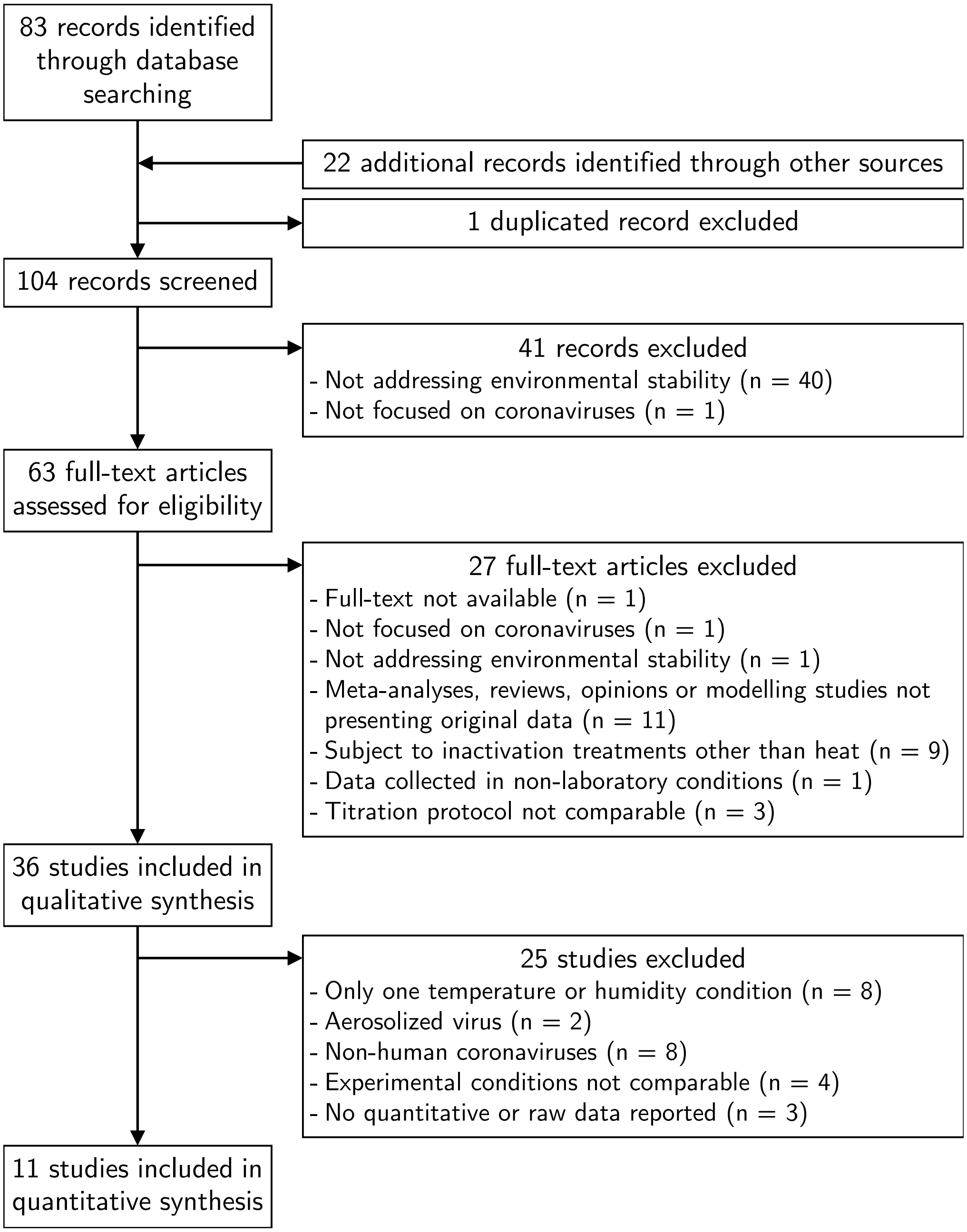

**Figure A29. Selection process of the studies included in the meta-analysis of the effect of temperature and humidity on human coronaviruses.**

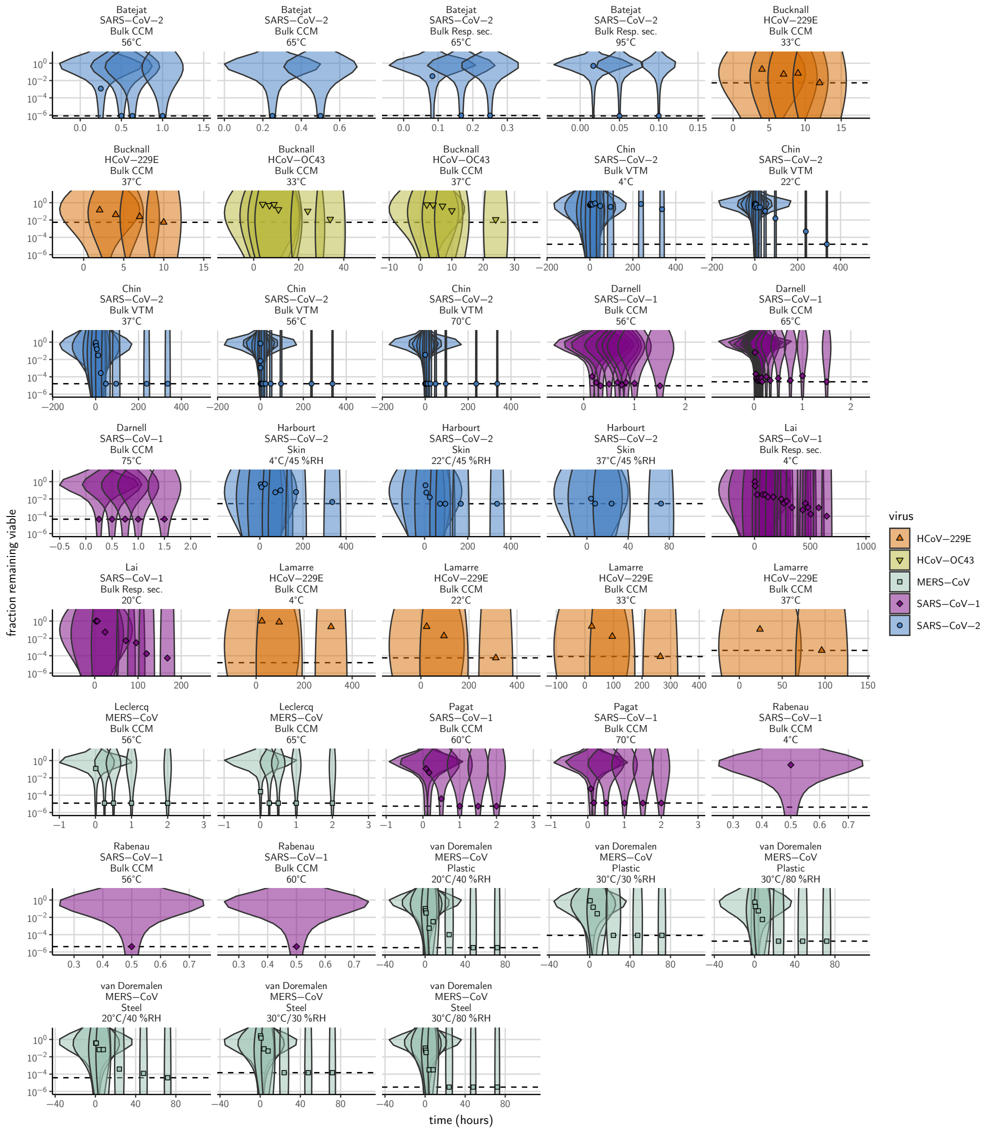

**Figure A30. Prior predictive check for empirical coronavirus decay from literature data.** Violin plots show distribution of simulated titers sampled from the prior predictive distribution. Points show estimated titers for each collected sample based on data extracted from the literature. Shape and color indicates virus. x-axis shows time since first available measure. Study author, virus, and experimental conditions—material, temperature, and relative humidity (RH)—indicated at the top of each panel. Black dotted line shows LOD for each experiment. Wide coverage of violins relative to datapoints shows that priors are agnostic over the titer values of interest, and that the priors regard both fast and slow decay rates as possible.

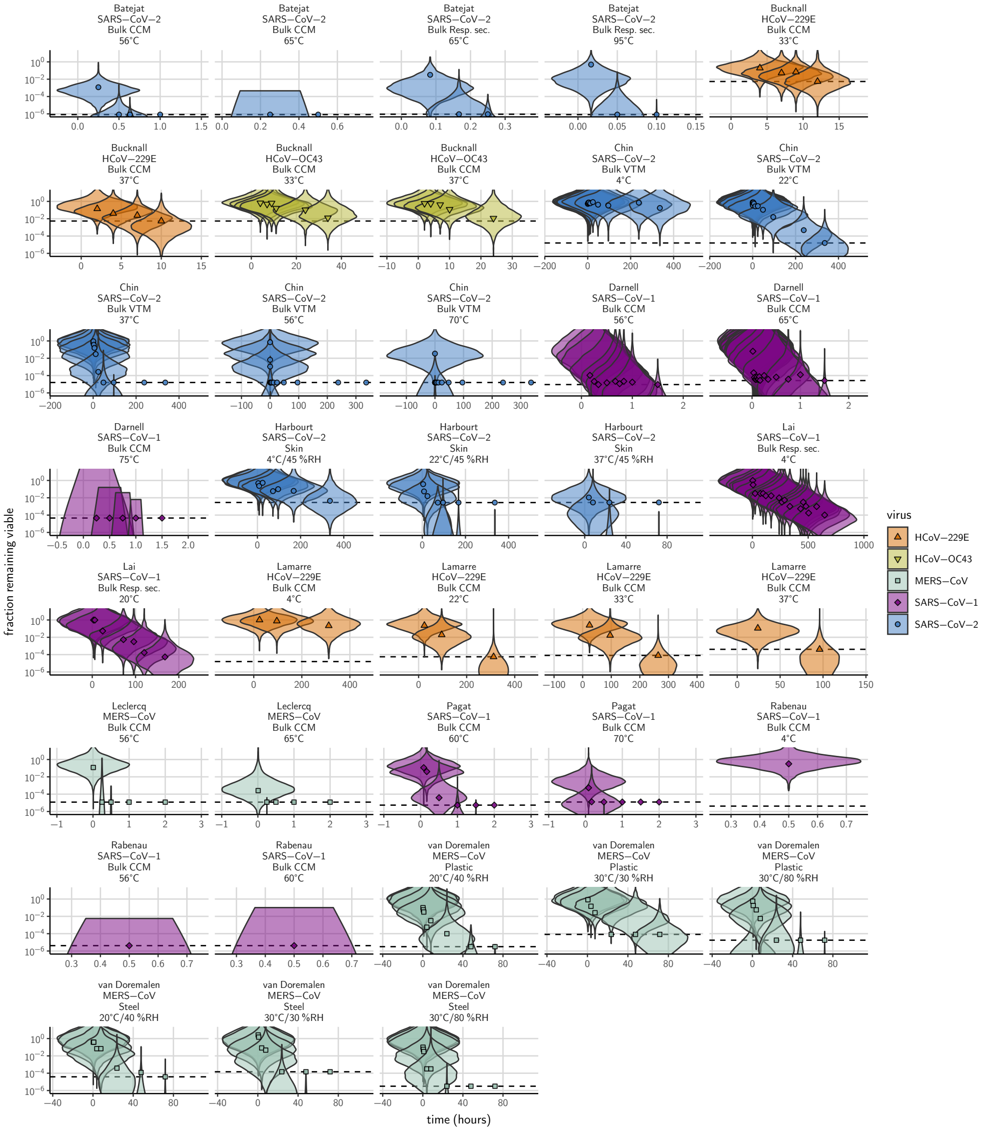

**Figure A31. Posterior predictive check for empirical coronavirus decay from literature data.** Violin plots show distribution of simulated titers sampled from the posterior predictive distribution. Points show estimated titers for each collected sample based on data extracted from the literature. Shape and color indicates virus. x-axis shows time since first available measure. Study author, virus, and experimental conditions—material, temperature, and relative humidity (RH)—indicated at the top of each panel. Black dotted line shows LOD for each experiment. Tight correspondence between distribution of posterior simulated titers and independently estimated titers suggests the model fits the data well.

#### 6.3 SARS-CoV-1 and MERS-CoV estimates

As noted in the Main Text Methods, we made half-life estimates for SARS-CoV-1 and MERS-CoV at 22 °C and 40 % RH during the evaporation and quasi-equilibrium phases using data collected by our group during previous studies [15]. We included these estimates in the meta-analysis alongside the estimates described above. Table A5 shows the estimated half-lives for these data, and Fig. A32 shows the fit of the simple regression model to these data.

**Table A5. Estimated half-lives in hours of SARS-CoV-1 and MERS-CoV on polypropylene as a function of temperature (T) and relative humidity (RH).** Estimated half-lives are reported as posterior median and the middle 95% credible interval.

| | T (°C) | RH (%) | virus | median half-life (h) | 2.5 % | 97.5 % |
|---|---|---|---|---|---|---|
| quasi-equilibrium phase | 22 | 40 | SARS-CoV-1 | 6.42 | 5.22 | 7.92 |
| | 22 | 40 | MERS-CoV | 3.16 | 2.53 | 3.97 |
| evaporation phase | 22 | | SARS-CoV-1 | 11.55 | 1.43 | 207.68 |
| | 22 | | MERS-CoV | 13.18 | 1.09 | 217.34 |

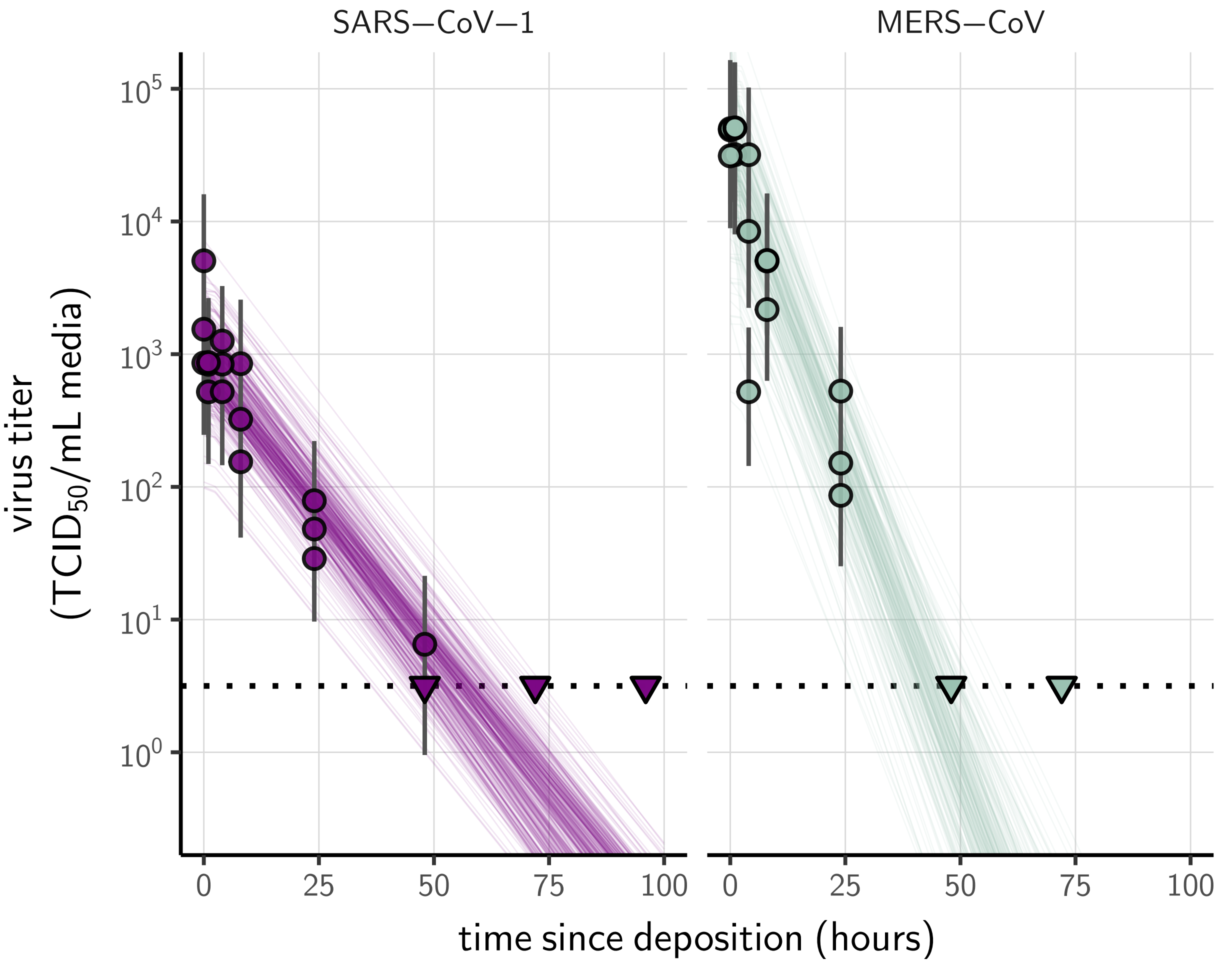

**Figure A32. Fit of simple regression model to SARS-CoV-1 and MERS-CoV data.** Points show posterior median estimated titers in $\log_{10}\text{TCID}_{50}/\text{mL}$ for each sample; lines show 95 % credible intervals. Time-points with no positive wells for any replicate are plotted as triangles at the approximate single-replicate limit of detection (LOD) of the assay—denoted by a black dotted line at $10^{0.5}$ $\text{TCID}_{50}/\text{mL}$ media—to indicate that a range of sub-LOD values are plausible. Three samples collected at each time-point. x-axis shows time since sample deposition. Lines are random draws (10 per sample) from the joint posterior distribution of the initial sample virus concentration and the estimated decay rate; the distribution of lines gives an estimate of the uncertainty in the decay rate and the variability of the initial titer for each experiment.

#### 6.4 Prediction of half-lives

##### 6.4.1 Absolute predictions

Where both temperature and humidity were available for a measurement from the literature, we were able to predict the absolute half-life directly from our modeled concentration fit, as parametrized from our own SARS-CoV-2 data. These predictions are plotted in Main Text Fig. 3c and Fig. A7.

##### 6.4.2 Relative predictions

For many studies, however, only temperature information was available. Moreover, heterogeneities both among viruses and among laboratory protocols could shift the half-live by a constant factor relative to our SARS-CoV-2-polypropylene-DMEM data. To account for this, we made within-study relative predictions for studies with at least two temperature and/or humidity conditions on the same side of the ERH for a given virus on a given surface. For each such set of experiments, we chose the experiment whose temperature was closest to 20 °C to serve as the reference experiment. If there were multiple such experiments, we picked the experiment with the relative humidity closest to the ERH.

Our mechanistic model implies that the ratio of a pair of half-lives $\eta_1$ and $\eta_2$ at ambient temperatures $T_1$ and $T_2$ and super-ERH relative humidities $h_1$ and $h_2$ is given by:

$$\frac{\eta_1}{\eta_2} = \left(\frac{\ln(h_2)}{\ln(h_1)}\right)^{\frac{1}{\alpha_c}} \exp\left[\frac{E_a}{R}\left(\frac{1}{T_1} - \frac{1}{T_2}\right)\right] \tag{67}$$

If $h_1$ and $h_2$ are both sub-ERH, we have:

$$\frac{\eta_1}{\eta_2} = \exp\left[\frac{E_a}{R}\left(\frac{1}{T_1} - \frac{1}{T_2}\right)\right] \tag{68}$$

Where no information about ambient relative humidity was available, we assumed humidities were shared across experiments and were super-ERH, and therefore used equation 67 with $h_1 = h_2$ to make predictions. Note that these predictions are independent of $\alpha_s$ and $A$; they rely only on relative rates of inactivation, not absolute ones. These relative predictions according to equations 67 and 68 are plotted in Fig. 3d.

#### 6.5 Discussion of the results

We report half-life estimates for each experimental condition in Table A2. This meta-analysis highlights the same qualitative effect of temperature as our data: higher temperatures are associated with faster virus decay (shorter half-lives), with SARS-CoV-2 half-life in bulk medium varying from several hours at 4 °C to less than 15 s at 95 °C. The direct comparison of coronavirus half-lives across humidities is difficult, as only a few studies measured virus decay at several humidities with a fixed temperature.

This data set includes data collected following heterogeneous experimental procedures, which can considerably impact virus inactivation kinetics. For instance, we included data collected from suspensions at different pH, which notably explains the difference between the half-lives estimated from Bucknall et al. 1972 [70] (cell culture medium at pH 7.4) and Lamarre et al. 1989 [77] (cell culture medium supplemented to reach pH 6) for HCoV-229E in bulk medium at 33 °C and 37 °C. Indeed, Lamarre et al. 1989 [77] showed that pH 6 is optimal for HCoV-229E stability, hence the higher half-lives reported by this study. We also included data collected from suspensions supplemented with varying levels of proteins (from 1 %[80] to 10 %[71, 74] of fetal calf serum) although protein concentration is known to impact virus inactivation kinetics [63, 81]. Containers used to expose samples to environmental conditions can also impact virus inactivation rate, but this information is rarely reported [21]. Notably, the two SARS-CoV-2 points in Main Text Fig. 3d that show shorter-than-predicted half-lives are from heated bulk medium in closed vials, where inactivation is known to be rapid [21].

Despite this heterogeneity of the data collection process, and the high uncertainty of some half-life estimates, we find good qualitative agreement between model predictions and model-free estimates (see Main Text, Fig. 3, and Fig. A7).

### 7 Methodological implications for experimental studies on virus stability

The characterization of the mechanisms by which humidity impacts virus stability allows us to draw methodological implications for future experimental studies. First, since solute concentration plays a critical role in the decay of viable virus, studies interested in virus viability should either include a measure of solute concentration over time (ideally via medium evaporation or precise measurements of sample mass through time), or focus on the quasi-equilibrium phase (during which solute concentration can be assumed to be constant). Second, since the evaporative kinetics and the resultant solute environments depend on the composition of the initial suspension medium, quantitative estimates of duration of virus viability based on experiments conducted in different media should be compared with caution. In our meta-analysis, we were able to make accurate relative predictions of data from multiple artificial medium formulations as well as from bodily fluids; this suggests that the underlying mechanisms are robust to variation in suspension medium, though absolute durations may vary. Third, given the non-linear relationship between virus half-life and relative humidity, studies interested in the effect of humidity on virus viability should include a wide range of conditions at constant temperature, including both sub- and super-ERH conditions.

Code for titer estimation and model fitting is freely available the online data and code repository, and could readily be adapted to the study of other viruses.

### References

[67] Christophe Batéjat, Quentin Grassin, and Jean-Claude Manuguerra. "Heat inactivation of the Severe Acute Respiratory Syndrome Coronavirus 2". In: *bioRxiv* (2020). DOI: 10.1101/2020.05.01.067769.

[68] Mike J Blandamer et al. "Activity of water in aqueous systems; a frequently neglected property". In: *Chemical Society Reviews* 34.5 (2005), pp. 440–458.

[69] C Brownie et al. "Estimating viral titres in solutions with low viral loads". In: *Biologicals* 39.4 (2011), pp. 224–230.

[70] Robert A Bucknall et al. "Studies with human coronaviruses II. Some properties of strains 229E and OC43". In: *Proceedings of the Society for Experimental Biology and Medicine* 139.3 (1972), pp. 722–727. DOI: 10.3181/00379727-139-36224.

[71] Miriam ER Darnell et al. "Inactivation of the coronavirus that induces severe acute respiratory syndrome, SARS-CoV". In: *Journal of virological methods* 121.1 (2004), pp. 85–91. DOI: 10.1016/j.jviromet.2004.06.006.

[72] *Dulbecco's Modified Eagle's Medium (DME) Formulation*. URL: https://www.sigmaaldrich.com/life-science/cell-culture/learning-center/media-formulations/dme.html (visited on 09/03/2020).

[73] Andrew Gelman et al. *Bayesian Data Analysis, Third Edition*. CRC Press, Nov. 1, 2013.

[74] David E. Harbourt et al. "Modeling the Stability of Severe Acute Respiratory Syndrome Coronavirus 2 (SARS-CoV-2) on Skin, Currency, and Clothing". In: *PLoS Neglected Tropical Diseases* 14.11 (Nov. 2020), pp. 1–8. DOI: 10.1371/journal.pntd.0008831.

[75] Alex Kale, Matthew Kay, and Jessica Hullman. "Visual reasoning strategies for effect size judgments and decisions". In: *IEEE Transactions on Visualization and Computer Graphics* (2020).

[76] Mary YY Lai, Peter KC Cheng, and Wilina WL Lim. "Survival of Severe Acute Respiratory Syndrome coronavirus". In: *Clinical Infectious Diseases* 41.7 (2005), e67–e71. DOI: 10.1086/433186.

[77] Alain Lamarre and Pierre J Talbot. "Effect of pH and temperature on the infectivity of human coronavirus 229E". In: *Canadian journal of microbiology* 35.10 (1989), pp. 972–974. DOI: 10.1139/m89-160.

[78] India Leclercq et al. "Heat inactivation of the Middle East Respiratory Syndrome coronavirus". In: *Influenza and other respiratory viruses* 8.5 (2014), pp. 585–586. DOI: 10.1111/irv.12261.

[79] David Moher et al. “Preferred Reporting Items for Systematic Reviews and Meta-Analyses: The PRISMA Statement”. In: *PLoS Medicine* 6 (2009), p. 6.
[80] Anne-Marie Pagat et al. “Evaluation of SARS-Coronavirus decontamination procedures”. In: *Applied Biosafety* 12.2 (2007), pp. 100–108. DOI: 10.1177/153567600701200206.
[81] Boris Pastorino et al. “Prolonged Infectivity of SARS-CoV-2 in Fomites”. In: *Emerging Infectious Diseases* 26.9 (2020), pp. 2256–2257. DOI: 10.3201/eid2609.201788.
[82] HF Rabenau et al. “Stability and inactivation of SARS coronavirus”. In: *Medical microbiology and immunology* 194.1-2 (2005), pp. 1–6. DOI: 10.1007/s00430-004-0219-0.
[83] John Redrow et al. “Modeling the evaporation and dispersion of airborne sputum droplets expelled from a human cough”. In: *Building and Environment* 46.10 (2011), pp. 2042–2051.
[84] Ankit Rohatgi. *WebPlotDigitizer*. 2019. URL: https://automeris.io/WebPlotDigitizer.
[85] Caroline ER Rowell and Hana M Dobrovolny. “Energy Requirements for Loss of Viral Infectivity”. In: *Food and environmental virology* (2020), pp. 1–14.
